# A framework for quantifying deviations from dynamic equilibrium theory

**DOI:** 10.1101/755645

**Authors:** Michael Kalyuzhny, Curtis H. Flather, Nadav M. Shnerb, Ronen Kadmon

## Abstract

Community assembly is governed by colonization and extinction processes, and the simplest model describing it is Dynamic Equilibrium (DE) theory, which assumes that communities are shaped solely by stochastic colonization and extinction events. Despite its potential to serve as a null model for community dynamics, there is currently no accepted methodology for measuring deviations from the theory and testing it. Here we propose a novel and easily applicable methodology for quantifying deviations from the predictions and assumptions of DE by comparing observed community time-series to a randomization-based null model. We show that this methodology has good statistical properties on simulated data, and it can detect deviations from both the assumptions and predictions of DE in the classical Florida Keys experiment. We discuss alternative methods and present guidelines for practical use of the methodology, hoping it will enhance the applicability of DE as a reference for studying changes in ecological communities.

## Introduction

The dynamics of ecological communities reflect the process that shape them. While understanding these processes has been a primary focus of ecological research for decades (Hutchinson 1961; MacArthur & Wilson 1967; Chesson 2000) most studies examine how such processes shape static patterns (e.g. the dependence of species diversity on area and isolation, MacArthur & Wilson 1963; Johnson 1975; Kadmon & Allouche 2007). In contrast, in recent years there is a growing interest in understanding temporal patterns of community dynamics and change (Vellend *et al.* 2013; Dornelas *et al.* 2014; Kalyuzhny *et al.* 2014a; Kalyuzhny *et al.* 2015; Magurran *et al.* 2015). It is increasingly understood that temporal patterns hold much valuable information on how ecological communities are assembled (Chisholm *et al.* 2014), and on their responses to anthropogenic effects (Elahi *et al.* 2015; Magurran *et al.* 2015).

Despite its promise, the analysis of empirically observed community dynamics proves quite complex (Dornelas *et al.* 2013). Community time series (where the abundance or presence of a set of species is recorded over time) are inherently high-dimensional, leading to questions on the proper way to examine them and cope with this dimensionality. They are also frequently quite short, temporally autocorrelated, noisy and prone to errors. While the latter is true for most ecological data, since the aim in time-series analysis is often estimating changes or variances, ignoring errors or mistreating them inherently introduces *biases* (on top of the errors), sometimes quite severe (Knape & de Valpine 2012; Kalyuzhny *et al.* 2014b). Even if these issues are dealt with, interpreting patterns of community dynamics is very difficult without having some reference (Magurran 2016). This is because many ecological theories predict that communities may undergo constant changes (e.g. MacArthur & Wilson 1967; May 1976; Hubbell 2001; Tilman 2004; Kessler & Shnerb 2015), so the naïve null hypothesis of ‘zero difference’, so common in traditional statistics, would lead to erroneous conclusions. Hence, the observed dynamics should be compared to the expected dynamics under some model. Nevertheless, with the difficulty of using a dynamic model as reference comes opportunity: support for a dynamic model provides evidence for the processes that it incorporates as assumptions, while rejecting it could point out to an important role of other processes.

The simplest dynamic model of ecological communities, and one of the most classic, is the Dynamic Equilibrium model (DE, MacArthur & Wilson 1963; MacArthur & Wilson 1967). The most applicable version of this model, which we consider here, is the discrete-time version of Simberloff’s species-level formulation (Simberloff 1969). The model considers a local community (island) that receives immigrants from a regional pool (mainland) with *Sreg* species. For every species *i* among them, if the species was absent at time *t* it will colonize the community by time *t* + 1 and become present with probability *C*_*i*_, while if it was present it will go extinct and become absent by time *t* + 1 with probability *E*_*i*_. Therefore, the assumptions of DE can be summarized as follows: 1) the processes of colonization and extinction occur at fixed rates; 2) the dynamics of species are independent of each other. These colonization and extinction probabilities may depend on properties of the environment and of the species. Based on these assumptions, the model predicts that species richness would fluctuate around an equilibrium and show limited changes, while composition would undergo some turnover.

The DE model has had remarkable success in explaining static patterns such as species-area and species-isolation relationships in multiple systems (MacArthur & Wilson 1967; Lomolino *et al.* 2006). However, its dynamic predictions have not often been tested. Several classical works supported the qualitative predictions of DE (Diamond 1969; Simberloff & Wilson 1969), while others brought contrasting evidence (Abbott & Grant 1976; Price 1976). A considerable part of the controversy was related to the difficulty of rigorously quantifying to what degree do communities deviate from the predictions of the theory, and what magnitude of changes would be considered excessive, thereby rejecting its predictions (Simberloff 1983; Schoener 2010). Recently, there is increased interest in studying observed changes and turnover in communities (e.g. Vellend *et al.* 2013; Dornelas *et al.* 2014; Magurran *et al.* 2015), motivated by concerns over how they are impacted by anthropogenic effects. A recent study has used DE as a null model for the magnitude of compositional turnover in ecological communities worldwide, finding that for 100 communities, turnover is typically much higher than expected under DE (Dornelas *et al.* 2014).

Previous tests of DE were either qualitative (Diamond 1969) or used ad-hoc quantitative methods whose properties have not been adequately studied. Furthermore, most previous studies have only tested the *predictions* of DE (mainly regarding richness and species composition), while the *assumptions* of the theory remained mostly untested. This is unfortunate, because any deviation from the predictions of a model must stem from violating some assumptions. To fulfill the potential of DE to inform research on community dynamics, there is a need for a methodology that would allow quantifying deviations from both the predictions and assumptions of DE and the statistical significance of such deviations.

We believe that such a methodology should satisfy the following requirements:

1. For data generated by DE, the quantified effect size should have a predetermined expected value (e.g. 0).
2. For such data, the type I error rate of the statistical test for the significance of the effect should equal the preset level (we used α = 0.05 for all analyses).
3. For data not generated by DE, the methodology should have good power to detect and quantify the deviation(s).
4. Since empirical community dynamics may show deviations from DE only in some respects, the methodology should be able to quantify such different facets.
5. The methodology should be robust to issues with the data such as false or incomplete detection of species and missing years.
6. The methodology should be easily applicable.

Here we propose such a methodology for testing both the assumptions and the predictions of DE by comparing a set of statistics to a novel randomization-based null model. We begin by describing the two elements of the methodology, namely the proposed Presence-Absence Randomization wIthin periodS (PARIS) null model and five proposed statistics for testing the assumptions and predictions of the model. We move forward to exploring the properties of the methodology, in particular its type I error rate, robustness and power. We then apply the methodology to the most classical dataset where DE has been tested – the defaunation experiment at Florida Keys (Simberloff & Wilson 1969). We conclude with discussing the interpretation of outputs of the methodology and with an extensive and quantitative comparison of our methodology with several previously proposed methods.

## The methodological framework

Our approach for testing DE consists of two components, which are, in principle, independent (i.e. each can be altered separately) – a null model for generating synthetic community time series, and statistics that characterize various dynamical aspects and are compared to the null model. The basic assumption is that samples of the community are taken in discrete regular time-interval (‘years’ without loss of generality), and some of the samples may be missing.

### The PARIS null model

The aim of the null model is generating synthetic time series of community composition that resemble the original time-series, but adhere to the assumptions of DE, namely, species independence and uniform colonization and extinction rates. This is done by the following algorithm, that is applied to single-species time series of presence-absence through time (Figure. 1a).

**Figure 1.**
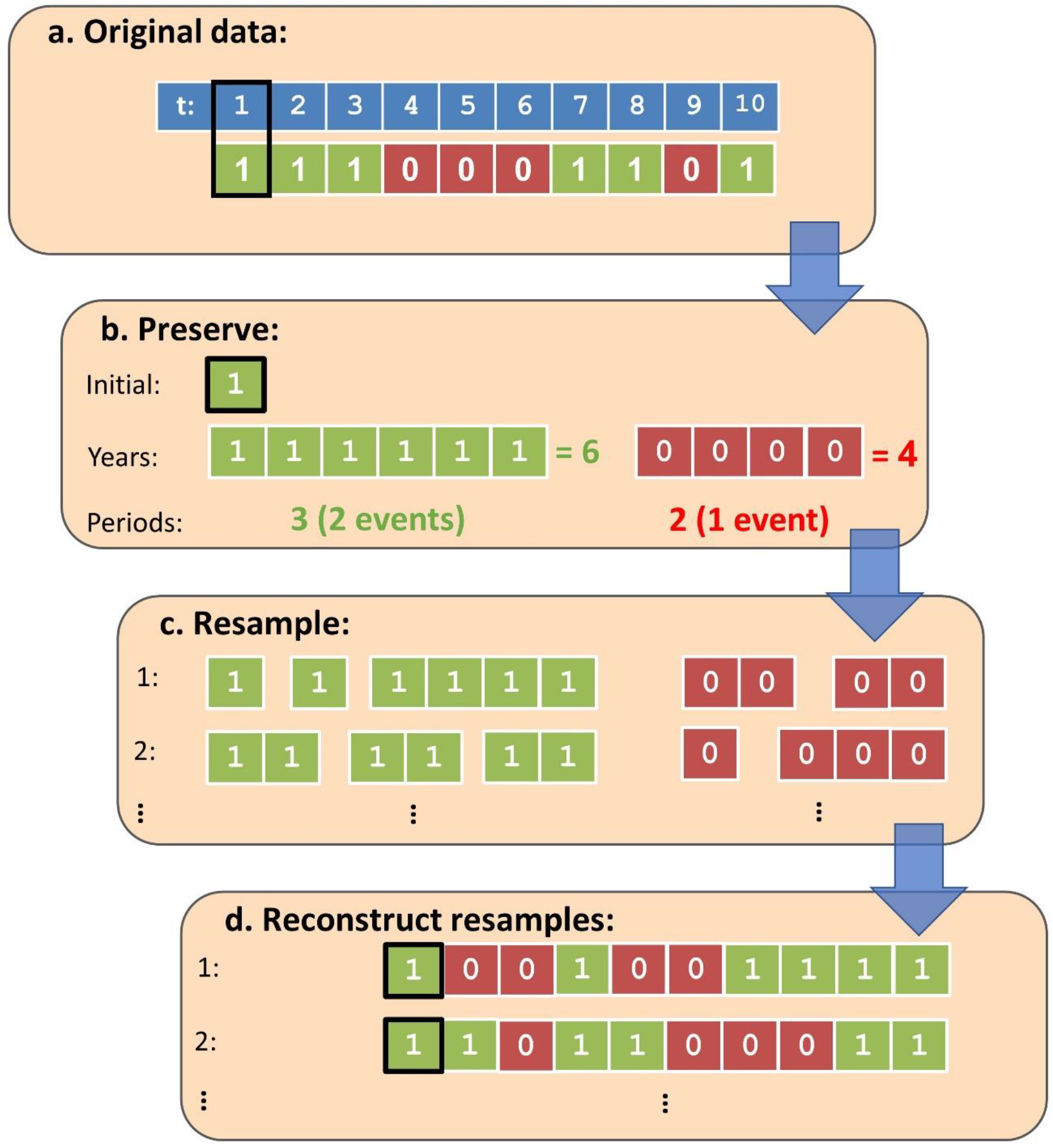
Example application of the PARIS null model to an observed time-series. (a) The algorithm gets as input a single species time-series of presence (‘1’) and absence (‘0’). (b) The algorithm preserves the initial state (purple box) the number of years of presence and absence and the number of periods they are divided into. This is done separately for presence and absence. Next (c), the presence and absence periods are resampled by dividing the total time of presence and absence at *s* – 1 random time points (extinction and colonization events, respectively). Finally (d), the time series is reconstructed by adding the presence and absence periods consequently, beginning with the initial state.

The PARIS algorithm preserves for each species the initial state (presence or absence), the total number of years of presence and the total number of years of absence, the number of colonization and extinction events as well as the total number of periods of presence and periods of absence (Figure 1b). The only thing that can change is the duration of these periods. These requirements imply that the final state is also preserved. Consider there were *y* years of presence in the empirical time series, separated into *s* periods by extinction events (that initiate periods of absence). The algorithm chooses *s* – 1 out of *y* – 1 years without replacement, simultaneously. Then, s – 1 events are assigned to happen after the chosen years (Figure 1c). The analogous procedure is applied for periods of absence being separated by colonization events. Please note that the number of events that need to be assigned may be lower than the total number of events, e.g. the absence years in Figure 1. If there was only one period of presence or absence, it will remain as in the original data in all resamples. Missing years are removed before the algorithm is applied and then are put back at their original timing (the implications of this are tested below). After resampling the duration of all presence and absence events separately, the algorithm combines them, beginning with the initial state (Figure 1d).

This is exemplified in figure 1, where the original time series is [1 1 1 0 0 0 1 1 0 1], where ‘1’ stands for a year of presence and ‘0’ stands for a year of absence. There are six years of presence within three periods and four years of absence within two periods, so two extinction events and one colonization event should be assigned. For example, choosing extinction to occur after the 1st and 2nd years of presence and colonization to occur after the 2nd year of absence will lead to the following synthetic time series: [1 0 0 1 0 0 1 1 1 1], as in the upper example in Figure 1. All such possible resamples are equiprobable.

This algorithm generates independent species time-series since it is applied to different species independently. Moreover, it approximately generates events that are characteristic of a process with uniform rates. This is because the basic assumption of a Poisson process (the continuous time analogue of our colonization and extinction processes) is that events occur independently and with equal probability at any time. As a result, given that an event has occurred within a time interval, it can occur at any time with equal probability, and the same is true given several events have occurred (Gallager 2012).

Hence, The PARIS model satisfies the assumptions of DE and can be used to generate multiple synthetic time series that ‘force’ these assumptions on the data. It is important to note that this model preserves the initial state of the community, as well as the final state (the presence and absence of each species in the first and final year in the data). Preserving the initial state is important, because dynamics generally depends on initial state. Moreover, the approach does not assume that the community is at steady state, making it robust to violations of this assumption.

### Summary statistics

Data describing the species composition of a community through time are inherently high-dimensional and can therefore be summarized in multiple ways. The statistics we propose quantify core aspects of the dynamics relative to the expectations of DE. The significance of their deviation from DE can be calculated by comparing their observed value to the distribution of expected values under the null model. We define five statistics: two statistics refer to the predictions (changes in richness and composition) and three statistics refer to the assumptions (species independence and uniformity of colonization and extinction rates). All statistics are constructed so that their mean if DE is true is zero. Other statistics can also be defined and tested in the same manner within the framework.

The most basic prediction of DE is that species richness would be relatively stable through time at steady state. To quantify the magnitude of changes in richness, we use the statistic *ds* (for parameter *k*, more on this later), which quantifies the magnitude of temporal changes in species richness, beyond the expectation of DE (Figure 2a):

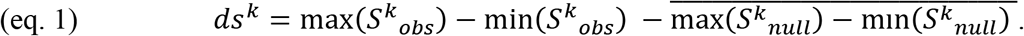

**Figure 2.**
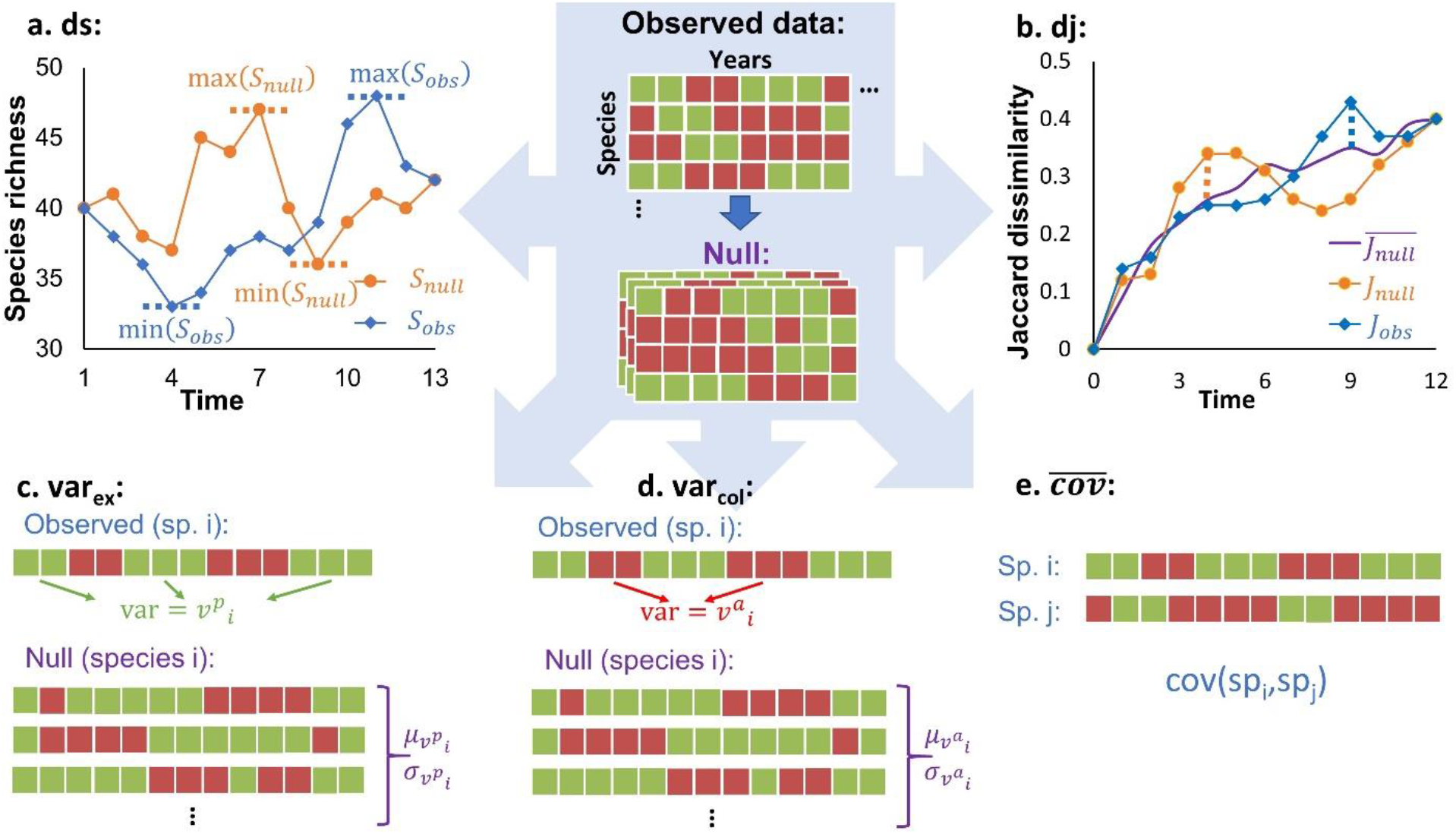
Computing the five proposed statistics, which are based on the observed and null community time-series of presence (green) and absence (red). Only one or a few resampled null communities are illustrated. In (a) excessive richness changes (*ds*) are computed by comparing the maximal richness change in the observed (blue) to the null (orange) time-series. In (b) excessive compositional changes are computed by comparing the maximal upward deviations (dashed lines) of: 1) the observed Jaccard curve (blue) from the expected curve under the null (purple), and 2) a single resampled community curve (orange) from the expected. In (c-d) the rate uniformity of extinction and colonization statistics (*var*_*ex*_ and *var*_*col*_) are computed using the variance of presence (*V*^*p*^_*i*_) and absence (*V*^*a*^_*i*_) period durations, standardized using the means and variances computed using the null. This computation is shown for only a single species, so these standardized measures should also be averaged over species. Finally, (e) presents the temporal covariance of a pair of species, and this should be averaged over all pairs to get 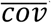.

In other words, for the observed time series of species richness, *S*_*obs*_, we compute the change between the maximal richness and the minimal richness observed. The same change is then computed for every synthetic time-series generated by the null model, *S*_*null*_. Finally, the expected change under the null is subtracted from the observed change.

Under the PARIS null model, richness in the first and last year in every randomization precisely equals the observed values, which can lower power. For this reason, it is useful to perform the calculation of *ds* for the ‘central’ part of the time series, after ‘cutting’ a proportion *k* of the years on both sides in the empirical data as well as in the randomizations. This is the meaning of the superscript ‘k’ in eq. 1. Positive *ds* indicates that the temporal changes in species richness are larger than expected, while negative *ds* indicates smaller changes. The scale at which these changes are larger is not defined, so a large *ds* can result from any combination of long-term trends and short-term variation.

The statistic *dj* quantifies the magnitude of temporal changes in composition, beyond the expectation of DE (Figure 2b):

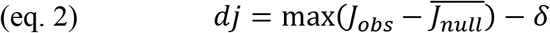

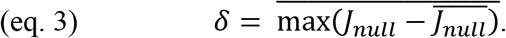

J is a curve describing the Jaccard dissimilarity of each year w.r.t the first year. *J*_*obs*_ is this curve for the observed data, while *J*_*null*_ is computed for a single realization of the null model. Hence, eq. 2 computes the difference between *J*_*obs*_ and the expected *J*_*null*_ in the year when this difference is maximal. Since some deviations from the expected *J*_*null*_ will occur even under DE, we subtract the quantity *δ*, to ensure that *dj* will have a mean of zero under DE. *δ* is the expected maximal deviation of a stochastic realization of the null model *J*_*null*_ from the expected *J*_*null*_. In analogy to *ds*, *dj* quantifies excess compositional changes without specifying the temporal scales at which they take place.

To test the assumptions of uniformity in colonization and extinction rates, we use the statistics *var_col_* and *var_ex_*, respectively (Figure 2c-d). These statistics are based on comparing the variance in the duration of absence and presence periods to the expectation under the null. For each species with two or more periods of presence or absence, we compute the variance in their durations (separately for presence and absence) and then standardize it using the null model.

These standardized variances are then averaged over species:

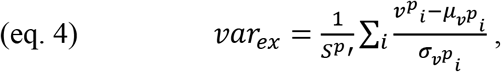

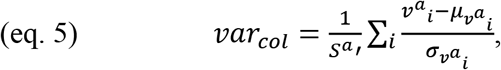

Where *V*^*p*^_*i*_ and *V*^*a*^_*i*_ are the *observed* variances in the durations of presence and absence for species i, respectively; *μ*_*v*_*p*_*i*_ and *μ*_*v*_*a*_*i*_ are the *expected* variances in the durations of presence and absence for species i, respectively; and *σ*_*v*_*p*_*i*_ and *σ*_*v*_*a*_*i*_ are the SD of the variances in the durations of presence and absence for species i, respectively, where the expectation and SD are calculated over resamples. These values are averaged over all species that have at least 2 events of presence or absence, respectively, *S*^*p’*^ and *S*^*a’*^.

Positive values of *var_col_* and *var_ex_* indicate that the variation in the duration of absence and presence periods, respectively, is higher than expected under a stochastic process with uniform rates. Negative values would indicate that the variation is too small (e.g. hypothetically the species is always observed for five years and then it disappears).

To test species independence, we compute the covariance between every pair of species time-series and average over the pairs to get the average covariance statistic (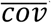, Figure 2e). Positive covariance indicates that species appear and disappear together in the data, on average, while negative covariance indicates that species replace each other.

All these statistics are expected to have a mean of zero under DE. A small exception is average covariance, which may have a positive value if the community initially is far from equilibrium. For all statistics except *dj*, a two-sided test is performed by comparing the observed statistic to a distribution of the statistic in multiple randomizations and computing the p values. For *dj* the test is one sided.

If data for several similar local communities are available within some dataset, it may be of interest to calculate the average statistics and their significance at the dataset level. This can be done by recording the randomization-produced statistics for each community separately. Then, the test is performed by comparing the observed average statistic to the distribution of averaged (over communities) randomized statistics. For example, if a dataset consists of 10 communities, we first compute the average *dj* (over the communities). We then compute and record 2·10^4^ randomization-generated *dj*s for each of the communities. Now we average the randomized statistics over the communities, to get 2·10^4^ average *dj*s, that will serve as the null distribution for the test.

### Performance of the methodology

We test three aspects of the methodology: First, we consider the behavior of all statistical tests under DE, asking: is the type I error rate excessive? and are effect sizes roughly zero? Next, we ask whether these properties are robust to issues with the data, specifically false detection, incomplete detection and missing years. Finally, we examine the ability of the methodology to detect and quantify deviations under an alternative model. Based on the results in this section we provide recommendations on the application of the methodology.

We examined the methodology using simulations representing multiple scenarios. Briefly, we considered all combinations of two levels of mean colonization rates and two levels of mean extinction rates (‘high’ and ‘low’), zero correlation between colonization and extinction or a negative correlation (i.e. species with high extinction rates have low colonization rates), and three levels of regional species pool sizes (S_reg_): 30, 90 and 270 species. For each of the 24 scenarios above we generated 5000 synthetic communities and calculated the five statistics and their P values using 500 resamples (of each community). To test the robustness of the methodology, we introduced errors into the data: For every presence record in every community, there was a probability *P*_*miss*_ to falsely miss the species (incomplete detection). Alternatively, we introduced a probability *P*_*false*_ to falsely detect a species that is not observed in the data. Finally, we transformed a given number of years into missing years. The effect size and P value were recorded at different levels of these factors. Finally, we tested the behavior of the methodology under an alternative community dynamics model, the State-Dependent Extinction (SDE) model. This model assumes that extinction rates fluctuate through time, and species may synchronously respond to these fluctuations. Full details of the testing procedure are presented in appendix S1.

An initial examination revealed that calculating *ds* for the ‘central’ part of the time series has similar type I error rates and more power than k = 0, so we used k = 0.06 throughout this work.

We found that under the multiple scenarios we considered, type I error rates for all the summary statistics were mostly very close to 0.05 (Figure 3). The effect size was also very close to zero (Figure S3). Introducing issues with the data (Figures S4-S9, Table 1) introduced some errors into the inference, but many of the statistics were quite robust. Most noticeably, *var_col_* and *var_ex_* were very sensitive to incomplete and false detection, consequently. These errors introduce very short periods (typically a year) of absence and presence, therefore strongly inflating the variances in the duration of absence and presence periods. Moreover, *dj* was sensitive to issues with the data, mostly if regional richness is high (hundreds of species) and the issues are severe (many missing years, many short presence or absence periods due to errors). In some cases, errors also generated negative values of *ds* and *dj* but this did not inflate type I error rates. Nevertheless, in most of the conditions we considered, the methodology was quite robust, and Table 1 can be used as a reference for considering the possible effect of issues with the data when interpreting results.

**Table 1.** Sensitivity of the five statistics with the PARIS null model to issues with the data. Each cell represents the worst-case scenario (over eight settings for the colonization and extinction rate) for different levels of regional richness (S_reg_) and different levels of missing years, incomplete detection probability (*P*_*miss*_) and false detection probability (*P*_*false*_). The color represents the probability of type I error (which ideally should be ~ 0.05): green: P < 0.075; yellow: 0.075 ≤ P < 0.1; red: P ≥ 0.1. The symbol represents the resulting standardized effect size (SES, which ideally should be ~ 0): no symbol: SES < 0.1; **X** : 0.1 ≤ SES < 0.2; **XX**: SES ≥ 0.2. The symbol (−) represents a negative SES.

| Statistic | $S_{reg}$ | Missing years (number of years): | | | | Incomplete detection ($P_{miss}$): | | | False detection ($P_{false}$): | | |
| --- | --- | --- | --- | --- | --- | --- | --- | --- | --- | --- | --- |
|  |  | 1 | 2 | 4 | 8 | .1 | .4 | .7 | .01 | .03 | .1 |
| dj | 30 |  |  |  |  |  |  |  |  | X(-) |  |
|  | 90 |  |  |  | XX | X |  |  | X(-) | X(-) | XX |
|  | 270 |  | X | XX | XX | XX | XX |  | X(-) | XX | XX |
| ds | 30 |  |  |  |  |  |  |  |  |  | X(-) |
|  | 90 |  |  |  |  |  |  |  |  | X(-) | X(-) |
|  | 270 |  |  |  |  | X(-) |  |  |  | X(-) |  |
| var<sub>col</sub> | 30 |  |  |  |  | XX | XX | XX |  |  |  |
|  | 90 |  |  |  |  | XX | XX | XX |  |  |  |
|  | 270 |  |  |  |  | XX | XX | XX |  |  |  |
| var<sub>ex</sub> | 30 |  |  |  |  |  |  |  | XX | XX | XX |
|  | 90 |  |  |  |  |  |  |  | XX | XX | XX |
|  | 270 |  |  |  |  |  |  |  | XX | XX | XX |
| $\overline{cov}$ | 30 | | | | | | | | | | |
|  | 90 |  |  |  |  |  |  |  |  |  |  |
|  | 270 |  |  |  |  |  |  |  |  |  |  |

**Figure 3.**
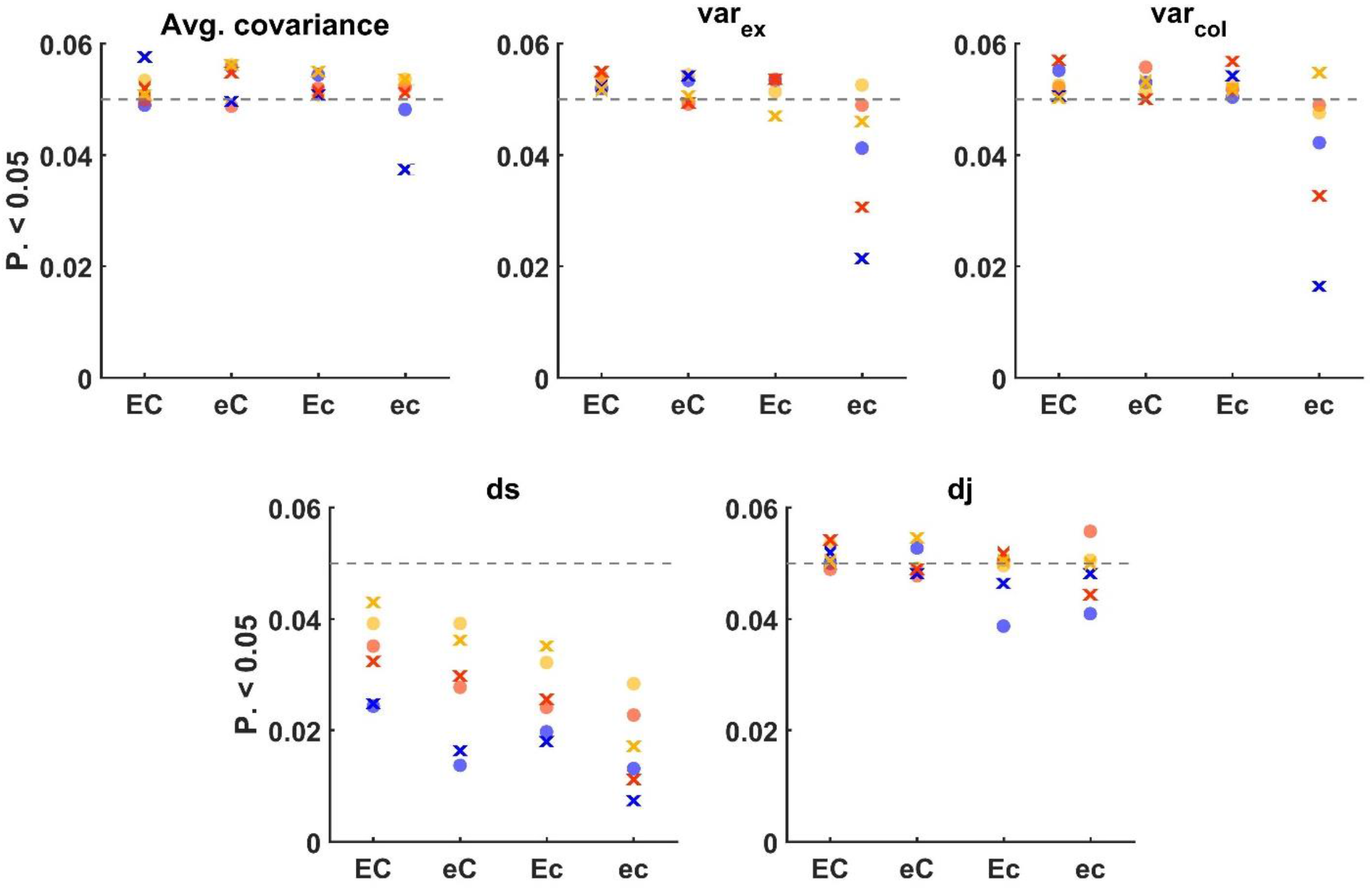
Proportion of rejected statistics under DE (type I errors) at various parameter regimes. Four combinations of rates are considered – high and low average colonization (C and c) and high and low average extinction (E and e). simulations were run for each combination of rates, correlation between colonization and extinction (‘o’ = no correlation, ‘x’ – negative correlation) and size of the species pool (30 = blue, 90 = orange, 270 = yellow). Every point represents one of these combinations, and the proportion of significant results was calculated over 5000 synthetic communities. The grey dashed line represents rejection probability of 0.05.

Under the alternative model, the methodology exhibited several desired properties, with the detailed analysis presented in Appendix S2. Generally, we found that the methodology has power to detect deviations from DE, and this power increases with the number of species in the pool, the rates (since more events mean more informative input to the methodology) and the magnitude of the deviations. We also examined cases in which only some of the assumptions are violated (e.g. the species are independent but the rates are not uniform), finding that a statistic quantifying violations from some assumption is mostly specific to this assumption and unsensitive to the violation of other assumptions.

### Guidelines for practical application of the methodology

Based on these and other tests of the performance, we would like to recommend a few guidelines for the application of the methodological framework. We recommend using data with at least ten samples in time (years) that has no seasonality and at least 20-30 species observed. This will give minimal power to detect deviations. Furthermore, having at least two periods of presence and two periods of absence (not necessarily for the same species) for some species is crucial for the PARIS null model to work. We further recommend using at least a few hundred resamples and k ≈ 0.06 (cutting 6% of samples on each side, which would give one year at the beginning and the end for a ten-year record) for the calculation of *ds*. Moreover, the values and significance of *var_ex_* and *var_col_* should be taken with a grain of salt in the presence of sampling errors, although they may still be useful to compare between different communities with similar levels of sampling errors. Regarding missing years, if type I errors for *dj* are a concern, we recommend using Table 2 as a guideline for the percent of maximal missing samples to be used. We recommend 5%-15%, depending on the total number of species observed. If type I errors for 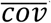 are a concern, a threshold of P = 0.08 may be used for significant results.

**Table 2.** Quantifying the deviations from DE in the Florida Keys experiment using our proposed methodology. For each island we present the five statistics quantifying deviations from the predictions (*ds*, *dj*) and assumptions (*var*_*ex*_, *Var*_*col*_, 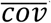) and their significance. We also present the difference between the observed average covariance and the expected average covariance in each island in the last column. We further present the average statistics over islands (and their significance). 2·10^4^ randomizations were used and k = 0.06 of the initial and final surveys were removed for calculating *ds*. *: P < 0.05, **: P < 0.02, ***: P < 0.005.

| *Island* | *ds* | *dj* | $var_{ex}$ | $Var_{col}$ | $\overline{cov}$ | $\overline{cov} - \overline{cov}_{null}$ |
| --- | --- | --- | --- | --- | --- | --- |
| **E1** | 0.37 | -0.027 | 1.268 | -0.315 | 0.02 | -0.003 |
| **E2** | 6.913 | 0.006 | 0.546 | -0.039 | 0.013 * | 0.004 * |
| **E3** | 6.927 * | 0.028 | 0.829 *** | -0.101 | 0.009 | 0.003 |
| **E7** | 6.027 | -0.015 | 0.251 | -0.01 | 0.014 ** | 0.005 ** |
| **E9** | 4.622 | -0.008 | 0.114 | 0.209 | 0.023 * | 0.006 * |
| **ST2** | -2.618 | -0.003 | 0.504 | -0.042 | 0.007 | -0.002 |
| **Average** | 3.707 ** | -0.003 | 0.585 *** | -0.05 | 0.014 | 0.002 |

Appendix S3 includes the full Matlab™ code used in our analyses. In particular, the *test_all4* function gets as input the number of resamples to perform, the parameter k and a matrix of the composition of species (columns) on years (rows), where ‘1’ represents presence, ‘0’ represents absence and a row of NaN values represents a missing year. The function calls the null2_comm3 function which resamples the communities using the PARIS algorithm, and exports the values and the significances of the five statistics. A simple example code (‘EXAMPLE.m’) is included in the appendix.

### Case study – Florida Keys experiment

The most classical test of DE was performed by Simberloff & Wilson (1969), who performed an experimental defaunation of six small mangrove-tree islands in the Florida Keys area. The arthropod composition and richness of the islands was surveyed before the defaunation and on a (roughly) regular basis for about a year. The experiment showed a gradual increase in richness to approximately the pre-defaunation richness, supporting the qualitative prediction of a richness equilibrium. Simberloff & Wilson (1969) and Simberloff (1969) also argued that the patterns of temporal dynamics, and specifically the curves of species richness through time (‘colonization curves’) are in good quantitative agreement with DE.

Here we revisit this classical dataset and examine the ability of our methodology to detect patterns of deviations from DE using the original data published by Simberloff & Wilson (1969). We applied our methodology to the six islands by calculating the five statistics and their significance for every island and the average values (and their significance) over all island. We further generated plots of the expected vs. observed species richness through time curves (which are used to calculate *ds*). We discarded the pre-defaunation survey and the last survey, which was done after a considerable time gap using a somewhat different methodology. We included all species whose presence was inferred but were not detected, which may be somewhat conservative, but in line with the original approach of Simberloff & Wilson (1969). As in the analysis of Simberloff (1969), we assumed that sampling was done at regular times, allowing us to use the PARIS null model.

We found that the curves of richness through time indeed qualitatively resembled the expected curves. However, we found that, on average, richness changes were significantly excessive (positive *ds*, Figure 4 and Table 2). For four of the islands, excess richness changes were 4.62-6.93, which are 18%-29% of the average richness, and this excess was statistically significant for one island. Island E1, whose colonization curve Simberloff and Wilson suggested does not fit the model, was in the best agreement with the null model, and island ST2 showed smaller changes then expected (although not significantly so). Unlike changes in richness, there was no significant excessive compositional turnover. Testing the assumptions of DE, we found that extinction rates were significantly not uniform on average (positive *var_ex_*) and for one of the islands, while colonization rates were not different from uniform. Finally, in three of the islands, species significantly positively covaried. Some level of positive covariance for this experiment is to be expected since most species were initially absent and are expected to arrive. However, this positive covariance was beyond the expectation of the null model, as seen in the last column of Table 2, which subtracted the expected covariance from the observed.

**Figure 4.**
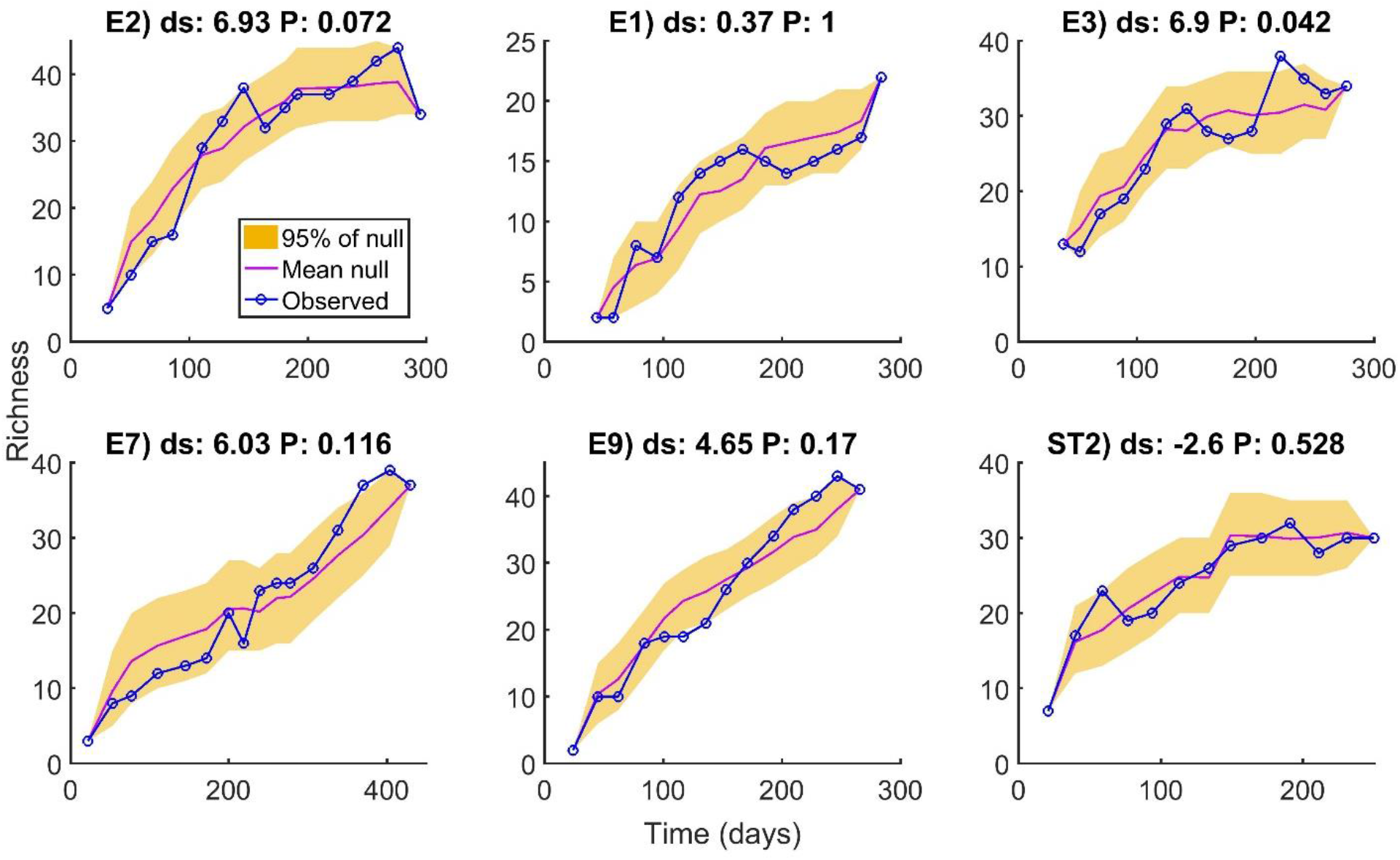
Curves of species richness through time in the six islands of the Florida Keys experiment. Each plot presents the observed richness (blue), the mean null (averaged over all resamples, purple) richness and the band capturing 95% of the resamples (orange). The excess richness changes statistic (*ds*) and its significance are presented for each island. 2·10^4^ randomizations were used.

## Discussion

### Recapitulation

Revisiting the requirements that we have set for a methodology for testing DE at the end of the introduction, we believe that our approach largely satisfies them all. It allows quantifying, easily and with readily available data, how different facets of empirically observed dynamics deviate from the expectations of DE. We have focused on statistics that quantify deviations from both the assumptions and the predictions. Since the two parts of the methodology, namely the null model and the set of statistics, are independent, it is possible to define multiple other statistics to test specific hypotheses of interests, using the PARIS null model as reference. Performing an extensive analysis of the framework under a variety of conditions, we have found that indeed, for data generated under DE, type I error rate never notably exceeds the predefined α = .05, and the average values of the statistics are very close to zero. Introducing a variety of issues in the data revealed some sensitivities in several cases, but the methodology was largely robust. Finally, at least for one generic alternative model, the more the dynamics deviate from DE, the larger the values of the statistics and statistical power will be.

We next examined the ability of our methodology to detect meaningful deviations from DE in a real dataset, the classical Florida Keys experiment. Since these islands are considered a classical quantitative support for DE, sample size is not large (13-16 surveys), and since we discarded the last survey (which was suggested by Wilson and Simberloff to represent different dynamics and whose data was collected differently) we did not expect to find major deviations from DE. Nevertheless, we found that, on average, changes in richness (a key prediction of the model) and variability in extinction rates (a key assumption of the model) are significantly larger than expected under the DE. We also found that the average covariance between species is often positive and significantly exceeded DE expectation in half of the islands. These overall results raise considerable concern about Simberloff and Wilson’s (1969) conclusion that their experimental results quantitatively support the DE model.

### Relation to previous methods

Attempts and approaches to test the dynamical predictions of DE go as far back as the original publication of the theory and its first tests. Here we recapitulate such approaches and compare them to our methodological framework.

The most straightforward approach to testing DE is running simulations of DE and comparing the resulting patterns to the observed patterns. In the original analysis of Simberloff (1969), he uses this approach to calculate confidence intervals for the curves of richness through time, finding that the confidence intervals were very wide, and could easily accommodate for the empirically observed curves. However, in this work there were no reliable estimates of the colonization and extinction rates to be used in the simulations. Later, Simberloff (1981, 1983) suggested using the maximum likelihood estimator (MLE) for each rate as the input for the simulations. For instance, considering extinction, this estimator is the proportion of presence observations followed by extinctions out of the total number of presence observations (that may be potentially followed by an extinction), and the analogous estimator (considering absence followed by colonization) is available for colonization rates (Simberloff 1981, 1983). However, our preliminary analyses have shown that the performance of this approach is upset by two different sources of bias. First, the number of species colonizing the community in a simulation cannot exceed the observed number, but it may be lower if some of the observed colonizers do not arrive. Furthermore, the rates of events are inherently upwards biased (by >30%) under multiple realistic scenarios (Figure S10). This stems from the fact that the time series ends abruptly, so the *full* duration of the final presence/absence event and the timing of its end are unknown, but its *observed* duration is included in the total time of presence/absence. For example, if a species tends to be regularly present for 10 years and then absent for 10 years, the data is 25 years in length, and the 10 years of the presence coincide with the beginning of the observation, we will observe one extinction event in 15 years rather than one in 10. On the other hand, with our approach one cannot get a deviation from the observed number of colonizing species or from the observed probability of extinction and colonization (in the entire timespan of the data), preventing both biases.

These biases interplay in a nontrivial manner with often harsh consequences. When we used DE simulations with rates estimated by the MLE as a null for out statistics, all the statistics quantifying the assumptions suffered from highly inflated type I error rates (Figure S11). Moreover, *var_ex_*, *var_col_* and *ds* sufferd from very hight effect size under DE, while *dj* often had highly negative effect sizes (Figure S12). Comparing statistical power for the statistics of the predictions (that did not have excessive type I errors) using the alternative SDE model, we found that *dj* was considerably stronger when PARIS was used as the null, while *ds* was stronger using DE simulations calibrated with maximum likelohood estimated rates. Nevertheless, using *ds* may be unwarranted because of the spurious finding of large effect sizes.

Other works have suggested global criteria for testing predictions of DE that do not fit null models to a specific dataset. Already MacArthur & Wilson (1963, 1967) have suggested an upper bounds on the Varince-Mean Ratio (VMR) in a richness time-series under DE, and this criterion has been in use since (Schoener 2010). They found that generally the VMR should be < 1 (although a lower upper bound may be found in restrictive conditions). We compared the statisticsl power of the VMR method to *ds* using the PARIS null model by comparing the ratio of communities with VMR > 1 to the communities with significant *ds* (Figure S13). *ds* is generally considerably more powerful, except when extinction rates are high and colonization rates are low, when VMR is sometimes more powerful. For this scenario the power of VMR is even stronger if the rates are uncorrelated (Figure S14), but this case is biologically less realistic.

A related approach is to test for temporal trends in richness. In this aproach, the lack of a trend is considered as support for equilibrium dynmics (althougth not nescesarilty DE, Rosenzweig 1995; Brown *et al.* 2001). However, tests are sometimes done by considering the statistical significance of the slope of OLS regression (e.g. Dornelas *et al.* 2014; Magurran *et al.* 2015). This approach does not consider the autocorrelation in the data (unlike our PARIS model, which generates autocorrelated time series). Moreover, if richness is initially not precisely at the equilibrium value, some short-term trend is expected. This essentially leads to very high (30%-70%, Figure S15) rates of type I error, with lower colonization and extinction rates leading to higher error rate. This stems from lower colonization and extinction rates having lower rate of return to equilibrium (MacArthur & Wilson 1967), leading to higher autocorrelation and more stochastic deviations from equilibrium.

The rates of compositional turnover in ecological communities have also been compared to the predictions of DE. Dornelas *et al.* (2014) used the linear slope of changes in Jaccard similarity though time w.r.t the initial survey, measuring this statistic for each of 100 communities and comparing to the expectations of DE. The latter were calculated by computing the slope for communities consisting of 50 species with rates uniformly distributed between zero and one over 300 time steps. We found that this methodology has a type I error rate of > 96% in all the scenarios we consider (Figure S16). This stems from the very high rates that were arbitrarily selected and because Jaccard similarity through time under DE is a nonlinear function that has an asymptote. Hence, the timescale over which a linear slope is fitted to this function is critically important, while the aforementioned approach uses 300 time steps arbitrarily.

### Interpretation and concluding remarks

An essential property of our methodology is that it extracts patterns that are informative of processes from the observed dynamics of single communities. A possible limitation of this approach is that it cannot detect effects that are static in time. If a biotic or a-biotic factor strongly affects community assembly but does not show temporal variation – our methodology cannot detect it. Examples of effects that cannot be detected include: species that are always present and exerts strong effects on other species; fixed environmental condition that affects assembly and so forth. For similar reasons, if a very rare species, arriving and disappearing, influences other species – the power to detect this will be low. It is also crucial to note that the rates which the null model uses are not the ‘basic’ colonization and extinction rates of the species, but are *already affected* by the biotic and abiotic environment. Therefore, good agreement with the null does not exclude effects of competition and certainly not static environmental filtering.

While considering only one community in time precludes the detection of patterns of assembly that are observed across multiple communities in space, this also has advantages. Community assembly studies that consider spatial variation in presence (Diamond 1975; Gotelli & McCabe 2002; Lyons *et al.* 2016) typically assume that the probability of a species to arrive in all the communities is identical. However, spatial structure in the species pool may lead to violations of this assumption. Our methodology, on the other hand, makes no assumptions regarding the pool, and unlike some of the aforementioned temporal approaches, it is fitted to a specific dataset, taking into account its timespan, colonization and extinction rates etc. Moreover, the analysis preserves the initial state of the community. As a result, our methodology is not sensitive to deviations from equilibrium and cannot be used to test whether the community is currently at equilibrium. This property is very important also because community dynamics generally depends on the initial state of the community. For example, richness trends and fast compositional turnover are expected if the community is initially not at equilibrium, and our methodology allows detecting deviations beyond those expected based on the initial state alone.

Ecological communities are highly variable in time and underpinning the mechanistic source of this variation is very challenging. Different communities can show different levels of temporal variation simply because of differences in rates, possibly due to different spatial scale. For example, a small community is expected to have higher rates of extinction than a large community. Moreover, when studying community dynamics, it is not even clear in many cases what empirical patterns to look at, and what insight can different points of view give. Our framework sets a reference point against which community dynamics can be studied, and since its statistics are relativistic, it can ‘filter’ some of the variation between communities that is simply a result of differences in rates, thus promoting a more solid understanding of community dynamics.

## Acknowledgements

We thank Micha Mandel for his help and consultation on the development of the methodological framework and Eyal Ben-Hur for his assistance in the graphical presentation of results. M.K. is supported by the Adams Fellowship Program of the Israel Academy of Sciences and Humanities.

