## Appendix S3 for "A framework for quantifying deviations from dynamic equilibrium theory": dist_pval.m

```
function [p] = dist_pval(dist,ref)
%dist_pval calculates a two sided p value based on comparing ref to a null
%distribution dist.

l=sum(~isnan(dist));
p=0;
if ref<nanmedian(dist)
    p=2*sum(dist<=ref)/l;
elseif ref>nanmedian(dist)
    p=2*sum(dist>=ref)/l;
elseif ref==nanmedian(dist)
    p=1;
end
   
end
```
