## Appendix S3 for "A framework for quantifying deviations from dynamic equilibrium theory": estimate_rates1.m

```
function [im,ex] = estimate_rates1(ts)
%estimate_rates1 estimates the vectors of colonization and extinction rates
%based on the simpe estimate, function 'measure_probs2'

sps=length(ts(1,:));
im=zeros(1,sps);
ex=zeros(1,sps);

for s=1:sps
    [im(s),ex(s)] = measure_probs2(ts(:,s));
end

end
```
