## Appendix S3 for "A framework for quantifying deviations from dynamic equilibrium theory": EXAMPLE.m

```
%generate a simple synthetic communnity time series and then apply the
%methodology:

% INPUT PARAMETERS:
im = 0.3; %immigration probability for all species
ex = 0.6; %extinction probability for all species
Sreg = 10; %species in pool
years = 20; %years in dataset
resamps = 200; %number of resamples
cut = 0.06; %percent of data to calculate ds

%GENERATE DATA:
dyn = MWDR_2(im*ones(Sreg,1) , ex*ones(Sreg,1) , binornd(1, (im/(im+ex))*ones(Sreg,1)) , 1 , years); %generate dynamics. initiate with the steady state distribution.

%COMPUTE AND TEST STATISTICS:
%the 'ds' statistic takes the cut parameter ('k' of the main text) to be
%0. dsc, on the contrary, uses the 'cut' parameter given above. Setting the
%last parameter, 'export_res' to 'false' prevents exporting the resampled
%statistics
[c_mean,c_mean_pval,var_ex,var_ex_pval,var_col,var_col_pval,ds,pval_ds,dsc,pval_dsc,dj,pval_dj,~,~,~,~,~,~] = test_all4(dyn,resamps,cut,false)
```
