## Appendix S3 for "A framework for quantifying deviations from dynamic equilibrium theory": generate_global_null.m

```
function [global_null] = generate_global_null()

% use the method of Dornelas et al. (2014, Science) to generate the
% distribution of expected slpes of similarity throuth time

global_null=zeros(1,1000);

parfor i=1:1000
    
    im=rand(50,1);
    ex=rand(50,1);
    isl_init=double(rand(50,1)<(im./(im+ex)));
    
    dyn_null=MWDR_2_mex(im,ex,isl_init,1,300);
    J_null=1-J_init(dyn_null);
    
    f=polyfit(1:299,J_null(2:end),1);
    global_null(i)=f(1);
    
end

end
```
