## Appendix S3 for "A framework for quantifying deviations from dynamic equilibrium theory": J_init.m

```
function [JT] = J_init(composition)
%J_init calculates the Jaccard dissimilarity of every time step compared to the first.
%Compisition is a 2D array of 1s and 0s where each row is a year and each
%column is a species.

t_tot=length(composition(:,1));
JT=zeros(1,t_tot);

for t=1:t_tot
    JT(t)=sum(abs(composition(t,:)-composition(1,:)))/(sum(abs(composition(t,:)-composition(1,:)))+sum((composition(t,:)+composition(1,:))==2));
end

end
```
