## Appendix S3 for "A framework for quantifying deviations from dynamic equilibrium theory": measure_probs2.m

```
function [c,e] = measure_probs2(ts)

if sum(ts)==length(ts) %only presence
    c=1;
    e=0;
elseif sum(ts)==0 %only absence
    c=0;
    e=1;
else %most cases, hopefully...

    %now calculate rates:
    d=diff(ts); %changes
    e=sum(d==-1)/sum(ts(1:end-1));
    c=sum(d==1)/sum(ts(1:end-1)==0);
    
    %account for case when only last year has '1'. if next check is not
    %used, we will get NaN in this case.
    if (sum(ts(1:end-1))==0)&&(ts(end)==1)
        e=0;
    end
    
    %analogy for colonization
    if (sum(ts(1:end-1))==(length(ts)-1))&&(ts(end)==0)
        c=0;
    end
    
end

end
```
