## Appendix S3 for "A framework for quantifying deviations from dynamic equilibrium theory": MWDR_2.m

```
function [composition] = MWDR_2(im,ex,isl_initial,t_initial,t_max)

%a discrete MW model. im and ex are column vectors of immigration and extinction
%probabilities. t_initial is the first year of sampling and t_max is the
%last year. the output, compotision, is a matrix where each row is a year
%and each column is a species.

S_reg=length(im);
isl=isl_initial; %generate initial island
composition=zeros(length(t_initial:t_max),S_reg);
composition(1,:)=isl;

for t=2:t_max
    %extinction, considering available species:
    f=find(isl); %indexes of available species
    %which species, from within f, will go extinct?
    to_ex=binornd(1,ex(f)',1,length(f));
    
    %colonization, considering non available species:
    f_comp=find(isl==0);
    to_im= binornd(1,im(f_comp)',1,length(f_comp));

    %perform processes:

    isl(f(logical(to_ex)))=0; %perform extinction using the indexes from f
    isl(f_comp(logical(to_im)))=1; %perform colonization
    
    if t>=t_initial %sample
        composition(t-t_initial+1,:)=isl;
    end

end
```
