## Appendix S3 for "A framework for quantifying deviations from dynamic equilibrium theory": null2_comm3.m

```
function [dyn_res]=null2_comm3(dyn)
%Resamples the entire community species by species. in the input, each row is a year and each column is a species

dyn_res=dyn;

for dd=1:length(dyn(1,:))
    dyn_res(:,dd)=null2_1ts3(dyn(:,dd)')';
    
end

end
```
