## Appendix S3 for "A framework for quantifying deviations from dynamic equilibrium theory": p_durations3.m

```
function [durations]=p_durations3(dyn)
% returns a vector of the durations of presence segments. this code removes
% all missing years (NaNs). dyn is a row vector of 1s and 0s.

dyn=[0 dyn 0];
dyn=dyn(~isnan(dyn));

d=diff(dyn);
durations=find(d<0)-find(d>0);

end
```
