## Appendix S3 for "A framework for quantifying deviations from dynamic equilibrium theory": plot_power_ratio_VMR.m

```
load('type2.mat')

%% plot for VMR vs dsc:

figure()

cc = 1; %correlation between rates

for rr = 1:4

    s = subplot(4,3,1 + (rr-1)*3);
    hold(s,'on');
    set(s,'FontName','Times New Roman','FontSize',18,'LineWidth',1.5);
    plot(c_levels,log((sum(s1(rr,cc).sc_dsc_pval < 0.05,3) + 1) ./ (sum(s1(rr,cc).sc_VM > 1,3) + 1) ),'-o','LineWidth',1.5)
    line(c_levels,zeros(1,length(c_levels)),'Color',[0.50 0.50 0.50],'LineWidth',1.5,'LineStyle','--')
 
    if rr == 4
        xlabel('c','FontWeight','bold')
    end
    
    if rr==1
        ylabel('EC','FontWeight','bold')
    elseif rr==2
        ylabel('eC','FontWeight','bold')
    elseif rr==3
        ylabel('Ec','FontWeight','bold')
    else
        ylabel('ec','FontWeight','bold')
    end

    s = subplot(4,3,2 + (rr-1)*3);
    hold(s,'on');
    set(s,'FontName','Times New Roman','FontSize',18,'LineWidth',1.5,'XTick',[1 2 3],'XTickLabel',{'1','2','S_r_e_g'});
    plot(g_levels,log((sum(s1(rr,cc).sg_dsc_pval < 0.05,3) + 1) ./ (sum(s1(rr,cc).sg_VM > 1,3) + 1) ),'-o','LineWidth',1.5)
    line(g_levels,zeros(1,length(g_levels)),'Color',[0.50 0.50 0.50],'LineWidth',1.5,'LineStyle','--')
    if rr == 4
        xlabel('g','FontWeight','bold')
    end

    s = subplot(4,3,3 + (rr-1)*3);
    hold(s,'on');
    set(s,'FontName','Times New Roman','FontSize',18,'LineWidth',1.5);
    plot(tau_levels,log((sum(s1(rr,cc).st_dsc_pval < 0.05,3) + 1) ./ (sum(s1(rr,cc).st_VM > 1,3) + 1) ),'-o','LineWidth',1.5)
    line(tau_levels,zeros(1,length(tau_levels)),'Color',[0.50 0.50 0.50],'LineWidth',1.5,'LineStyle','--')
    if rr == 4
        xlabel('\tau','FontWeight','bold')
    end

end
```
