## Appendix S3 for "A framework for quantifying deviations from dynamic equilibrium theory": plot_type1_dornelas.m

```
load('type1_1.mat')

%% plot of significance only:

f = figure()

%%%%%%plot average covariance:

axes1 = axes('Parent',f);
hold(axes1,'on');
hold on

for rr = 1:4
    scatter(rr*ones(3,1),sum(s1(rr,2).Dornelas_pval<0.05,2)/ittr,[],[0 0 1; 1 0.2 0; 1 0.7 0],'o','LineWidth',2)
    scatter(rr*ones(3,1),sum(s1(rr,1).Dornelas_pval<0.05,2)/ittr,[],[0 0 1; 1 0.2 0; 1 0.7 0],'x','LineWidth',2)
end

set(axes1,'XTick',[1 2 3 4],'XTickLabel',{'EC', 'eC', 'Ec', 'ec'},'FontSize',18,'FontWeight','bold','LineWidth',1.5);
xlim([0.5 4.5])
%ylim([0 0.06])
ylabel('Jaccard P < 0.05')
```
