## Appendix S3 for "A framework for quantifying deviations from dynamic equilibrium theory": plot_type1_holes.m

```
load('type1_holes.mat')

sss=s2;
c = 1; %level of correlation between rates
%% fix the data and transform each Pvalue where the effect size is NaN into NaN.
for rr=1:length(rates_levels(:,1))
    for cc=1:2
        sss(rr,cc).dj_pval(isnan(sss(rr,cc).dj))=NaN;
        sss(rr,cc).ds_pval(isnan(sss(rr,cc).ds))=NaN;
        sss(rr,cc).dsc_pval(isnan(sss(rr,cc).dsc))=NaN;
        sss(rr,cc).c_mean_pval(isnan(sss(rr,cc).c_mean))=NaN;
        sss(rr,cc).var_ex_pval(isnan(sss(rr,cc).var_ex))=NaN;
        sss(rr,cc).var_col_pval(isnan(sss(rr,cc).var_col))=NaN;
    end
end

%%

%plot the significance:

figure()

for rr=1:4
    
    %plot for dj:
    
    sub((rr-1)+1)=subplot(5,4,(rr-1)+1);
    plot(na_levels,sum(sss(rr,c).dj_pval<0.05,3)./sum(~isnan(sss(rr,c).dj),3),'-o','linewidth',1.5)
    set(sub((rr-1)+1),'FontName','Times New Roman','FontSize',16,'LineWidth',1.5)
    if rr==1
        ylabel('P dj<.05','fontsize',16,'FontWeight','bold')
    end
    ylim([0 0.2])
    line([0 10],[0.05 0.05],'LineStyle','--','Color',[.5 .5 .5])
    line([0 10],[0.075 0.075],'LineStyle','--','Color',[.5 .5 .5])
    line([0 10],[0.1 0.1],'LineStyle','--','Color',[.5 .5 .5])

    if rr==1
        title('EC','fontsize',16,'FontWeight','bold')
    elseif rr==2
        title('eC','fontsize',16,'FontWeight','bold')
    elseif rr==3
        title('Ec','fontsize',16,'FontWeight','bold')
    else
        title('ec','fontsize',16,'FontWeight','bold')
    end
    
    %plot for ds:
    
    sub((rr-1)+5)=subplot(5,4,(rr-1)+5);
    plot(na_levels,sum(sss(rr,c).dsc_pval<0.05,3)./sum(~isnan(sss(rr,c).dsc),3),'-o','linewidth',1.5)
    set(sub((rr-1)+5),'FontName','Times New Roman','FontSize',16,'LineWidth',1.5)
    if rr==1
        ylabel('P ds<.05','fontsize',16,'FontWeight','bold')
    end
    ylim([0 0.05])
    line([0 10],[0.05 0.05],'LineStyle','--','Color',[.5 .5 .5])
    line([0 10],[0.075 0.075],'LineStyle','--','Color',[.5 .5 .5])
    line([0 10],[0.1 0.1],'LineStyle','--','Color',[.5 .5 .5])
    
    %plot for var_col:
    
    sub((rr-1)+9)=subplot(5,4,(rr-1)+9);
    plot(na_levels,sum(sss(rr,c).var_col_pval<0.05,3)./sum(~isnan(sss(rr,c).var_col),3),'-o','linewidth',1.5)
    set(sub((rr-1)+9),'FontName','Times New Roman','FontSize',16,'LineWidth',1.5)
    if rr==1
        ylabel('P var_c_o_l<.05','fontsize',16,'FontWeight','bold')
    end
    ylim([0 0.1])
    line([0 10],[0.05 0.05],'LineStyle','--','Color',[.5 .5 .5])
    line([0 10],[0.075 0.075],'LineStyle','--','Color',[.5 .5 .5])
    line([0 10],[0.1 0.1],'LineStyle','--','Color',[.5 .5 .5])
    %plot for var_ex:
    
    sub((rr-1)+13)=subplot(5,4,(rr-1)+13);
    plot(na_levels,sum(sss(rr,c).var_ex_pval<0.05,3)./sum(~isnan(sss(rr,c).var_ex),3),'-o','linewidth',1.5)
    set(sub((rr-1)+13),'FontName','Times New Roman','FontSize',16,'LineWidth',1.5)
    if rr==1
        ylabel('P var_e_x<.05','fontsize',16,'FontWeight','bold')
    end
    ylim([0 0.1])
    line([0 10],[0.05 0.05],'LineStyle','--','Color',[.5 .5 .5])
    line([0 10],[0.075 0.075],'LineStyle','--','Color',[.5 .5 .5])
    line([0 10],[0.1 0.1],'LineStyle','--','Color',[.5 .5 .5])
    
    %plot for c mean:
    
    sub((rr-1)+17)=subplot(5,4,(rr-1)+17);
    plot(na_levels,sum(sss(rr,c).c_mean_pval<0.05,3)./sum(~isnan(sss(rr,c).c_mean),3),'-o','linewidth',1.5)
    set(sub((rr-1)+17),'FontName','Times New Roman','FontSize',16,'LineWidth',1.5)
    if rr==1
        ylabel('P avg. cov.<.05','fontsize',16,'FontWeight','bold')
    end
    if rr==2
        xlabel('Missing years','fontsize',16,'FontWeight','bold')
    end
    ylim([0.02 0.06])
    line([0 10],[0.05 0.05],'LineStyle','--','Color',[.5 .5 .5])
    line([0 10],[0.075 0.075],'LineStyle','--','Color',[.5 .5 .5])
    line([0 10],[0.1 0.1],'LineStyle','--','Color',[.5 .5 .5])
    
end

%% plot the STANDARDIZED effect size:

figure()

for rr=1:4
    
    %plot for dj:
    
    sub((rr-1)+1)=subplot(5,4,(rr-1)+1);
    plot(na_levels,nanmean(sss(rr,c).dj,3)./nanstd(sss(rr,c).dj,[],3),'-o','linewidth',1.5)
    set(sub((rr-1)+1),'FontName','Times New Roman','FontSize',16,'LineWidth',1.5)
    if rr==1
        ylabel('dj','fontsize',16,'FontWeight','bold')
    end
    ylim([-.1 .5])
    line([0 10],[0 0],'LineStyle','--','Color',[.5 .5 .5])
    line([0 10],[0.1 0.1],'LineStyle','--','Color',[.5 .5 .5])
    line([0 10],[0.2 0.2],'LineStyle','--','Color',[.5 .5 .5])

    if rr==1
        title('EC','fontsize',16,'FontWeight','bold')
    elseif rr==2
        title('eC','fontsize',16,'FontWeight','bold')
    elseif rr==3
        title('Ec','fontsize',16,'FontWeight','bold')
    else
        title('ec','fontsize',16,'FontWeight','bold')
    end
    
    %plot for dsc:
    
    sub((rr-1)+5)=subplot(5,4,(rr-1)+5);
    plot(na_levels,nanmean(sss(rr,c).dsc,3)./nanstd(sss(rr,c).dsc,[],3),'-o','linewidth',1.5)
    set(sub((rr-1)+5),'FontName','Times New Roman','FontSize',16,'LineWidth',1.5)
    if rr==1
        ylabel('ds','fontsize',16,'FontWeight','bold')
    end
    ylim([-.05 .03])
    line([0 10],[0 0],'LineStyle','--','Color',[.5 .5 .5])
    line([0 10],[0.1 0.1],'LineStyle','--','Color',[.5 .5 .5])
    line([0 10],[0.2 0.2],'LineStyle','--','Color',[.5 .5 .5])
    
    %plot for var_col:
    
    sub((rr-1)+9)=subplot(5,4,(rr-1)+9);
    plot(na_levels,nanmean(sss(rr,c).var_col,3)./nanstd(sss(rr,c).var_col,[],3),'-o','linewidth',1.5)
    set(sub((rr-1)+9),'FontName','Times New Roman','FontSize',16,'LineWidth',1.5)
    if rr==1
        ylabel('var_c_o_l','fontsize',16,'FontWeight','bold')
    end
    ylim([-.05 .03])
    line([0 10],[0 0],'LineStyle','--','Color',[.5 .5 .5])
    line([0 10],[0.1 0.1],'LineStyle','--','Color',[.5 .5 .5])
    line([0 10],[0.2 0.2],'LineStyle','--','Color',[.5 .5 .5])

    %plot for var_ex:
    
    sub((rr-1)+13)=subplot(5,4,(rr-1)+13);
    plot(na_levels,nanmean(sss(rr,c).var_ex,3)./nanstd(sss(rr,c).var_ex,[],3),'-o','linewidth',1.5)
    set(sub((rr-1)+13),'FontName','Times New Roman','FontSize',16,'LineWidth',1.5)
    if rr==1
        ylabel('var_e_x','fontsize',16,'FontWeight','bold')
    end
    ylim([-.05 .03])
    line([0 10],[0 0],'LineStyle','--','Color',[.5 .5 .5])
    line([0 10],[0.1 0.1],'LineStyle','--','Color',[.5 .5 .5])
    line([0 10],[0.2 0.2],'LineStyle','--','Color',[.5 .5 .5])
    
    %plot for c nanmean:
    
    sub((rr-1)+17)=subplot(5,4,(rr-1)+17);
    plot(na_levels,nanmean(sss(rr,c).c_mean,3)./nanstd(sss(rr,c).c_mean,[],3),'-o','linewidth',1.5)
    set(sub((rr-1)+17),'FontName','Times New Roman','FontSize',16,'LineWidth',1.5)
    if rr==1
        ylabel('Avg. cov.','fontsize',16,'FontWeight','bold')
    end
    if rr==2
        xlabel('Missing years','fontsize',16,'FontWeight','bold')
    end
    ylim([-.05 .03])
    line([0 10],[0 0],'LineStyle','--','Color',[.5 .5 .5])
    line([0 10],[0.1 0.1],'LineStyle','--','Color',[.5 .5 .5])
    line([0 10],[0.2 0.2],'LineStyle','--','Color',[.5 .5 .5])
end
```
