## Appendix S3 for "A framework for quantifying deviations from dynamic equilibrium theory": plot_type1_OLS.m

```
load('type1_1.mat')

%% plot of significance only:

f = figure()

%%%%%%plot average covariance:

axes1 = axes('Parent',f);
hold(axes1,'on');
hold on

for rr = 1:4
    scatter(rr*ones(3,1),sum(s1(rr,2).OLS_pval<0.05,2)/ittr,[],[0 0 1; 1 0.2 0; 1 0.7 0],'o','LineWidth',2)
    scatter(rr*ones(3,1),sum(s1(rr,1).OLS_pval<0.05,2)/ittr,[],[0 0 1; 1 0.2 0; 1 0.7 0],'x','LineWidth',2)
end

set(axes1,'XTick',[1 2 3 4],'XTickLabel',{'EC', 'eC', 'Ec', 'ec'},'FontSize',18,'FontWeight','bold','LineWidth',1.5);
xlim([0.5 4.5])
%ylim([0 0.06])
line([0.5 4.5],[0.05 0.05],'Color',[0.50 0.50 0.50],'LineWidth',1.5,'LineStyle','--')
line([0.5 4.5],[0.075 0.075],'Color',[0.50 0.50 0.50],'LineWidth',1.5,'LineStyle','--')
line([0.5 4.5],[0.1 0.1],'Color',[0.50 0.50 0.50],'LineWidth',1.5,'LineStyle','--')
ylabel('OLS P < 0.05')
```
