## Appendix S3 for "A framework for quantifying deviations from dynamic equilibrium theory": plot_type1_ver2_ML.m

```
load('type1_1.mat')

%% plot of significance only:

fig3 = figure()

%%%%%%plot average covariance:

sub1 = subplot(2,3,1,'Parent',fig3);
hold on

for rr = 1:4
    scatter(rr*ones(3,1),sum(s1(rr,2).c_mean_ML_pval<0.05,2)./sum(~isnan(s1(rr,2).c_mean_ML),2),[],[0 0 1; 1 0.2 0; 1 0.7 0],'o','LineWidth',2)
    scatter(rr*ones(3,1),sum(s1(rr,1).c_mean_ML_pval<0.05,2)./sum(~isnan(s1(rr,1).c_mean_ML),2),[],[0 0 1; 1 0.2 0; 1 0.7 0],'x','LineWidth',2)
end

set(sub1,'XTick',[1 2 3 4],'XTickLabel',{'EC', 'eC', 'Ec', 'ec'},'FontSize',18,'FontWeight','bold','LineWidth',1.5);
xlim([0.5 4.5])
%ylim([0 0.06])
line([0.5 4.5],[0.05 0.05],'Color',[0.50 0.50 0.50],'LineWidth',1.5,'LineStyle','--')
line([0.5 4.5],[0.075 0.075],'Color',[0.50 0.50 0.50],'LineWidth',1.5,'LineStyle','--')
line([0.5 4.5],[0.1 0.1],'Color',[0.50 0.50 0.50],'LineWidth',1.5,'LineStyle','--')
ylabel('P < 0.05')
title('Avg. cov.')

%%%%%%plot var_ex:

sub1 = subplot(2,3,2,'Parent',fig3);
hold on

for rr = 1:4
    scatter(rr*ones(3,1),sum(s1(rr,2).var_ex_ML_pval<0.05,2)./sum(~isnan(s1(rr,2).var_ex_ML),2),[],[0 0 1; 1 0.2 0; 1 0.7 0],'o','LineWidth',2)
    scatter(rr*ones(3,1),sum(s1(rr,1).var_ex_ML_pval<0.05,2)./sum(~isnan(s1(rr,1).var_ex_ML),2),[],[0 0 1; 1 0.2 0; 1 0.7 0],'x','LineWidth',2)
end

set(sub1,'XTick',[1 2 3 4],'XTickLabel',{'EC', 'eC', 'Ec', 'ec'},'FontSize',18,'FontWeight','bold','LineWidth',1.5);
xlim([0.5 4.5])
%ylim([0 0.06])
line([0.5 4.5],[0.05 0.05],'Color',[0.50 0.50 0.50],'LineWidth',1.5,'LineStyle','--')
line([0.5 4.5],[0.075 0.075],'Color',[0.50 0.50 0.50],'LineWidth',1.5,'LineStyle','--')
line([0.5 4.5],[0.1 0.1],'Color',[0.50 0.50 0.50],'LineWidth',1.5,'LineStyle','--')
ylabel('P < 0.05')
title('var_e_x')

%%%%%%plot var_col:

sub1 = subplot(2,3,3,'Parent',fig3);
hold on

for rr = 1:4
    scatter(rr*ones(3,1),sum(s1(rr,2).var_col_ML_pval<0.05,2)./sum(~isnan(s1(rr,2).var_col_ML),2),[],[0 0 1; 1 0.2 0; 1 0.7 0],'o','LineWidth',2)
    scatter(rr*ones(3,1),sum(s1(rr,1).var_col_ML_pval<0.05,2)./sum(~isnan(s1(rr,1).var_col_ML),2),[],[0 0 1; 1 0.2 0; 1 0.7 0],'x','LineWidth',2)
end

set(sub1,'XTick',[1 2 3 4],'XTickLabel',{'EC', 'eC', 'Ec', 'ec'},'FontSize',18,'FontWeight','bold','LineWidth',1.5);
xlim([0.5 4.5])
%ylim([0 0.06])
line([0.5 4.5],[0.05 0.05],'Color',[0.50 0.50 0.50],'LineWidth',1.5,'LineStyle','--')
line([0.5 4.5],[0.075 0.075],'Color',[0.50 0.50 0.50],'LineWidth',1.5,'LineStyle','--')
line([0.5 4.5],[0.1 0.1],'Color',[0.50 0.50 0.50],'LineWidth',1.5,'LineStyle','--')
ylabel('P < 0.05')
title('var_c_o_l')

%%%%%%plot dsc:

sub1 = subplot(2,3,4,'Parent',fig3);
hold on

for rr = 1:4
    scatter(rr*ones(3,1),sum(s1(rr,2).dsc_ML_pval<0.05,2)./sum(~isnan(s1(rr,2).dsc_ML),2),[],[0 0 1; 1 0.2 0; 1 0.7 0],'o','LineWidth',2)
    scatter(rr*ones(3,1),sum(s1(rr,1).dsc_ML_pval<0.05,2)./sum(~isnan(s1(rr,1).dsc_ML),2),[],[0 0 1; 1 0.2 0; 1 0.7 0],'x','LineWidth',2)
end

set(sub1,'XTick',[1 2 3 4],'XTickLabel',{'EC', 'eC', 'Ec', 'ec'},'FontSize',18,'FontWeight','bold','LineWidth',1.5);
xlim([0.5 4.5])
%ylim([0 0.06])
line([0.5 4.5],[0.05 0.05],'Color',[0.50 0.50 0.50],'LineWidth',1.5,'LineStyle','--')
line([0.5 4.5],[0.075 0.075],'Color',[0.50 0.50 0.50],'LineWidth',1.5,'LineStyle','--')
line([0.5 4.5],[0.1 0.1],'Color',[0.50 0.50 0.50],'LineWidth',1.5,'LineStyle','--')
ylabel('P < 0.05')
title('ds')


%%%%%%plot dj:

sub1 = subplot(2,3,5,'Parent',fig3);
hold on

for rr = 1:4
    scatter(rr*ones(3,1),sum(s1(rr,2).dj_ML_pval<0.05,2)./sum(~isnan(s1(rr,2).dj_ML),2),[],[0 0 1; 1 0.2 0; 1 0.7 0],'o','LineWidth',2)
    scatter(rr*ones(3,1),sum(s1(rr,1).dj_ML_pval<0.05,2)./sum(~isnan(s1(rr,1).dj_ML),2),[],[0 0 1; 1 0.2 0; 1 0.7 0],'x','LineWidth',2)
end

set(sub1,'XTick',[1 2 3 4],'XTickLabel',{'EC', 'eC', 'Ec', 'ec'},'FontSize',18,'FontWeight','bold','LineWidth',1.5);
xlim([0.5 4.5])
%ylim([0 0.06])
line([0.5 4.5],[0.05 0.05],'Color',[0.50 0.50 0.50],'LineWidth',1.5,'LineStyle','--')
line([0.5 4.5],[0.075 0.075],'Color',[0.50 0.50 0.50],'LineWidth',1.5,'LineStyle','--')
line([0.5 4.5],[0.1 0.1],'Color',[0.50 0.50 0.50],'LineWidth',1.5,'LineStyle','--')
ylabel('P < 0.05')
title('dj')

hold off

%% Figure for effect size:

fig3 = figure()

%%%%%%plot average covariance:

sub1 = subplot(2,3,1,'Parent',fig3);
hold on

for rr = 1:4
    scatter(rr*ones(3,1),nanmean(s1(rr,2).c_mean_ML,2)./nanstd(s1(rr,2).c_mean_ML,0,2),[],[0 0 1; 1 0.2 0; 1 0.7 0],'o','LineWidth',2)
    scatter(rr*ones(3,1),nanmean(s1(rr,1).c_mean_ML,2)./nanstd(s1(rr,1).c_mean_ML,0,2),[],[0 0 1; 1 0.2 0; 1 0.7 0],'x','LineWidth',2)
end

set(sub1,'XTick',[1 2 3 4],'XTickLabel',{'EC', 'eC', 'Ec', 'ec'},'FontSize',18,'FontWeight','bold','LineWidth',1.5);
xlim([0.5 4.5])
line([0.5 4.5],[0 0],'Color',[0.50 0.50 0.50],'LineWidth',1.5,'LineStyle','--')
ylabel('SES')
title('Avg. cov.')


%%%%%%plot var_ex:

sub1 = subplot(2,3,2,'Parent',fig3);
hold on

for rr = 1:4
    scatter(rr*ones(3,1),nanmean(s1(rr,2).var_ex_ML,2)./nanstd(s1(rr,2).var_ex_ML,0,2),[],[0 0 1; 1 0.2 0; 1 0.7 0],'o','LineWidth',2)
    scatter(rr*ones(3,1),nanmean(s1(rr,1).var_ex_ML,2)./nanstd(s1(rr,1).var_ex_ML,0,2),[],[0 0 1; 1 0.2 0; 1 0.7 0],'x','LineWidth',2)
end

set(sub1,'XTick',[1 2 3 4],'XTickLabel',{'EC', 'eC', 'Ec', 'ec'},'FontSize',18,'FontWeight','bold','LineWidth',1.5);
xlim([0.5 4.5])
line([0.5 4.5],[-.2 -.2],'Color',[0.50 0.50 0.50],'LineWidth',1.5,'LineStyle','--')
line([0.5 4.5],[-.1 -.1],'Color',[0.50 0.50 0.50],'LineWidth',1.5,'LineStyle','--')
line([0.5 4.5],[0 0],'Color',[0.50 0.50 0.50],'LineWidth',1.5,'LineStyle','--')
line([0.5 4.5],[.2 .2],'Color',[0.50 0.50 0.50],'LineWidth',1.5,'LineStyle','--')
line([0.5 4.5],[.1 .1],'Color',[0.50 0.50 0.50],'LineWidth',1.5,'LineStyle','--')
ylabel('SES')
title('var_e_x')


%%%%%%plot var_col:

sub1 = subplot(2,3,3,'Parent',fig3);
hold on

for rr = 1:4
    scatter(rr*ones(3,1),nanmean(s1(rr,2).var_col_ML,2)./nanstd(s1(rr,2).var_col_ML,0,2),[],[0 0 1; 1 0.2 0; 1 0.7 0],'o','LineWidth',2)
    scatter(rr*ones(3,1),nanmean(s1(rr,1).var_col_ML,2)./nanstd(s1(rr,1).var_col_ML,0,2),[],[0 0 1; 1 0.2 0; 1 0.7 0],'x','LineWidth',2)
end

set(sub1,'XTick',[1 2 3 4],'XTickLabel',{'EC', 'eC', 'Ec', 'ec'},'FontSize',18,'FontWeight','bold','LineWidth',1.5);
xlim([0.5 4.5])
line([0.5 4.5],[-.2 -.2],'Color',[0.50 0.50 0.50],'LineWidth',1.5,'LineStyle','--')
line([0.5 4.5],[-.1 -.1],'Color',[0.50 0.50 0.50],'LineWidth',1.5,'LineStyle','--')
line([0.5 4.5],[0 0],'Color',[0.50 0.50 0.50],'LineWidth',1.5,'LineStyle','--')
line([0.5 4.5],[.2 .2],'Color',[0.50 0.50 0.50],'LineWidth',1.5,'LineStyle','--')
line([0.5 4.5],[.1 .1],'Color',[0.50 0.50 0.50],'LineWidth',1.5,'LineStyle','--')
ylabel('SES')
title('var_c_o_l')

%%%%%%plot dsc:

sub1 = subplot(2,3,4,'Parent',fig3);
hold on

for rr = 1:4
    scatter(rr*ones(3,1),nanmean(s1(rr,2).dsc_ML,2)./nanstd(s1(rr,2).dsc_ML,0,2),[],[0 0 1; 1 0.2 0; 1 0.7 0],'o','LineWidth',2)
    scatter(rr*ones(3,1),nanmean(s1(rr,1).dsc_ML,2)./nanstd(s1(rr,1).dsc_ML,0,2),[],[0 0 1; 1 0.2 0; 1 0.7 0],'x','LineWidth',2)
end

set(sub1,'XTick',[1 2 3 4],'XTickLabel',{'EC', 'eC', 'Ec', 'ec'},'FontSize',18,'FontWeight','bold','LineWidth',1.5);
xlim([0.5 4.5])
line([0.5 4.5],[-.2 -.2],'Color',[0.50 0.50 0.50],'LineWidth',1.5,'LineStyle','--')
line([0.5 4.5],[-.1 -.1],'Color',[0.50 0.50 0.50],'LineWidth',1.5,'LineStyle','--')
line([0.5 4.5],[0 0],'Color',[0.50 0.50 0.50],'LineWidth',1.5,'LineStyle','--')
line([0.5 4.5],[.2 .2],'Color',[0.50 0.50 0.50],'LineWidth',1.5,'LineStyle','--')
line([0.5 4.5],[.1 .1],'Color',[0.50 0.50 0.50],'LineWidth',1.5,'LineStyle','--')
ylabel('SES')
title('ds')


%%%%%%plot dj:

sub1 = subplot(2,3,5,'Parent',fig3);
hold on

for rr = 1:4
    scatter(rr*ones(3,1),nanmean(s1(rr,2).dj_ML,2)./nanstd(s1(rr,2).dj_ML,0,2),[],[0 0 1; 1 0.2 0; 1 0.7 0],'o','LineWidth',2)
    scatter(rr*ones(3,1),nanmean(s1(rr,1).dj_ML,2)./nanstd(s1(rr,1).dj_ML,0,2),[],[0 0 1; 1 0.2 0; 1 0.7 0],'x','LineWidth',2)
end

set(sub1,'XTick',[1 2 3 4],'XTickLabel',{'EC', 'eC', 'Ec', 'ec'},'FontSize',18,'FontWeight','bold','LineWidth',1.5);
xlim([0.5 4.5])
line([0.5 4.5],[-.2 -.2],'Color',[0.50 0.50 0.50],'LineWidth',1.5,'LineStyle','--')
line([0.5 4.5],[-.1 -.1],'Color',[0.50 0.50 0.50],'LineWidth',1.5,'LineStyle','--')
line([0.5 4.5],[0 0],'Color',[0.50 0.50 0.50],'LineWidth',1.5,'LineStyle','--')
line([0.5 4.5],[.2 .2],'Color',[0.50 0.50 0.50],'LineWidth',1.5,'LineStyle','--')
line([0.5 4.5],[.1 .1],'Color',[0.50 0.50 0.50],'LineWidth',1.5,'LineStyle','--')
ylabel('SES')
title('dj')

hold off
```
