## Appendix S3 for "A framework for quantifying deviations from dynamic equilibrium theory": plot_type1_ver3.m

```
load('type1_1.mat')

%% plot of significance only:

fig3 = figure()

%%%%%%plot average covariance:

sub1 = subplot(2,3,1,'Parent',fig3);
hold on

for rr = 1:4
    scatter(rr*ones(3,1),sum(s1(rr,2).c_mean_pval<0.05,2)./sum(~isnan(s1(rr,2).c_mean),2),80,[0 0 1; 1 0.2 0; 1 0.7 0],'filled','o','LineWidth',2,  'MarkerFaceAlpha', .6)
    scatter(rr*ones(3,1),sum(s1(rr,1).c_mean_pval<0.05,2)./sum(~isnan(s1(rr,1).c_mean),2),100,[0 0 1; 1 0.2 0; 1 0.7 0],'x','LineWidth',3)
end

set(sub1,'XTick',[1 2 3 4],'XTickLabel',{'EC', 'eC', 'Ec', 'ec'},'FontSize',18,'FontWeight','bold','LineWidth',1.5);
xlim([0.5 4.5])
ylim([0 0.06])
line([0.5 4.5],[0.05 0.05],'Color',[0.50 0.50 0.50],'LineWidth',1.5,'LineStyle','--')
ylabel('P < 0.05')
title('Avg. cov.')


%%%%%%plot var_ex:

sub1 = subplot(2,3,2,'Parent',fig3);
hold on

for rr = 1:4
    scatter(rr*ones(3,1),sum(s1(rr,2).var_ex_pval<0.05,2)./sum(~isnan(s1(rr,2).var_ex),2),80,[0 0 1; 1 0.2 0; 1 0.7 0],'filled','o','LineWidth',2,  'MarkerFaceAlpha', .6)
    scatter(rr*ones(3,1),sum(s1(rr,1).var_ex_pval<0.05,2)./sum(~isnan(s1(rr,1).var_ex),2),100,[0 0 1; 1 0.2 0; 1 0.7 0],'x','LineWidth',3)
end

set(sub1,'XTick',[1 2 3 4],'XTickLabel',{'EC', 'eC', 'Ec', 'ec'},'FontSize',18,'FontWeight','bold','LineWidth',1.5);
xlim([0.5 4.5])
ylim([0 0.06])
line([0.5 4.5],[0.05 0.05],'Color',[0.50 0.50 0.50],'LineWidth',1.5,'LineStyle','--')
%ylabel('P. < 0.05')
title('var_e_x')


%%%%%%plot var_col:

sub1 = subplot(2,3,3,'Parent',fig3);
hold on

for rr = 1:4
    scatter(rr*ones(3,1),sum(s1(rr,2).var_col_pval<0.05,2)./sum(~isnan(s1(rr,2).var_col),2),80,[0 0 1; 1 0.2 0; 1 0.7 0],'filled','o','LineWidth',2,  'MarkerFaceAlpha', .6)
    scatter(rr*ones(3,1),sum(s1(rr,1).var_col_pval<0.05,2)./sum(~isnan(s1(rr,1).var_col),2),100,[0 0 1; 1 0.2 0; 1 0.7 0],'x','LineWidth',3)
end

set(sub1,'XTick',[1 2 3 4],'XTickLabel',{'EC', 'eC', 'Ec', 'ec'},'FontSize',18,'FontWeight','bold','LineWidth',1.5);
xlim([0.5 4.5])
ylim([0 0.06])
line([0.5 4.5],[0.05 0.05],'Color',[0.50 0.50 0.50],'LineWidth',1.5,'LineStyle','--')
%ylabel('P < 0.05')
title('var_c_o_l')

%%%%%%plot dsc:

sub1 = subplot(2,3,4.5,'Parent',fig3);
hold on

for rr = 1:4
    scatter(rr*ones(3,1),sum(s1(rr,2).dsc_pval<0.05,2)./sum(~isnan(s1(rr,2).dsc),2),80,[0 0 1; 1 0.2 0; 1 0.7 0],'filled','o','LineWidth',2,  'MarkerFaceAlpha', .6)
    scatter(rr*ones(3,1),sum(s1(rr,1).dsc_pval<0.05,2)./sum(~isnan(s1(rr,1).dsc),2),100,[0 0 1; 1 0.2 0; 1 0.7 0],'x','LineWidth',3)
end

set(sub1,'XTick',[1 2 3 4],'XTickLabel',{'EC', 'eC', 'Ec', 'ec'},'FontSize',18,'FontWeight','bold','LineWidth',1.5);
xlim([0.5 4.5])
ylim([0 0.06])
line([0.5 4.5],[0.05 0.05],'Color',[0.50 0.50 0.50],'LineWidth',1.5,'LineStyle','--')
ylabel('P < 0.05')
title('ds')


%%%%%%plot dj:

sub1 = subplot(2,3,5.5,'Parent',fig3);
hold on

for rr = 1:4
    scatter(rr*ones(3,1),sum(s1(rr,2).dj_pval<0.05,2)./sum(~isnan(s1(rr,2).dj),2),80,[0 0 1; 1 0.2 0; 1 0.7 0],'filled','o','LineWidth',2,  'MarkerFaceAlpha', .6)
    scatter(rr*ones(3,1),sum(s1(rr,1).dj_pval<0.05,2)./sum(~isnan(s1(rr,1).dj),2),100,[0 0 1; 1 0.2 0; 1 0.7 0],'x','LineWidth',3)
end

set(sub1,'XTick',[1 2 3 4],'XTickLabel',{'EC', 'eC', 'Ec', 'ec'},'FontSize',18,'FontWeight','bold','LineWidth',1.5);
xlim([0.5 4.5])
ylim([0 0.06])
line([0.5 4.5],[0.05 0.05],'Color',[0.50 0.50 0.50],'LineWidth',1.5,'LineStyle','--')
%ylabel('P. < 0.05')
title('dj')

hold off

%% Figure for effect size:

fig3 = figure()

%%%%%%plot average covariance:

sub1 = subplot(2,3,1,'Parent',fig3);
hold on

for rr = 1:4
    scatter(rr*ones(3,1),nanmean(s1(rr,2).c_mean,2),80,[0 0 1; 1 0.2 0; 1 0.7 0],'filled','o','LineWidth',2,  'MarkerFaceAlpha', .6)
    scatter(rr*ones(3,1),nanmean(s1(rr,1).c_mean,2),100,[0 0 1; 1 0.2 0; 1 0.7 0],'x','LineWidth',3)
end

set(sub1,'XTick',[1 2 3 4],'XTickLabel',{'EC', 'eC', 'Ec', 'ec'},'FontSize',18,'FontWeight','bold','LineWidth',1.5);
xlim([0.5 4.5])
line([0.5 4.5],[0 0],'Color',[0.50 0.50 0.50],'LineWidth',1.5,'LineStyle','--')
ylabel('Average value')
title('Avg. cov.')


%%%%%%plot var_ex:

sub1 = subplot(2,3,2,'Parent',fig3);
hold on

for rr = 1:4
    scatter(rr*ones(3,1),nanmean(s1(rr,2).var_ex,2),80,[0 0 1; 1 0.2 0; 1 0.7 0],'filled','o','LineWidth',2,  'MarkerFaceAlpha', .6)
    scatter(rr*ones(3,1),nanmean(s1(rr,1).var_ex,2),100,[0 0 1; 1 0.2 0; 1 0.7 0],'x','LineWidth',3)
end

set(sub1,'XTick',[1 2 3 4],'XTickLabel',{'EC', 'eC', 'Ec', 'ec'},'FontSize',18,'FontWeight','bold','LineWidth',1.5);
xlim([0.5 4.5])
line([0.5 4.5],[0 0],'Color',[0.50 0.50 0.50],'LineWidth',1.5,'LineStyle','--')
%ylabel('Average')
title('var_e_x')


%%%%%%plot var_col:

sub1 = subplot(2,3,3,'Parent',fig3);
hold on

for rr = 1:4
    scatter(rr*ones(3,1),nanmean(s1(rr,2).var_col,2),80,[0 0 1; 1 0.2 0; 1 0.7 0],'filled','o','LineWidth',2,  'MarkerFaceAlpha', .6)
    scatter(rr*ones(3,1),nanmean(s1(rr,1).var_col,2),100,[0 0 1; 1 0.2 0; 1 0.7 0],'x','LineWidth',3)
end

set(sub1,'XTick',[1 2 3 4],'XTickLabel',{'EC', 'eC', 'Ec', 'ec'},'FontSize',18,'FontWeight','bold','LineWidth',1.5);
xlim([0.5 4.5])
line([0.5 4.5],[0 0],'Color',[0.50 0.50 0.50],'LineWidth',1.5,'LineStyle','--')
%ylabel('Average')
title('var_c_o_l')

%%%%%%plot dsc:

sub1 = subplot(2,3,4.5,'Parent',fig3);
hold on

for rr = 1:4
    scatter(rr*ones(3,1),nanmean(s1(rr,2).dsc,2),80,[0 0 1; 1 0.2 0; 1 0.7 0],'filled','o','LineWidth',2,  'MarkerFaceAlpha', .6)
    scatter(rr*ones(3,1),nanmean(s1(rr,1).dsc,2),100,[0 0 1; 1 0.2 0; 1 0.7 0],'x','LineWidth',3)
end

set(sub1,'XTick',[1 2 3 4],'XTickLabel',{'EC', 'eC', 'Ec', 'ec'},'FontSize',18,'FontWeight','bold','LineWidth',1.5);
xlim([0.5 4.5])
line([0.5 4.5],[0 0],'Color',[0.50 0.50 0.50],'LineWidth',1.5,'LineStyle','--')
ylabel('Average value')
title('ds')


%%%%%%plot dj:

sub1 = subplot(2,3,5.5,'Parent',fig3);
hold on

for rr = 1:4
    scatter(rr*ones(3,1),nanmean(s1(rr,2).dj,2),80,[0 0 1; 1 0.2 0; 1 0.7 0],'filled','o','LineWidth',2,  'MarkerFaceAlpha', .6)
    scatter(rr*ones(3,1),nanmean(s1(rr,1).dj,2),100,[0 0 1; 1 0.2 0; 1 0.7 0],'x','LineWidth',3)
end

set(sub1,'XTick',[1 2 3 4],'XTickLabel',{'EC', 'eC', 'Ec', 'ec'},'FontSize',18,'FontWeight','bold','LineWidth',1.5);
xlim([0.5 4.5])
line([0.5 4.5],[0 0],'Color',[0.50 0.50 0.50],'LineWidth',1.5,'LineStyle','--')
%ylabel('Average')
title('dj')

hold off
```
