## Appendix S3 for "A framework for quantifying deviations from dynamic equilibrium theory": plot_type2_SDE_ver2.m

```
%%

load('type2.mat')

%% PLOT OF EFFECT SIZE:

figure()
%determine which rate setting and correlation setting:
rr=1;
cc=1;

%row 1: dj:

s = subplot(5,3,1);
hold(s,'on');
set(s,'FontName','Times New Roman','FontSize',16,'LineWidth',1.5);
plot(c_levels,nanmean(s1(rr,cc).sc_dj,3),'-o','LineWidth',1.5)
ylabel('dj')

s = subplot(5,3,2);
hold(s,'on');
set(s,'FontName','Times New Roman','FontSize',16,'LineWidth',1.5,'XTick',[1 2 3],'XTickLabel',{'1','2','S_r_e_g'});
plot(g_levels,nanmean(s1(rr,cc).sg_dj,3),'-o','LineWidth',1.5)

s = subplot(5,3,3);
hold(s,'on');
set(s,'FontName','Times New Roman','FontSize',16,'LineWidth',1.5);
plot(tau_levels,nanmean(s1(rr,cc).st_dj,3),'-o','LineWidth',1.5)

%plot for ds:

s = subplot(5,3,4);
hold(s,'on');
set(s,'FontName','Times New Roman','FontSize',16,'LineWidth',1.5);
plot(c_levels,nanmean(s1(rr,cc).sc_dsc,3),'-o','LineWidth',1.5)
ylabel('ds')

s = subplot(5,3,5);
hold(s,'on');
set(s,'FontName','Times New Roman','FontSize',16,'LineWidth',1.5,'XTick',[1 2 3],'XTickLabel',{'1','2','S_r_e_g'});
plot(g_levels,nanmean(s1(rr,cc).sg_dsc,3),'-o','LineWidth',1.5)

s = subplot(5,3,6);
hold(s,'on');
set(s,'FontName','Times New Roman','FontSize',16,'LineWidth',1.5);
plot(tau_levels,nanmean(s1(rr,cc).st_dsc,3),'-o','LineWidth',1.5)

%plot for var_col:

s = subplot(5,3,7);
hold(s,'on');
set(s,'FontName','Times New Roman','FontSize',16,'LineWidth',1.5);
plot(c_levels,nanmean(s1(rr,cc).sc_var_col,3),'-o','LineWidth',1.5)
ylabel('var_c_o_l')

s = subplot(5,3,8);
hold(s,'on');
set(s,'FontName','Times New Roman','FontSize',16,'LineWidth',1.5,'XTick',[1 2 3],'XTickLabel',{'1','2','S_r_e_g'});
plot(g_levels,nanmean(s1(rr,cc).sg_var_col,3),'-o','LineWidth',1.5)

s = subplot(5,3,9);
hold(s,'on');
set(s,'FontName','Times New Roman','FontSize',16,'LineWidth',1.5);
plot(tau_levels,nanmean(s1(rr,cc).st_var_col,3),'-o','LineWidth',1.5)

%plot for var_ex:

s = subplot(5,3,10);
hold(s,'on');
set(s,'FontName','Times New Roman','FontSize',16,'LineWidth',1.5);
plot(c_levels,nanmean(s1(rr,cc).sc_var_ex,3),'-o','LineWidth',1.5)
ylabel('var_e_x')

s = subplot(5,3,11);
hold(s,'on');
set(s,'FontName','Times New Roman','FontSize',16,'LineWidth',1.5,'XTick',[1 2 3],'XTickLabel',{'1','2','S_r_e_g'});
plot(g_levels,nanmean(s1(rr,cc).sg_var_ex,3),'-o','LineWidth',1.5)
xlabel('g')

s = subplot(5,3,12);
hold(s,'on');
set(s,'FontName','Times New Roman','FontSize',16,'LineWidth',1.5);
plot(tau_levels,nanmean(s1(rr,cc).st_var_ex,3),'-o','LineWidth',1.5)
xlabel('tau')

%plot for c mean:

s = subplot(5,3,13);
hold(s,'on');
set(s,'FontName','Times New Roman','FontSize',16,'LineWidth',1.5);
plot(c_levels,nanmean(s1(rr,cc).sc_c_mean,3),'-o','LineWidth',1.5)
xlabel('c')
ylabel('Avg. cov.')

s = subplot(5,3,14);
hold(s,'on');
set(s,'FontName','Times New Roman','FontSize',16,'LineWidth',1.5,'XTick',[1 2 3],'XTickLabel',{'1','2','S_r_e_g'}   );
plot(g_levels,nanmean(s1(rr,cc).sg_c_mean,3),'-o','LineWidth',1.5)
xlabel('g')

s = subplot(5,3,15);
hold(s,'on');
set(s,'FontName','Times New Roman','FontSize',16,'LineWidth',1.5);
plot(tau_levels,nanmean(s1(rr,cc).st_c_mean,3),'-o','LineWidth',1.5)
xlabel('\tau')
legend({'S_r_e_g = 30', 'S_r_e_g = 90', 'S_r_e_g = 270'},'Orientation','horizontal')

%% PLOT OF SIGNIFICANCE:

figure()
%determine which rate setting and correlation setting:

%row 1: dJ:

s = subplot(5,3,1);
hold(s,'on');
set(s,'FontName','Times New Roman','FontSize',16,'LineWidth',1.5);
plot(c_levels,sum(s1(rr,cc).sc_dj_pval<=0.05,3)./sum(~isnan(s1(rr,cc).sc_dj),3),'-o','LineWidth',1.5) %each is divided 
ylabel('dj P<0.05')
line(c_levels,[0.05 0.05 0.05 0.05 ; 0.075 0.075 0.075 0.075 ; 0.1 0.1 0.1 0.1],'LineStyle','--','Color',[.5 .5 .5])

s = subplot(5,3,2);
hold(s,'on');
set(s,'FontName','Times New Roman','FontSize',16,'LineWidth',1.5,'XTick',[1 2 3],'XTickLabel',{'1','2','S_r_e_g'});
plot(g_levels,sum(s1(rr,cc).sg_dj_pval<=0.05,3)./sum(~isnan(s1(rr,cc).sg_dj),3),'-o','LineWidth',1.5)
line(g_levels,[0.05 0.05 0.05 ; 0.075 0.075 0.075 ; 0.1 0.1 0.1]','LineStyle','--','Color',[.5 .5 .5])

s = subplot(5,3,3);
hold(s,'on');
set(s,'FontName','Times New Roman','FontSize',16,'LineWidth',1.5);
plot(tau_levels,sum(s1(rr,cc).st_dj_pval<=0.05,3)./sum(~isnan(s1(rr,cc).st_dj),3),'-o','LineWidth',1.5)
line(tau_levels,[0.05 0.05 0.05 ; 0.075 0.075 0.075 ; 0.1 0.1 0.1]','LineStyle','--','Color',[.5 .5 .5])

%plot for ds

s = subplot(5,3,4);
hold(s,'on');
set(s,'FontName','Times New Roman','FontSize',16,'LineWidth',1.5);
plot(c_levels,sum(s1(rr,cc).sc_dsc_pval<=0.05,3)./sum(~isnan(s1(rr,cc).sc_dsc),3),'-o','LineWidth',1.5)
ylabel('ds P<0.05')
line(c_levels,[0.05 0.05 0.05 0.05 ; 0.075 0.075 0.075 0.075 ; 0.1 0.1 0.1 0.1],'LineStyle','--','Color',[.5 .5 .5])

s = subplot(5,3,5);
hold(s,'on');
set(s,'FontName','Times New Roman','FontSize',16,'LineWidth',1.5,'XTick',[1 2 3],'XTickLabel',{'1','2','S_r_e_g'});
plot(g_levels,sum(s1(rr,cc).sg_dsc_pval<=0.05,3)./sum(~isnan(s1(rr,cc).sg_dsc),3),'-o','LineWidth',1.5)
line(g_levels,[0.05 0.05 0.05 ; 0.075 0.075 0.075 ; 0.1 0.1 0.1]','LineStyle','--','Color',[.5 .5 .5])

s = subplot(5,3,6);
hold(s,'on');
set(s,'FontName','Times New Roman','FontSize',16,'LineWidth',1.5);
plot(tau_levels,sum(s1(rr,cc).st_dsc_pval<=0.05,3)./sum(~isnan(s1(rr,cc).st_dsc),3),'-o','LineWidth',1.5)
line(tau_levels,[0.05 0.05 0.05 ; 0.075 0.075 0.075 ; 0.1 0.1 0.1]','LineStyle','--','Color',[.5 .5 .5])

%plot for var_col:

s = subplot(5,3,7);
hold(s,'on');
set(s,'FontName','Times New Roman','FontSize',16,'LineWidth',1.5);
plot(c_levels,sum(s1(rr,cc).sc_var_col_pval<=0.05,3)./sum(~isnan(s1(rr,cc).sc_var_col),3),'-o','LineWidth',1.5)
line(c_levels,[0.05 0.05 0.05 0.05 ; 0.075 0.075 0.075 0.075 ; 0.1 0.1 0.1 0.1],'LineStyle','--','Color',[.5 .5 .5])
ylabel('Var_c_o_l P<0.05')

s = subplot(5,3,8);
hold(s,'on');
set(s,'FontName','Times New Roman','FontSize',16,'LineWidth',1.5,'XTick',[1 2 3],'XTickLabel',{'1','2','S_r_e_g'});
plot(g_levels,sum(s1(rr,cc).sg_var_col_pval<=0.05,3)./sum(~isnan(s1(rr,cc).sg_var_col),3),'-o','LineWidth',1.5)
line(g_levels,[0.05 0.05 0.05 ; 0.075 0.075 0.075 ; 0.1 0.1 0.1]','LineStyle','--','Color',[.5 .5 .5])

s = subplot(5,3,9);
hold(s,'on');
set(s,'FontName','Times New Roman','FontSize',16,'LineWidth',1.5);
plot(tau_levels,sum(s1(rr,cc).st_var_col_pval<=0.05,3)./sum(~isnan(s1(rr,cc).st_var_col),3),'-o','LineWidth',1.5)
line(tau_levels,[0.05 0.05 0.05 ; 0.075 0.075 0.075 ; 0.1 0.1 0.1]','LineStyle','--','Color',[.5 .5 .5])

%plot for var_ex:

s = subplot(5,3,10);
hold(s,'on');
set(s,'FontName','Times New Roman','FontSize',16,'LineWidth',1.5);
plot(c_levels,sum(s1(rr,cc).sc_var_ex_pval<=0.05,3)./sum(~isnan(s1(rr,cc).sc_var_ex),3),'-o','LineWidth',1.5)
line(c_levels,[0.05 0.05 0.05 0.05 ; 0.075 0.075 0.075 0.075 ; 0.1 0.1 0.1 0.1],'LineStyle','--','Color',[.5 .5 .5])
ylabel('Var_e_x P<0.05')

s = subplot(5,3,11);
hold(s,'on');
set(s,'FontName','Times New Roman','FontSize',16,'LineWidth',1.5,'XTick',[1 2 3],'XTickLabel',{'1','2','S_r_e_g'});
plot(g_levels,sum(s1(rr,cc).sg_var_ex_pval<=0.05,3)./sum(~isnan(s1(rr,cc).sg_var_ex),3),'-o','LineWidth',1.5)
line(g_levels,[0.05 0.05 0.05 ; 0.075 0.075 0.075 ; 0.1 0.1 0.1]','LineStyle','--','Color',[.5 .5 .5])

s = subplot(5,3,12);
hold(s,'on');
set(s,'FontName','Times New Roman','FontSize',16,'LineWidth',1.5);
plot(tau_levels,sum(s1(rr,cc).st_var_ex_pval<=0.05,3)./sum(~isnan(s1(rr,cc).st_var_ex),3),'-o','LineWidth',1.5)
line(tau_levels,[0.05 0.05 0.05 ; 0.075 0.075 0.075 ; 0.1 0.1 0.1]','LineStyle','--','Color',[.5 .5 .5])

%plot for c mean:

s = subplot(5,3,13);
hold(s,'on');
set(s,'FontName','Times New Roman','FontSize',16,'LineWidth',1.5);
plot(c_levels,sum(s1(rr,cc).sc_c_mean_pval<=0.05,3)./sum(~isnan(s1(rr,cc).sc_c_mean),3),'-o','LineWidth',1.5)
line(c_levels,[0.05 0.05 0.05 0.05 ; 0.075 0.075 0.075 0.075 ; 0.1 0.1 0.1 0.1],'LineStyle','--','Color',[.5 .5 .5])
xlabel('c')
ylabel('Avg. cov. P<0.05')

s = subplot(5,3,14);
hold(s,'on');
set(s,'FontName','Times New Roman','FontSize',16,'LineWidth',1.5,'XTick',[1 2 3],'XTickLabel',{'1','2','S_r_e_g'});
plot(g_levels,sum(s1(rr,cc).sg_c_mean_pval<=0.05,3)./sum(~isnan(s1(rr,cc).sg_c_mean),3),'-o','LineWidth',1.5)
line(g_levels,[0.05 0.05 0.05 ; 0.075 0.075 0.075 ; 0.1 0.1 0.1]','LineStyle','--','Color',[.5 .5 .5])
xlabel('g')

s = subplot(5,3,15);
hold(s,'on');
set(s,'FontName','Times New Roman','FontSize',16,'LineWidth',1.5);
plot(tau_levels,sum(s1(rr,cc).st_c_mean_pval<=0.05,3)./sum(~isnan(s1(rr,cc).st_c_mean),3),'-o','LineWidth',1.5)
line(tau_levels,[0.05 0.05 0.05 ; 0.075 0.075 0.075 ; 0.1 0.1 0.1]','LineStyle','--','Color',[.5 .5 .5])
xlabel('\tau')
legend({'S_r_e_g = 30', 'S_r_e_g = 90', 'S_r_e_g = 270'},'Orientation','horizontal')
```
