## Appendix S3 for "A framework for quantifying deviations from dynamic equilibrium theory": README.docx

Code last tested: 31 July 2019	Written by: Michael Kalyuzhny

### How to use the code in this appendix?

The code in this appendix should allow reproducing all the results, and enable the applicability of the methodology for other studies. The code is written in the Matlab™.

The simplest example for using the methodology for testing Dynamic Equilibrium (DE) is given in ‘EXAMPLE.m’. This self-explanatory code simulates DE using the ‘MWDR_2’ function and then computes and tests the five statistics using the ‘test_all4’ function. This is what you need to use for practical application of the methodology. If you wish to reproduce the results, continue reading.

The major results of the work are generated using script files that begin with ‘runs_’. These files generate the results while the corresponding ‘plot_’ file generates the relevant graphical representation. Please note that to expediate computation, parallel for loops have been used and the most time-consuming procedures (generating data and testing) have been transformed to MEX files. The original functions are still available, and to use them just remove the suffix ‘_mex’ from the relevant functions. Please note that even with these time-saving tricks, it might take a few days-a week to generate all results on a desktop computer as of 2019.

Please note that ‘ds’ refers here to k = 0 (taking all data into account) while ‘dsc’ considers the given k parameter.

While many of the files in this folder are auxiliary functions, the following files are used to generate the main results:

- ‘runs_type1.m’, ‘runs_type1_false.m’, ‘runs_type1_holes.m’, ‘runs_type1_incomplete.m’ launches the analysis of type 1 errors, with and without issues with the data.
- ‘runs_type2.mat’ launches the analysis of statistical power under the SDE model.

The following files generate more (minor) results and plot results:

- ‘plot_power_ratio_VMR.m’ plots the ration of statistical power of our methodology and the VMR method (Figure S13-S14).
- ‘plot_type1_false2.m’ plots the figures of the standardized effect size and significance for the case of false detection (Figure S4-S5).
- ‘plot_type1_holes.m’ plots the figures of the standardized effect size and significance for the case of missing years (Figure S8-S9).
- ‘plot_type1_incomplete2.m’ plots the figures of the standardized effect size and significance for the case of incomplete detection (Figure S6-S7).
- ‘plot_type1_ver3.m’ plots the significance and effect size of the statistics (Figure 3, Figure S3
- ‘plot_type2_SDE_ver2.m’ generates the figures of statistical power under the alternative SDE model.
- ‘plot_type1_OLS’ generates a plot of type I errors tested using OLS regression (Figure S15)
- ‘plot_type1_ver2_ML’ generates plots of type I errors tested DE simulations with ML estimated rates as reference (Figures S11-S12).
- ‘plot_type1_dornelas.m’ generates the code for the type 1 errors with the method of Dornelas *et al.* (2014, Figure S16)
- ‘runs_bias.m’ performs the bias calculation under the MLE method and generates the corresponding figure (Figure S10).
- ‘simberloff_analysis_final.m’ performs all the analysis of the Florida Keys experiment (Figure 4, table 2).
