## Appendix S3 for "A framework for quantifying deviations from dynamic equilibrium theory": runs_bias.m

```
%compute the bias in the estimate of extinction rates. assume that the
%species is initially present and that the rate of immigration is fixed.
%calculation is based on running a simulation of DE with number of species
%as in the number of itterations + 1 which is always absent (otherwise the
%algorithm crashes)

%IMPORTANT: running with 80 years and 10^6 itterations requires ~20GB RAM.

len_ts_levels = [10 20 40 80]; %durations of the time series
rates = .01:.01:.99; %levels of rates of extinction
ittr_per_rate = 10000; %itterations
im_now = .1; %the othe rate (immigration)
%bias0 = zeros(length(len_ts_levels),length(rates)); %results
bias1 = zeros(length(len_ts_levels),length(rates)); %results

%{
%initial: 0
for ll = 1:length(len_ts_levels)
    parfor rr = 1:length(rates)
        [ll rr]
        %biases = zeros(1,ittr_per_rate);
        dyn = MWDR_2_mex([im_now*ones(ittr_per_rate,1); 0], [rates(rr)*ones(ittr_per_rate,1); 0], [zeros(ittr_per_rate,1); 1],1,len_ts_levels(ll)); %run simulation
        [~,ex_est] = estimate_rates1(dyn);
        ex_est(sum(dyn(:,1:end-1)) == 0) = NaN; %if species was not seen - do not consider it!
        average_est = nanmean(ex_est(1:end-1));

        bias0(ll,rr) = (average_est - rates(rr))/rates(rr);
        
    end
end
%}

%initial: 1

for ll = 1:length(len_ts_levels)
    parfor rr = 1:length(rates)
        [ll rr]
        %biases = zeros(1,ittr_per_rate);
        dyn = MWDR_2_mex([im_now*ones(ittr_per_rate,1); 0], [rates(rr)*ones(ittr_per_rate,1); 0], [ones(ittr_per_rate,1); 0],1,len_ts_levels(ll)); %run simulation
        [~,ex_est] = estimate_rates1(dyn);
        average_est = mean(ex_est(1:end-1));
        bias1(ll,rr) = (average_est - rates(rr))/rates(rr);
        
    end
end

save('bias.mat')

%% plot results - old

f = figure();

axes1 = axes('Parent',f);
hold on
hold(axes1,'on');
set(axes1,'FontName','Times New Roman','FontSize',20,'LineWidth',2);
plot(rates,100*bias1,'-','LineWidth',1.5,'MarkerSize',4)
%plot(rates,100*bias0,'-o','LineWidth',1.5,'MarkerSize',4)
xlabel('Probability')
ylabel('Relative bias (%)')
legend({'Length = 10', 'Length = 20', 'Length = 40', 'Length = 80'})
```
