## Appendix S3 for "A framework for quantifying deviations from dynamic equilibrium theory": runs_type1.m

```
tic
sr_reg_levels=[30 90 270]; %regional richness
rates_levels=[0.4 0.4;0.4 0.04;0.04 0.4;0.04 0.04]; %each row is one regime. first column is colonization and the second is extinction. the numbers are both the mean and the variance.
t_max=40; %time series length
resamps=500;
ittr=5000;
cut_percent = 0.06; %percent of the time series to "cut" on both sides"

s1=struct;

global_null = generate_global_null(); %for the Dornelas methodology

%s1 has the following dimensions: 1: levels of rates (4). 2: levels of
%correlations (2)

sl=length(sr_reg_levels);

for rr=1:length(rates_levels(:,1))
    for cc=1:2 %levels of correlation between rates
        
        %arrays of the statistics that will be assigned after the parfor
        %loop:
        s1(rr,cc).dj=zeros(sl,ittr);
        s1(rr,cc).dj_pval=zeros(sl,ittr);
        s1(rr,cc).ds=zeros(sl,ittr);
        s1(rr,cc).ds_pval=zeros(sl,ittr);
        s1(rr,cc).dsc=zeros(sl,ittr);
        s1(rr,cc).dsc_pval=zeros(sl,ittr);
        s1(rr,cc).c_mean=zeros(sl,ittr);
        s1(rr,cc).c_mean_pval=zeros(sl,ittr);
        s1(rr,cc).var_ex=zeros(sl,ittr);
        s1(rr,cc).var_ex_pval=zeros(sl,ittr);
        s1(rr,cc).var_col=zeros(sl,ittr);
        s1(rr,cc).var_col_pval=zeros(sl,ittr);
        
        %statistics for the ML analysis:
        s1(rr,cc).dj_ML=zeros(sl,ittr);
        s1(rr,cc).dj_ML_pval=zeros(sl,ittr);
        s1(rr,cc).ds_ML=zeros(sl,ittr);
        s1(rr,cc).ds_ML_pval=zeros(sl,ittr);
        s1(rr,cc).dsc_ML=zeros(sl,ittr);
        s1(rr,cc).dsc_ML_pval=zeros(sl,ittr);
        s1(rr,cc).c_mean_ML=zeros(sl,ittr);
        s1(rr,cc).c_mean_ML_pval=zeros(sl,ittr);
        s1(rr,cc).var_ex_ML=zeros(sl,ittr);
        s1(rr,cc).var_ex_ML_pval=zeros(sl,ittr);
        s1(rr,cc).var_col_ML=zeros(sl,ittr);
        s1(rr,cc).var_col_ML_pval=zeros(sl,ittr);
        
        s1(rr,cc).VM=zeros(sl,ittr);
        s1(rr,cc).OLS_pval=zeros(sl,ittr); 
        s1(rr,cc).Dornelas_pval=zeros(sl,ittr); 
            
        for ss=1:length(sr_reg_levels) %itterate over levels of regional richness
            [rr cc ss]
            s1(rr,cc).m1_e=rates_levels(rr,2); %mean rate of extinction
            s1(rr,cc).v1_e=rates_levels(rr,2); %var rate of extinction
            s1(rr,cc).m1_c=rates_levels(rr,1); %mean rate of colonization
            s1(rr,cc).v1_c=rates_levels(rr,1); %var rate of colonization
            
            %transform the rates:
            mx1_e=log((s1(rr,cc).m1_e^2)/sqrt(s1(rr,cc).v1_e+s1(rr,cc).m1_e^2));
            vx1_e=sqrt(log(s1(rr,cc).v1_e/(s1(rr,cc).m1_e^2)+1));
            mx1_c=log((s1(rr,cc).m1_c^2)/sqrt(s1(rr,cc).v1_c+s1(rr,cc).m1_c^2));
            vx1_c=sqrt(log(s1(rr,cc).v1_c/(s1(rr,cc).m1_c^2)+1));
            
            %prepare the arrays for this round of the parfor loop:
            dj=zeros(1,ittr);
            dj_pval=zeros(1,ittr);
            ds=zeros(1,ittr);
            ds_pval=zeros(1,ittr);
            dsc=zeros(1,ittr);
            dsc_pval=zeros(1,ittr);
            c_mean=zeros(1,ittr);
            c_mean_pval=zeros(1,ittr);
            var_ex=zeros(1,ittr);
            var_ex_pval=zeros(1,ittr);
            var_col=zeros(1,ittr);
            var_col_pval=zeros(1,ittr);
            
            %statistics for the ML:
            dj_ML=zeros(1,ittr);
            dj_ML_pval=zeros(1,ittr);
            ds_ML=zeros(1,ittr);
            ds_ML_pval=zeros(1,ittr);
            dsc_ML=zeros(1,ittr);
            dsc_ML_pval=zeros(1,ittr);
            c_mean_ML=zeros(1,ittr);
            c_mean_ML_pval=zeros(1,ittr);
            var_ex_ML=zeros(1,ittr);
            var_ex_ML_pval=zeros(1,ittr);
            var_col_ML=zeros(1,ittr);
            var_col_ML_pval=zeros(1,ittr);
            
            VM = zeros(1,ittr);
            OLS_pval = zeros(1,ittr);
            Dornelas_pval = zeros(1,ittr);
            
            parfor ii=1:ittr
                
                % maker sure there are at least one time series with a
                % presence_absence_presence sequence and one with
                % absence_presence_absence
                dyn=0;
                sdd=0; %numer of events in a time series
                
                while (sum((sdd>=2)&(dyn(1,:)==0))<1)||(sum((sdd>=2)&(dyn(1,:)==1))<1)
                    
                    im_rates=sort(lognrnd(mx1_c,vx1_c,sr_reg_levels(ss),1)); %generate immigration rates
                    %extinction rates
                    if cc==1 %negative correlations with colonization
                        ex_rates=sort(lognrnd(mx1_e,vx1_e,sr_reg_levels(ss),1),'descend'); %that way we have the same values but with full negative tradeoff
                    else %no correlation
                        ex_rates=lognrnd(mx1_e,vx1_e,sr_reg_levels(ss),1); 
                    end

                    %transform rates to probabilities:
                    im=1-exp(-im_rates);
                    ex=1-exp(-ex_rates);

                    isl_initial=binornd(1,im./(im+ex)); %initial composition
                    dyn=MWDR_2_mex(im,ex,isl_initial,1,t_max); %generate time_series
                    dd=diff(dyn);
                    sdd=sum(abs(dd)); %numer of events in a time series
                end
                
                %now perform the tests!!!
                [c_mean(ii),c_mean_pval(ii),var_ex(ii),var_ex_pval(ii),var_col(ii),var_col_pval(ii),ds(ii),ds_pval(ii),dsc(ii),dsc_pval(ii),dj(ii),dj_pval(ii),~,~,~,~,~,~]=test_all4_mex(dyn,resamps,cut_percent,false) %PARIS
                [c_mean_ML(ii),c_mean_ML_pval(ii),var_ex_ML(ii),var_ex_ML_pval(ii),var_col_ML(ii),var_col_ML_pval(ii),ds_ML(ii),ds_ML_pval(ii),dsc_ML(ii),dsc_ML_pval(ii),dj_ML(ii),dj_ML_pval(ii)]=test_all_ML_mex(dyn,resamps,cut_percent) %ML
                
                sr_ts = sum(dyn,2);
                VM(ii) = var(sr_ts)/mean(sr_ts); %VM
                
                %OLS:
                fit = fitlm(1:length(sr_ts),sr_ts);
                OLS_pval(ii) = fit.Coefficients.pValue(2)
                
                %calculate Dornelas-like slope of similarity thgouth time:
                J_sim_emp=1-J_init(dyn);
                f=polyfit(1:(length(sr_ts)-1),J_sim_emp(2:end),1);
                slope = f(1);
                Dornelas_pval(ii) = dist_pval(global_null,slope); 
            end
            
            %assign after parfor:
            
            s1(rr,cc).dj(ss,:)=dj;
            s1(rr,cc).dj_pval(ss,:)=dj_pval;
            s1(rr,cc).ds(ss,:)=ds;
            s1(rr,cc).ds_pval(ss,:)=ds_pval;
            s1(rr,cc).dsc(ss,:)=dsc;
            s1(rr,cc).dsc_pval(ss,:)=dsc_pval;
            s1(rr,cc).c_mean(ss,:)=c_mean;
            s1(rr,cc).c_mean_pval(ss,:)=c_mean_pval;
            s1(rr,cc).var_ex(ss,:)=var_ex;
            s1(rr,cc).var_ex_pval(ss,:)=var_ex_pval;
            s1(rr,cc).var_col(ss,:)=var_col;
            s1(rr,cc).var_col_pval(ss,:)=var_col_pval;
            
            %ML statistics:
            s1(rr,cc).dj_ML(ss,:)=dj_ML;
            s1(rr,cc).dj_ML_pval(ss,:)=dj_ML_pval;
            s1(rr,cc).ds_ML(ss,:)=ds_ML;
            s1(rr,cc).ds_ML_pval(ss,:)=ds_ML_pval;
            s1(rr,cc).dsc_ML(ss,:)=dsc_ML;
            s1(rr,cc).dsc_ML_pval(ss,:)=dsc_ML_pval;
            s1(rr,cc).c_mean_ML(ss,:)=c_mean_ML;
            s1(rr,cc).c_mean_ML_pval(ss,:)=c_mean_ML_pval;
            s1(rr,cc).var_ex_ML(ss,:)=var_ex_ML;
            s1(rr,cc).var_ex_ML_pval(ss,:)=var_ex_ML_pval;
            s1(rr,cc).var_col_ML(ss,:)=var_col_ML;
            s1(rr,cc).var_col_ML_pval(ss,:)=var_col_ML_pval; 
            
            s1(rr,cc).VM(ss,:)=VM; 
            s1(rr,cc).OLS_pval(ss,:)=OLS_pval;
            s1(rr,cc).Dornelas_pval(ss,:)=Dornelas_pval;
        
            save('type1_1.mat')
        end
    end
end
        
toc
```
