## Appendix S3 for "A framework for quantifying deviations from dynamic equilibrium theory": runs_type1_false.m

```
%in this version the time each species has a fixed probability to be
%spontaneously detected when absent (false detection) with probability
%given by err_levels

tic
sr_reg_levels=[30 90 270];
rates_levels=[0.4 0.4;0.4 0.04;0.04 0.4;0.04 0.04]; %each row is one regime. first column is colonization and the second is extinction. the numbers are both the mean and the variance.
err_levels=[1 0.99 0.97 0.9]; %levles of (inverse) false detection
t_max=40;
resamps=500;
ittr=5000;
cut_level = 0.06; %what proportion of the time series to cut on both sides for the calculation of dsrel?

s2=struct;

%s2 has the following dimensions: 1: levels of rates (4). 2: levels of
%correlations (2). 

sl=length(sr_reg_levels);
el=length(err_levels);

for rr=1:length(rates_levels(:,1))
    for cc=1:2
        
        s2(rr,cc).dj=zeros(sl,el,ittr);
        s2(rr,cc).dj_pval=zeros(sl,el,ittr);
        s2(rr,cc).ds=zeros(sl,el,ittr);
        s2(rr,cc).ds_pval=zeros(sl,el,ittr);
        s2(rr,cc).dsc=zeros(sl,el,ittr);
        s2(rr,cc).dsc_pval=zeros(sl,el,ittr);
        s2(rr,cc).c_mean=zeros(sl,el,ittr);
        s2(rr,cc).c_mean_pval=zeros(sl,el,ittr);
        s2(rr,cc).var_ex=zeros(sl,el,ittr);
        s2(rr,cc).var_ex_pval=zeros(sl,el,ittr);
        s2(rr,cc).var_col=zeros(sl,el,ittr);
        s2(rr,cc).var_col_pval=zeros(sl,el,ittr);    
        
        for ss=1:length(sr_reg_levels)
            
            s2(rr,cc).m1_e=rates_levels(rr,2); %mean rate of extinction
            s2(rr,cc).v1_e=rates_levels(rr,2); %var rate of extinction
            s2(rr,cc).m1_c=rates_levels(rr,1); %mean rate of colonization
            s2(rr,cc).v1_c=rates_levels(rr,1); %var rate of colonization
            
            %transform the rates:
            mx1_e=log((s2(rr,cc).m1_e^2)/sqrt(s2(rr,cc).v1_e+s2(rr,cc).m1_e^2));
            vx1_e=sqrt(log(s2(rr,cc).v1_e/(s2(rr,cc).m1_e^2)+1));
            mx1_c=log((s2(rr,cc).m1_c^2)/sqrt(s2(rr,cc).v1_c+s2(rr,cc).m1_c^2));
            vx1_c=sqrt(log(s2(rr,cc).v1_c/(s2(rr,cc).m1_c^2)+1));
            
            %prepare the arrays:
            
            dj=zeros(el,ittr);
            dj_pval=zeros(el,ittr);
            ds=zeros(el,ittr);
            ds_pval=zeros(el,ittr);
            dsc=zeros(el,ittr);
            dsc_pval=zeros(el,ittr);
            c_mean=zeros(el,ittr);
            c_mean_pval=zeros(el,ittr);
            var_ex=zeros(el,ittr);
            var_ex_pval=zeros(el,ittr);
            var_col=zeros(el,ittr);
            var_col_pval=zeros(el,ittr);

            for ii=1:ittr
                [rr cc ss ii]
                
                % maker sure there are at least one time series with a
                % presence_absence_presence sequence and one with
                % absence_presence_absence
                dyn=0;
                sdd=0; %numer of events in a time series
                
                while (sum((sdd>=2)&(dyn(1,:)==0))<1)||(sum((sdd>=2)&(dyn(1,:)==1))<1)
                    
                    im_rates=sort(lognrnd(mx1_c,vx1_c,sr_reg_levels(ss),1)); %generate immigration rates
                    %extinction rates
                    if cc==1 %negative correlations with colonization
                        ex_rates=sort(lognrnd(mx1_e,vx1_e,sr_reg_levels(ss),1),'descend'); %that way we have the same values but with full negative tradeoff
                    else %no correlation
                        ex_rates=lognrnd(mx1_e,vx1_e,sr_reg_levels(ss),1); 
                    end

                    %transform rates to probabilities:
                    im=1-exp(-im_rates);
                    ex=1-exp(-ex_rates);

                    isl_initial=binornd(1,im./(im+ex));
                    dyn=MWDR_2_mex(im,ex,isl_initial,1,t_max);
                    dd=diff(dyn);
                    sdd=sum(abs(dd)); %numer of events in a time series
                end

                inv_dyn=abs(dyn-1);
                
                parfor eee=1:length(err_levels) %now add the errors of false detection
                    dyn_sampled=binornd(inv_dyn,err_levels(eee)); %there is a probability to transform absence (now '1') into an presence (now '0')
                    dyn_sampled=abs(dyn_sampled-1); %back reversal of 1s and 0s
                    while sum(dyn_sampled(1,:))==0 %make sure there are species present in the first year
                        dyn_sampled=binornd(dyn,err_levels(eee));
                        dyn_sampled=abs(dyn_sampled-1);
                    end
                    
                    
                    %now perform the tests!!!
                    [c_mean(eee,ii),c_mean_pval(eee,ii),var_ex(eee,ii),var_ex_pval(eee,ii),var_col(eee,ii),var_col_pval(eee,ii),ds(eee,ii),ds_pval(eee,ii),dsc(eee,ii),dsc_pval(eee,ii),dj(eee,ii),dj_pval(eee,ii),~,~,~,~,~,~]=test_all4_mex(dyn_sampled,resamps,cut_level,false);
                    
                end
            end
            
            s2(rr,cc).dj(ss,:,:)=dj;
            s2(rr,cc).dj_pval(ss,:,:)=dj_pval;
            s2(rr,cc).ds(ss,:,:)=ds;
            s2(rr,cc).ds_pval(ss,:,:)=ds_pval;
            s2(rr,cc).dsc(ss,:,:)=dsc;
            s2(rr,cc).dsc_pval(ss,:,:)=dsc_pval;
            s2(rr,cc).c_mean(ss,:,:)=c_mean;
            s2(rr,cc).c_mean_pval(ss,:,:)=c_mean_pval;
            s2(rr,cc).var_ex(ss,:,:)=var_ex;
            s2(rr,cc).var_ex_pval(ss,:,:)=var_ex_pval;
            s2(rr,cc).var_col(ss,:,:)=var_col;
            s2(rr,cc).var_col_pval(ss,:,:)=var_col_pval;      
            
            save('type1_false.mat')
        end
    end
end
        
toc
```
