## Appendix S3 for "A framework for quantifying deviations from dynamic equilibrium theory": runs_type2.m

```
% in this version we included the dj2 and beta2 statistics and changed dabs
% to be the mean rather than the sum

tic
sr_reg_levels=[30 90 270];
rates_levels=[0.4 0.4;0.4 0.04;0.04 0.4;0.04 0.04]; %each row is one regime. first column is colonization and the second is extinction. the numbers are both the mean and the variance.
c_levels=[0 0.5 1 2];
g_levels=[1 2 3]; %3 is actually Sreg
tau_levels=[0.02 0.1 0.5];
t_max=40;
resamps=500;
ittr=5000;
cut_percent = 0.06; %percent of the time series to remove on both sides for the calculation of DSC

%s1=struct;

%s1 has the following dimensions: 1: levels of rates (4). 2: levels of
%correlations (2).

sl=length(sr_reg_levels);
cl=length(c_levels);
gl=length(g_levels);
tl=length(tau_levels);

for rr=1:length(rates_levels(:,1))
    for cc=1:2 %levels of correlation between rates
        
        s1(rr,cc).m1_e=rates_levels(rr,2); %mean rate of extinction
        s1(rr,cc).v1_e=rates_levels(rr,2); %var rate of extinction
        s1(rr,cc).m1_c=rates_levels(rr,1); %mean rate of colonization
        s1(rr,cc).v1_c=rates_levels(rr,1); %var rate of colonization

        %transform the rates:
        mx1_e=log((s1(rr,cc).m1_e^2)/sqrt(s1(rr,cc).v1_e+s1(rr,cc).m1_e^2));
        vx1_e=sqrt(log(s1(rr,cc).v1_e/(s1(rr,cc).m1_e^2)+1));
        mx1_c=log((s1(rr,cc).m1_c^2)/sqrt(s1(rr,cc).v1_c+s1(rr,cc).m1_c^2));
        vx1_c=sqrt(log(s1(rr,cc).v1_c/(s1(rr,cc).m1_c^2)+1));
        
        %%%%%%%%%%%%%%%%%%%%%%%%%%%%%%%%%%%%%%%%%
        %%%%%%%%%%%%%%%%%%%explore the S-C plane:
        %%%%%%%%%%%%%%%%%%%%%%%%%%%%%%%%%%%%%%%%%
        
        %arrays of the statistics that will be assigned after the parfor
        %loop:
        s1(rr,cc).sc_dj=zeros(cl,sl,ittr);
        s1(rr,cc).sc_dj_pval=zeros(cl,sl,ittr);
        s1(rr,cc).sc_ds=zeros(cl,sl,ittr);
        s1(rr,cc).sc_ds_pval=zeros(cl,sl,ittr);
        s1(rr,cc).sc_dsc=zeros(cl,sl,ittr);
        s1(rr,cc).sc_dsc_pval=zeros(cl,sl,ittr);
        s1(rr,cc).sc_c_mean=zeros(cl,sl,ittr);
        s1(rr,cc).sc_c_mean_pval=zeros(cl,sl,ittr);
        s1(rr,cc).sc_var_ex=zeros(cl,sl,ittr);
        s1(rr,cc).sc_var_ex_pval=zeros(cl,sl,ittr);
        s1(rr,cc).sc_var_col=zeros(cl,sl,ittr);
        s1(rr,cc).sc_var_col_pval=zeros(cl,sl,ittr);
        
        %for ML:
        s1(rr,cc).sc_dj_ML=zeros(cl,sl,ittr);
        s1(rr,cc).sc_dj_ML_pval=zeros(cl,sl,ittr);
        s1(rr,cc).sc_ds_ML=zeros(cl,sl,ittr);
        s1(rr,cc).sc_ds_ML_pval=zeros(cl,sl,ittr);
        s1(rr,cc).sc_dsc_ML=zeros(cl,sl,ittr);
        s1(rr,cc).sc_dsc_ML_pval=zeros(cl,sl,ittr);
        s1(rr,cc).sc_c_ML_mean=zeros(cl,sl,ittr);
        s1(rr,cc).sc_c_mean_ML_pval=zeros(cl,sl,ittr);
        s1(rr,cc).sc_var_ex_ML=zeros(cl,sl,ittr);
        s1(rr,cc).sc_var_ex_ML_pval=zeros(cl,sl,ittr);
        s1(rr,cc).sc_var_col_ML=zeros(cl,sl,ittr);
        s1(rr,cc).sc_var_col_ML_pval=zeros(cl,sl,ittr);
        
        s1(rr,cc).sc_VM=zeros(cl,sl,ittr); 
          
        for ss=1:length(sr_reg_levels)
            for ccc=1:length(c_levels)
                [rr cc 1 ss ccc]

                t_initial=5*1/tau_levels(2);

                %prepare the arrays for this round of the parfor loop:
                dj=zeros(1,ittr);
                dj_pval=zeros(1,ittr);
                ds=zeros(1,ittr);
                ds_pval=zeros(1,ittr);
                dsc=zeros(1,ittr);
                dsc_pval=zeros(1,ittr);
                c_mean=zeros(1,ittr);
                c_mean_pval=zeros(1,ittr);
                var_ex=zeros(1,ittr);
                var_ex_pval=zeros(1,ittr);
                var_col=zeros(1,ittr);
                var_col_pval=zeros(1,ittr);
                
                %for ML:
                dj_ML=zeros(1,ittr);
                dj_ML_pval=zeros(1,ittr);
                ds_ML=zeros(1,ittr);
                ds_ML_pval=zeros(1,ittr);
                dsc_ML=zeros(1,ittr);
                dsc_ML_pval=zeros(1,ittr);
                c_mean_ML=zeros(1,ittr);
                c_mean_ML_pval=zeros(1,ittr);
                var_ex_ML=zeros(1,ittr);
                var_ex_ML_pval=zeros(1,ittr);
                var_col_ML=zeros(1,ittr);
                var_col_ML_pval=zeros(1,ittr);
                
                VM=zeros(1,ittr);

                parfor ii=1:ittr
                    
                % maker sure there are at least one time series with a
                % presence_absence_presence sequence and one with
                % absence_presence_absence
                dyn=0;
                sdd=0; %numer of events in a time series
                    while (sum((sdd>=2)&(dyn(1,:)==0))<1)||(sum((sdd>=2)&(dyn(1,:)==1))<1)
                        im_rates=sort(lognrnd(mx1_c,vx1_c,sr_reg_levels(ss),1));
                        if cc==1 %negative correlations
                            ex_rates=sort(lognrnd(mx1_e,vx1_e,sr_reg_levels(ss),1),'descend'); %that way we have the same values but with full negative tradeoff
                        else %no correlation
                            ex_rates=lognrnd(mx1_e,vx1_e,sr_reg_levels(ss),1); 
                        end

                        im=1-exp(-im_rates);
                        ex=1-exp(-ex_rates);

                        isl_initial=binornd(1,im./(im+ex));
                        dyn=SDE_mex(im,ex,c_levels(ccc),g_levels(2),tau_levels(2),isl_initial,t_initial,t_initial+t_max-1)
                        dd=diff(dyn);
                        sdd=sum(abs(dd)); %numer of events in a time series
                    end

                    [c_mean(ii),c_mean_pval(ii),var_ex(ii),var_ex_pval(ii),var_col(ii),var_col_pval(ii),ds(ii),ds_pval(ii),dsc(ii),dsc_pval(ii),dj(ii),dj_pval(ii),~,~,~,~,~,~]=test_all4_mex(dyn,resamps,cut_percent,false) %PARIS
                    [c_mean_ML(ii),c_mean_ML_pval(ii),var_ex_ML(ii),var_ex_ML_pval(ii),var_col_ML(ii),var_col_ML_pval(ii),ds_ML(ii),ds_ML_pval(ii),dsc_ML(ii),dsc_ML_pval(ii),dj_ML(ii),dj_ML_pval(ii)]=test_all_ML_mex(dyn,resamps,cut_percent) %ML
                    
                    sr_ts = sum(dyn,2);
                    VM(ii) = var(sr_ts)/mean(sr_ts); %variance to mean ratio

                end
                
                
                s1(rr,cc).sc_dj(ccc,ss,:)=dj;
                s1(rr,cc).sc_dj_pval(ccc,ss,:)=dj_pval;
                s1(rr,cc).sc_ds(ccc,ss,:)=ds;
                s1(rr,cc).sc_ds_pval(ccc,ss,:)=ds_pval;
                s1(rr,cc).sc_dsc(ccc,ss,:)=dsc;
                s1(rr,cc).sc_dsc_pval(ccc,ss,:)=dsc_pval;
                s1(rr,cc).sc_c_mean(ccc,ss,:)=c_mean;
                s1(rr,cc).sc_c_mean_pval(ccc,ss,:)=c_mean_pval;
                s1(rr,cc).sc_var_ex(ccc,ss,:)=var_ex;
                s1(rr,cc).sc_var_ex_pval(ccc,ss,:)=var_ex_pval;
                s1(rr,cc).sc_var_col(ccc,ss,:)=var_col;
                s1(rr,cc).sc_var_col_pval(ccc,ss,:)=var_col_pval; 
                
                %ML:
                s1(rr,cc).sc_dj_ML(ccc,ss,:)=dj_ML;
                s1(rr,cc).sc_dj_ML_pval(ccc,ss,:)=dj_ML_pval;
                s1(rr,cc).sc_ds_ML(ccc,ss,:)=ds_ML;
                s1(rr,cc).sc_ds_ML_pval(ccc,ss,:)=ds_ML_pval;
                s1(rr,cc).sc_dsc_ML(ccc,ss,:)=dsc_ML;
                s1(rr,cc).sc_dsc_ML_pval(ccc,ss,:)=dsc_ML_pval;
                s1(rr,cc).sc_c_mean_ML(ccc,ss,:)=c_mean_ML;
                s1(rr,cc).sc_c_mean_ML_pval(ccc,ss,:)=c_mean_ML_pval;
                s1(rr,cc).sc_var_ex_ML(ccc,ss,:)=var_ex_ML;
                s1(rr,cc).sc_var_ex_ML_pval(ccc,ss,:)=var_ex_ML_pval;
                s1(rr,cc).sc_var_col_ML(ccc,ss,:)=var_col_ML;
                s1(rr,cc).sc_var_col_ML_pval(ccc,ss,:)=var_col_ML_pval; 
                
                s1(rr,cc).sc_VM(ccc,ss,:)=VM; %variance to mean ratio

                save(type2.mat)
            end
        end
        
        %%%%%%%%%%%%%%%%%%%%%%%%%%%%%%%%%%%
        %%%%%%%%%%%%%explore the S-G plane:
        %%%%%%%%%%%%%%%%%%%%%%%%%%%%%%%%%%%
        
        s1(rr,cc).sg_dj=zeros(gl,sl,ittr);
        s1(rr,cc).sg_dj_pval=zeros(gl,sl,ittr);
        s1(rr,cc).sg_ds=zeros(gl,sl,ittr);
        s1(rr,cc).sg_ds_pval=zeros(gl,sl,ittr);
        s1(rr,cc).sg_dsc=zeros(gl,sl,ittr);
        s1(rr,cc).sg_dsc_pval=zeros(gl,sl,ittr);
        s1(rr,cc).sg_c_mean=zeros(gl,sl,ittr);
        s1(rr,cc).sg_c_mean_pval=zeros(gl,sl,ittr);
        s1(rr,cc).sg_var_ex=zeros(gl,sl,ittr);
        s1(rr,cc).sg_var_ex_pval=zeros(gl,sl,ittr);
        s1(rr,cc).sg_var_col=zeros(gl,sl,ittr);
        s1(rr,cc).sg_var_col_pval=zeros(gl,sl,ittr);  
        
        %ML:
        s1(rr,cc).sg_dj_ML=zeros(gl,sl,ittr);
        s1(rr,cc).sg_dj_ML_pval=zeros(gl,sl,ittr);
        s1(rr,cc).sg_ds_ML=zeros(gl,sl,ittr);
        s1(rr,cc).sg_ds_ML_pval=zeros(gl,sl,ittr);
        s1(rr,cc).sg_dsc_ML=zeros(gl,sl,ittr);
        s1(rr,cc).sg_dsc_ML_pval=zeros(gl,sl,ittr);
        s1(rr,cc).sg_c_mean_ML=zeros(gl,sl,ittr);
        s1(rr,cc).sg_c_mean_ML_pval=zeros(gl,sl,ittr);
        s1(rr,cc).sg_var_ex_ML=zeros(gl,sl,ittr);
        s1(rr,cc).sg_var_ex_ML_pval=zeros(gl,sl,ittr);
        s1(rr,cc).sg_var_col_ML=zeros(gl,sl,ittr);
        s1(rr,cc).sg_var_col_ML_pval=zeros(gl,sl,ittr); 
        
        s1(rr,cc).sg_VM=zeros(gl,sl,ittr); %variance to mean ratio
        
        for ss=1:length(sr_reg_levels)
            for ggg=1:length(g_levels)
                [rr cc 2 ss ggg]

                t_initial=5*1/tau_levels(2);

                %prepare the arrays for this round of the parfor loop:
                dj=zeros(1,ittr);
                dj_pval=zeros(1,ittr);
                ds=zeros(1,ittr);
                ds_pval=zeros(1,ittr);
                dsc=zeros(1,ittr);
                dsc_pval=zeros(1,ittr);
                c_mean=zeros(1,ittr);
                c_mean_pval=zeros(1,ittr);
                var_ex=zeros(1,ittr);
                var_ex_pval=zeros(1,ittr);
                var_col=zeros(1,ittr);
                var_col_pval=zeros(1,ittr);
                
                %ML:
                dj_ML=zeros(1,ittr);
                dj_ML_pval=zeros(1,ittr);
                ds_ML=zeros(1,ittr);
                ds_ML_pval=zeros(1,ittr);
                dsc_ML=zeros(1,ittr);
                dsc_ML_pval=zeros(1,ittr);
                c_mean_ML=zeros(1,ittr);
                c_mean_ML_pval=zeros(1,ittr);
                var_ex_ML=zeros(1,ittr);
                var_ex_ML_pval=zeros(1,ittr);
                var_col_ML=zeros(1,ittr);
                var_col_ML_pval=zeros(1,ittr);
                
                VM=zeros(1,ittr);
                
                %correct the number of groups:
                groups=g_levels(ggg);
                if groups==3
                    groups=sr_reg_levels(ss);
                end

                parfor ii=1:ittr
                % maker sure there are at least one time series with a
                % presence_absence_presence sequence and one with
                % absence_presence_absence
                dyn=0;
                sdd=0; %numer of events in a time series
                    while (sum((sdd>=2)&(dyn(1,:)==0))<1)||(sum((sdd>=2)&(dyn(1,:)==1))<1)
                        im_rates=sort(lognrnd(mx1_c,vx1_c,sr_reg_levels(ss),1));
                        if cc==1 %negative correlations
                            ex_rates=sort(lognrnd(mx1_e,vx1_e,sr_reg_levels(ss),1),'descend'); %that way we have the same values but with full negative tradeoff
                        else %no correlation
                            ex_rates=lognrnd(mx1_e,vx1_e,sr_reg_levels(ss),1); 
                        end

                        im=1-exp(-im_rates);
                        ex=1-exp(-ex_rates);

                        isl_initial=binornd(1,im./(im+ex));
                        dyn=SDE_mex(im,ex,c_levels(3),groups,tau_levels(2),isl_initial,t_initial,t_initial+t_max-1)
                        
                        dd=diff(dyn);
                        sdd=sum(abs(dd)); %numer of events in a time series

                    end

                    [c_mean(ii),c_mean_pval(ii),var_ex(ii),var_ex_pval(ii),var_col(ii),var_col_pval(ii),ds(ii),ds_pval(ii),dsc(ii),dsc_pval(ii),dj(ii),dj_pval(ii),~,~,~,~,~,~]=test_all4_mex(dyn,resamps,cut_percent,false) %PARIS
                    [c_mean_ML(ii),c_mean_ML_pval(ii),var_ex_ML(ii),var_ex_ML_pval(ii),var_col_ML(ii),var_col_ML_pval(ii),ds_ML(ii),ds_ML_pval(ii),dsc_ML(ii),dsc_ML_pval(ii),dj_ML(ii),dj_ML_pval(ii)]=test_all_ML_mex(dyn,resamps,cut_percent) %ML
                    
                    sr_ts = sum(dyn,2);
                    VM(ii) = var(sr_ts)/mean(sr_ts); %variance to mean ratio
                    
                end

                s1(rr,cc).sg_dj(ggg,ss,:)=dj;
                s1(rr,cc).sg_dj_pval(ggg,ss,:)=dj_pval;
                s1(rr,cc).sg_ds(ggg,ss,:)=ds;
                s1(rr,cc).sg_ds_pval(ggg,ss,:)=ds_pval;
                s1(rr,cc).sg_dsc(ggg,ss,:)=dsc;
                s1(rr,cc).sg_dsc_pval(ggg,ss,:)=dsc_pval;
                s1(rr,cc).sg_c_mean(ggg,ss,:)=c_mean;
                s1(rr,cc).sg_c_mean_pval(ggg,ss,:)=c_mean_pval;
                s1(rr,cc).sg_var_ex(ggg,ss,:)=var_ex;
                s1(rr,cc).sg_var_ex_pval(ggg,ss,:)=var_ex_pval;
                s1(rr,cc).sg_var_col(ggg,ss,:)=var_col;
                s1(rr,cc).sg_var_col_pval(ggg,ss,:)=var_col_pval;
                
                %ML:
                s1(rr,cc).sg_dj_ML(ggg,ss,:)=dj_ML;
                s1(rr,cc).sg_dj_ML_pval(ggg,ss,:)=dj_ML_pval;
                s1(rr,cc).sg_ds_ML(ggg,ss,:)=ds_ML;
                s1(rr,cc).sg_ds_ML_pval(ggg,ss,:)=ds_ML_pval;
                s1(rr,cc).sg_dsc_ML(ggg,ss,:)=dsc_ML;
                s1(rr,cc).sg_dsc_ML_pval(ggg,ss,:)=dsc_ML_pval;
                s1(rr,cc).sg_c_mean_ML(ggg,ss,:)=c_mean_ML;
                s1(rr,cc).sg_c_mean_ML_pval(ggg,ss,:)=c_mean_ML_pval;
                s1(rr,cc).sg_var_ex_ML(ggg,ss,:)=var_ex_ML;
                s1(rr,cc).sg_var_ex_ML_pval(ggg,ss,:)=var_ex_ML_pval;
                s1(rr,cc).sg_var_col_ML(ggg,ss,:)=var_col_ML;
                s1(rr,cc).sg_var_col_ML_pval(ggg,ss,:)=var_col_ML_pval;
                
                s1(rr,cc).sg_VM(ggg,ss,:)=VM;

                save(type2.mat)
            end
        end
        
        %%%%%%%%%%%%%%%%%%%%%%%%%%%%%%%%%%%%%%%%%
        %%%%%%%%%%%%%%%%%explore the S-TAU plane:
        %%%%%%%%%%%%%%%%%%%%%%%%%%%%%%%%%%%%%%%%%
        
        s1(rr,cc).st_dj=zeros(tl,sl,ittr);
        s1(rr,cc).st_dj_pval=zeros(tl,sl,ittr);
        s1(rr,cc).st_ds=zeros(tl,sl,ittr);
        s1(rr,cc).st_ds_pval=zeros(tl,sl,ittr);
        s1(rr,cc).st_dsc=zeros(tl,sl,ittr);
        s1(rr,cc).st_dsc_pval=zeros(tl,sl,ittr);
        s1(rr,cc).st_c_mean=zeros(tl,sl,ittr);
        s1(rr,cc).st_c_mean_pval=zeros(tl,sl,ittr);
        s1(rr,cc).st_var_ex=zeros(tl,sl,ittr);
        s1(rr,cc).st_var_ex_pval=zeros(tl,sl,ittr);
        s1(rr,cc).st_var_col=zeros(tl,sl,ittr);
        s1(rr,cc).st_var_col_pval=zeros(tl,sl,ittr); 
        
        %ML:
        s1(rr,cc).st_dj_ML=zeros(tl,sl,ittr);
        s1(rr,cc).st_dj_ML_pval=zeros(tl,sl,ittr);
        s1(rr,cc).st_ds_ML=zeros(tl,sl,ittr);
        s1(rr,cc).st_ds_ML_pval=zeros(tl,sl,ittr);
        s1(rr,cc).st_dsc_ML=zeros(tl,sl,ittr);
        s1(rr,cc).st_dsc_ML_pval=zeros(tl,sl,ittr);
        s1(rr,cc).st_c_mean_ML=zeros(tl,sl,ittr);
        s1(rr,cc).st_c_mean_ML_pval=zeros(tl,sl,ittr);
        s1(rr,cc).st_var_ex_ML=zeros(tl,sl,ittr);
        s1(rr,cc).st_var_ex_ML_pval=zeros(tl,sl,ittr);
        s1(rr,cc).st_var_col_ML=zeros(tl,sl,ittr);
        s1(rr,cc).st_var_col_ML_pval=zeros(tl,sl,ittr);
        
        s1(rr,cc).st_VM=zeros(tl,sl,ittr);
        
        for ss=1:length(sr_reg_levels)
            for ttt=1:length(tau_levels)
                [rr cc 3 ss ttt]

                t_initial=5*1/tau_levels(ttt);

                %prepare the arrays for this round of the parfor loop:
                dj=zeros(1,ittr);
                dj_pval=zeros(1,ittr);
                ds=zeros(1,ittr);
                ds_pval=zeros(1,ittr);
                dsc=zeros(1,ittr);
                dsc_pval=zeros(1,ittr);
                c_mean=zeros(1,ittr);
                c_mean_pval=zeros(1,ittr);
                var_ex=zeros(1,ittr);
                var_ex_pval=zeros(1,ittr);
                var_col=zeros(1,ittr);
                var_col_pval=zeros(1,ittr);
                
                %ML:
                dj_ML=zeros(1,ittr);
                dj_ML_pval=zeros(1,ittr);
                ds_ML=zeros(1,ittr);
                ds_ML_pval=zeros(1,ittr);
                dsc_ML=zeros(1,ittr);
                dsc_ML_pval=zeros(1,ittr);
                c_mean_ML=zeros(1,ittr);
                c_mean_ML_pval=zeros(1,ittr);
                var_ex_ML=zeros(1,ittr);
                var_ex_ML_pval=zeros(1,ittr);
                var_col_ML=zeros(1,ittr);
                var_col_ML_pval=zeros(1,ittr);
                
                VM = zeros(1,ittr); %variance to mean ratio

                parfor ii=1:ittr
                % maker sure there are at least one time series with a
                % presence_absence_presence sequence and one with
                % absence_presence_absence
                dyn=0;
                sdd=0; %numer of events in a time series
                    while (sum((sdd>=2)&(dyn(1,:)==0))<1)||(sum((sdd>=2)&(dyn(1,:)==1))<1)
                        im_rates=sort(lognrnd(mx1_c,vx1_c,sr_reg_levels(ss),1));
                        if cc==1 %negative correlations
                            ex_rates=sort(lognrnd(mx1_e,vx1_e,sr_reg_levels(ss),1),'descend'); %that way we have the same values but with full negative tradeoff
                        else %no correlation
                            ex_rates=lognrnd(mx1_e,vx1_e,sr_reg_levels(ss),1); 
                        end

                        im=1-exp(-im_rates);
                        ex=1-exp(-ex_rates);

                        isl_initial=binornd(1,im./(im+ex));
                        dyn=SDE_mex(im,ex,c_levels(3),g_levels(2),tau_levels(ttt),isl_initial,t_initial,t_initial+t_max-1)
                        
                        dd=diff(dyn);
                        sdd=sum(abs(dd)); %numer of events in a time series

                    end
                    
                    [c_mean(ii),c_mean_pval(ii),var_ex(ii),var_ex_pval(ii),var_col(ii),var_col_pval(ii),ds(ii),ds_pval(ii),dsc(ii),dsc_pval(ii),dj(ii),dj_pval(ii),~,~,~,~,~,~]=test_all4_mex(dyn,resamps,cut_percent,false) %PARIS
                    [c_mean_ML(ii),c_mean_ML_pval(ii),var_ex_ML(ii),var_ex_ML_pval(ii),var_col_ML(ii),var_col_ML_pval(ii),ds_ML(ii),ds_ML_pval(ii),dsc_ML(ii),dsc_ML_pval(ii),dj_ML(ii),dj_ML_pval(ii)]=test_all_ML_mex(dyn,resamps,cut_percent) %ML
                    
                    sr_ts = sum(dyn,2);
                    VM(ii) = var(sr_ts)/mean(sr_ts); %variance to mean ratio
                    
                end
                
                s1(rr,cc).st_dj(ttt,ss,:)=dj;
                s1(rr,cc).st_dj_pval(ttt,ss,:)=dj_pval;
                s1(rr,cc).st_ds(ttt,ss,:)=ds;
                s1(rr,cc).st_ds_pval(ttt,ss,:)=ds_pval;
                s1(rr,cc).st_dsc(ttt,ss,:)=dsc;
                s1(rr,cc).st_dsc_pval(ttt,ss,:)=dsc_pval;
                s1(rr,cc).st_c_mean(ttt,ss,:)=c_mean;
                s1(rr,cc).st_c_mean_pval(ttt,ss,:)=c_mean_pval;
                s1(rr,cc).st_var_ex(ttt,ss,:)=var_ex;
                s1(rr,cc).st_var_ex_pval(ttt,ss,:)=var_ex_pval;
                s1(rr,cc).st_var_col(ttt,ss,:)=var_col;
                s1(rr,cc).st_var_col_pval(ttt,ss,:)=var_col_pval;
                
                %ML:
                s1(rr,cc).st_dj_ML(ttt,ss,:)=dj_ML;
                s1(rr,cc).st_dj_ML_pval(ttt,ss,:)=dj_ML_pval;
                s1(rr,cc).st_ds_ML(ttt,ss,:)=ds_ML;
                s1(rr,cc).st_ds_ML_pval(ttt,ss,:)=ds_ML_pval;
                s1(rr,cc).st_dsc_ML(ttt,ss,:)=dsc_ML;
                s1(rr,cc).st_dsc_ML_pval(ttt,ss,:)=dsc_ML_pval;
                s1(rr,cc).st_c_mean_ML(ttt,ss,:)=c_mean_ML;
                s1(rr,cc).st_c_mean_ML_pval(ttt,ss,:)=c_mean_ML_pval;
                s1(rr,cc).st_var_ex_ML(ttt,ss,:)=var_ex_ML;
                s1(rr,cc).st_var_ex_ML_pval(ttt,ss,:)=var_ex_ML_pval;
                s1(rr,cc).st_var_col_ML(ttt,ss,:)=var_col_ML;
                s1(rr,cc).st_var_col_ML_pval(ttt,ss,:)=var_col_ML_pval;
                
                s1(rr,cc).st_VM(ttt,ss,:)=VM;

                save(type2.mat)
            end
        end
         
    end
end
        
toc
```
