## Appendix S3 for "A framework for quantifying deviations from dynamic equilibrium theory": SDE.m

```
function [composition] = SDE(im,ex0,c,g,tau,isl_initial,t_initial,t_max)

%a discrete Presence-Dependent extinction model. im and ex0 are column vectors of immigration and extinction
%probabilities in thr basal state. 
% c is the multiplier of the extinction probabilities in the bad state, g
% is the number of groups, tau is the switch probability of each group per
% turn

S_reg=length(im);
isl=isl_initial; %generate initial island
composition=zeros(length(t_initial:t_max),S_reg);
composition(1,:)=isl;

%calculate the extinction proability in the bad state:
mu0=log(ex0./(1-ex0));
mu1=mu0+c;
ex1=exp(mu1)./(exp(mu1)+1); %

ex=[ex0  ex1]; 
cycles=floor(S_reg/g); %the number of species per group

for t=2:t_max
    
    %switch environment:
    to_switch=binornd(1,tau,g,1); %which groups switched?
    
    %for each species the ex matrix holds its current and alternative
    %extinction rates and switchess between them if needed.
    ind_switch=logical(repmat(to_switch,cycles,1)); %which species switched?
    ex_tmp=ex(ind_switch,1); %the temporary first column, only changed values
    ex(ind_switch,1)=ex(ind_switch,2);
    ex(ind_switch,2)=ex_tmp;
    
    %extinction, considering available species:
    f=find(isl); %indexes of available species
    to_ex=binornd(1,ex(f,1)',1,length(f));
    
    %colonization, considering non available species:
    f_comp=find(isl==0);
    to_im= binornd(1,im(f_comp)',1,length(f_comp));

    %perform processes:

    isl(f(logical(to_ex)))=0; %perform extinction using the indexes from f
    isl(f_comp(logical(to_im)))=1; %perform colonization
    
    if t>=t_initial %sample
        composition(t-t_initial+1,:)=isl;
    end

end
```
