## Appendix S3 for "A framework for quantifying deviations from dynamic equilibrium theory": simberloff_analysis_final.m

```
% pre processing:

load('raw.mat')

resamps = 20000;
cut = 0.06;
nans_add = [3 3 4 3 4 3]; %number of missing samples before final sample

for si = 1:6
    
    data(si).dyn_short = data(si).dyn(1:end-1,:); %remove last census

    %add missing years before last census:
    last_ind = length(data(si).dyn(:,1));
    data(si).dyn_long = data(si).dyn;
    data(si).dyn_long(last_ind +nans_add(si),:) = data(si).dyn_long(end,:);
    data(si).dyn_long(last_ind :(last_ind +nans_add(si)-1),:) = NaN;

    data(si).dyn_short(:,sum(data(si).dyn_short) == 0) = []; %remove empty columns    
    data(si).dyn_long(:,nansum(data(si).dyn_long) == 0) = []; %remove empty columns
    
end

%%

c_mean_export_short = nan(6,resamps);
var_ex_export_short = nan(6,resamps);
var_col_export_short = nan(6,resamps);
ds_export_short = nan(6,resamps);
dsc_export_short = nan(6,resamps);
dj_export_short = nan(6,resamps);

c_mean_export_long = nan(6,resamps);
var_ex_export_long = nan(6,resamps);
var_col_export_long = nan(6,resamps);
ds_export_long = nan(6,resamps);
dsc_export_long = nan(6,resamps);
dj_export_long = nan(6,resamps);

%analyze data:
for si = 1:6
    [data(si).c_mean_short,data(si).c_mean_short_pval,data(si).var_ex_short,data(si).var_ex_short_pval,data(si).var_col_short,data(si).var_col_short_pval,data(si).ds_short,data(si).pval_ds_short,data(si).dsc_short,data(si).pval_dsc_short,data(si).dj_short,data(si).pval_dj_short,c_mean_export_short(si,:),var_ex_export_short(si,:),var_col_export_short(si,:),ds_export_short(si,:),dsc_export_short(si,:),dj_export_short(si,:)] = test_all4_mex(data(si).dyn_short,resamps,cut,true);
    data(si).c_mean_short_corr = data(si).c_mean_short - mean(c_mean_export_short(si,:));
    
    [data(si).c_mean_long,data(si).c_mean_long_pval,data(si).var_ex_long,data(si).var_ex_long_pval,data(si).var_col_long,data(si).var_col_long_pval,data(si).ds_long,data(si).pval_ds_long,data(si).dsc_long,data(si).pval_dsc_long,data(si).dj_long,data(si).pval_dj_long,c_mean_export_long(si,:),var_ex_export_long(si,:),var_col_export_long(si,:),ds_export_long(si,:),dsc_export_long(si,:),dj_export_long(si,:)] = test_all4_mex(data(si).dyn_long,resamps,cut,true);
    data(si).c_mean_long_corr = data(si).c_mean_long - mean(c_mean_export_long(si,:));
end

% a test for all:

c_mean_short_pval_all = dist_pval(mean(c_mean_export_short,1),mean([data.c_mean_short]))
var_ex_short_pval_all = dist_pval(mean(var_ex_export_short,1),mean([data.var_ex_short]))
var_col_short_pval_all = dist_pval(mean(var_col_export_short,1),mean([data.var_col_short]))
dsc_short_pval_all = dist_pval(mean(dsc_export_short,1),mean([data.dsc_short]))
dj_short_pval_all = dist_pval(mean(dj_export_short,1),mean([data.dj_short]))
        
c_mean_long_pval_all = dist_pval(mean(c_mean_export_long,1),mean([data.c_mean_long]))
var_ex_long_pval_all = dist_pval(mean(var_ex_export_long,1),mean([data.var_ex_long]))
var_col_long_pval_all = dist_pval(mean(var_col_export_long,1),mean([data.var_col_long]))
dsc_long_pval_all = dist_pval(mean(dsc_export_long,1),mean([data.dsc_long]))
dj_long_pval_all = dist_pval(mean(dj_export_long,1),mean([data.dj_long]))

%% compute relative measures:

for si = 1:6
    data(si).dsc_short_rel = data(si).dsc_short/sum(nanmean(data(si).dyn_short));
    data(si).dsc_long_rel = data(si).dsc_long/sum(nanmean(data(si).dyn_long));
end

%% figures - long:
%{
figs = figure();
resamps = 100;

for si = 1:6
    [sr_real,sr_null_save] = export_curves1(data(si).dyn_long,resamps);

    t_tot = data(si).time;
    last_ind = length(data(si).dyn(:,1));
    t_tot(last_ind +nans_add(si)) = t_tot(end);
    t_tot(last_ind :(last_ind +nans_add(si)-1)) = NaN;
    
    subplot1 = subplot(2,3,si,'Parent',figs);
    hold(subplot1,'on');
    set(subplot1,'CLim',[1 2],'FontName','Cakibri','FontSize',18,'LineWidth',1.5);
    errorbar(t_tot,mean(sr_null_save),2*std(sr_null_save),'LineWidth',2)
    hold on
    plot(t_tot,sr_real,'-o','LineWidth',2,'MarkerSize',8)
    hold off
    title(['site: ' num2str(data(si).plot) ' ds_r_e_l: ' num2str(data(si).dsc_long_rel,3) ' P: ' num2str(data(si).pval_dsc_long)])
    
    if si == 1
        ylabel('Richness')
        legend('Null','Observed','FontSize',16)
    end
    
    if si == 5
        xlabel('Time (days)')
    end
end
%}    
%% figures - short - old:
%{
figs = figure()
resamps = 100;

for si = 1:6
    [sr_real,sr_null_save] = export_curves1(data(si).dyn_short,resamps);

    t_tot = data(si).time(1:end-1);

    subplot1 =subplot(2,3,si)
    errorbar(t_tot,mean(sr_null_save),2*std(sr_null_save))
    hold on
    plot(t_tot,sr_real,'-o')
    hold off
    title(['site: ' num2str(data(si).plot) ' dsc: ' num2str(data(si).dsc_short) ' P: ' num2str(data(si).pval_dsc_short)])
    xlabel('Time (days)')
    ylabel('Richness')
end
%}
%% figures - short - new:

figs = figure()
resamps = 20000;
xlims = [300 300 300 450 300 250];
ylims = [45 25 40 40 45 40];

for si = 1:6
    [sr_real,sr_null_save] = export_curves1(data(si).dyn_short,resamps);

    t_tot = data(si).time(1:end-1);
    sr_null_save = sort(sr_null_save); %sort SR in each sample
    
    
    subplot1 =subplot(2,3,si)
    hold on
    x_fill=[t_tot fliplr(t_tot)];
    y_fill=[sr_null_save(round(0.025*resamps),:) fliplr(sr_null_save(round(0.975*resamps),:))];
    fill(x_fill, y_fill , 1,'facecolor',[0.95 0.7 0],'edgecolor','none','facealpha', 0.5);
    plot(t_tot,mean(sr_null_save),'color',[0.8 0 1],'linewidth',1.5)
    plot(t_tot,sr_real,'-o','color','blue','linewidth',1.5)
    hold off
    title([num2str(data(si).plot) ') ds: ' num2str(round(data(si).dsc_short,2)) ' P: ' num2str(round(data(si).pval_dsc_short,3))])
    if si ==5
        xlabel('Time (days)')
    end
    if si == 1
        ylabel('Richness')
        legend({'95% of null' , 'Mean null' , 'Observed'})
    end
    xlim([0 xlims(si)]);
    ylim([0 ylims(si)]);

   
    set(subplot1,'FontName','Cakibri','FontSize',18,'LineWidth',1.5)

end

%%

save('Simberloff.mat')

%% single figure for all islands:
```
