## Appendix S3 for "A framework for quantifying deviations from dynamic equilibrium theory": test_all4.m

```
function [c_mean,c_mean_pval,var_ex,var_ex_pval,var_col,var_col_pval,ds,pval_ds,dsc,pval_dsc,dj,pval_dj,c_mean_export,var_ex_export,var_col_export,ds_export,dsc_export,dj_export]=test_all4(dyn,resamps,cut,export_res)

%performs 6 tests overall, of the assumptions and the predictions. For each
%test it returns the statistic and it's corresponding P value.
%in this version, dsc is calculated on a censored time series, where
%ceil(cut) years are removed on both edges. 'cut' is a proportion. ds is
%not censored, unlike dsc

%in dyn, each row is a year and column is a species. resamps is the number
%of resamples.

% in version three there is an option to export the resampled statistics,
% to be used in computing site level p value. the function gets the boolean
% input 'export_res' to indicate whether it should export the resamples. if
% it shouldn't, it returns 'nan'.

%in version 4 the tests of ds and dsc are two sided.

sps=length(dyn(1,:));
upper_triu=logical(triu(ones(sps),1));
t_tot=length(dyn(:,1));
cut_years = ceil(cut*t_tot);

%empirical Jaccard compared to the initial:
j_emp=J_init(dyn);

%empirical ds and dsc:
sr_real=sum(dyn,2);
sr_real2 = sr_real((cut_years+1):(end-cut_years));
dsc_emp=max(sr_real2)-min(sr_real2);
ds_emp=max(sr_real)-min(sr_real);

%empirical mean statistic:
c_mat=nancov(dyn);
c_vals=c_mat(upper_triu); %take upper right triangle, without main diagonal
c_mean=mean(c_vals);

%for var_ex:
vars_ex=nan(sps,1);
lens_ex=zeros(sps,1);
sp_take_ex=false(1,sps);
for ss=1:sps
    move_durs=p_durations3(dyn(:,ss)');
    lens_ex(ss)=length(move_durs);
    if lens_ex(ss)>1 %consider only species with more than one presence segment
        vars_ex(ss)=var(move_durs,1);
        sp_take_ex(ss)=true;
    end
end

ff_ex=find(sp_take_ex)';

%for var_col:
dyn2=abs(dyn-1);
vars_col=nan(sps,1);
lens_col=zeros(sps,1);
sp_take_col=false(1,sps);
for ss=1:sps
    move_durs=p_durations3(dyn2(:,ss)');
    lens_col(ss)=length(move_durs);
    if lens_col(ss)>1 %consider only species with more than one absence segment
        vars_col(ss)=var(move_durs,1);
        sp_take_col(ss)=true;
    end
end

ff_col=find(sp_take_col)';


%resample:
c_mean_res=zeros(1,resamps);
vars_ex_res=nan(sps,resamps);
vars_col_res=nan(sps,resamps);
ds_res=zeros(1,resamps);
dsc_res = zeros(1,resamps);
j_res=zeros(resamps,t_tot);

for i=1:resamps
    dyn_null2=null2_comm3(dyn);
    
    %for ds:
    sr_null=sum(dyn_null2,2);
    ds_res(i)=max(sr_null)-min(sr_null);
    
    %for dsc:
    sr_null2 = sr_null((cut_years+1):(end-cut_years));
    dsc_res(i)=max(sr_null2)-min(sr_null2);
    
    %for dj:
    j_res(i,:)=J_init(dyn_null2);

    %for c_mean
    c_mat_res=nancov(dyn_null2);
    c_vals_res=c_mat_res(upper_triu); %take upper right triangle, without main diagonal
    c_mean_res(i)=mean(c_vals_res);

    %for the var_ex statistic:
    for kk=1:length(ff_ex) %only for the species we wish to consider
        ss=ff_ex(kk);
        move_durs_res=p_durations3(dyn_null2(:,ss)');
        vars_ex_res(ss,i)=var(move_durs_res,1);
    end
    
    %for the var_col statistic:
    dyn_null3=abs(dyn_null2-1);
    for kk=1:length(ff_col) %only for the species we wish to consider
        ss=ff_col(kk);
        move_durs_res=p_durations3(dyn_null3(:,ss)');
        vars_col_res(ss,i)=var(move_durs_res,1);
    end
    
end

%%%%%%%%%final calculations:

c_mean_pval=dist_pval(c_mean_res,c_mean);

%calclate var_ex:
sigma_v_ex=std(vars_ex_res,1,2); %std per species
sigma_v_ex(sigma_v_ex<0.001)=0; %correct numeric errors;
mu_v_ex=mean(vars_ex_res,2); %mean per species

di_ex=(vars_ex-mu_v_ex)./sigma_v_ex; %per species standardized difference
di_ex(sigma_v_ex<0.0001)=NaN; %correct numeric errors
di_ex=di_ex(isfinite(di_ex)); 
var_ex=mean(di_ex);

%resample var_ex:
di_ex_res=(vars_ex_res-mu_v_ex*ones(1,resamps))./(sigma_v_ex*ones(1,resamps));
di_ex_res(~isfinite(di_ex_res))=NaN;
d_ex_res=nanmean(di_ex_res);

var_ex_pval=dist_pval(d_ex_res,var_ex);

%calclate var_col:
sigma_v_col=std(vars_col_res,1,2); %std per species
sigma_v_col(sigma_v_col<0.001)=0; 
mu_v_col=mean(vars_col_res,2); %mean per species

di_col=(vars_col-mu_v_col)./sigma_v_col; %per species standardized difference
di_col(sigma_v_col<0.0001)=NaN; %correct numeric errors
di_col=di_col(isfinite(di_col)); 
var_col=mean(di_col);

%resample var_col:
di_col_res=(vars_col_res-mu_v_col*ones(1,resamps))./(sigma_v_col*ones(1,resamps));
di_col_res(~isfinite(di_col_res))=NaN;
d_col_res=nanmean(di_col_res);

var_col_pval=dist_pval(d_col_res,var_col);

%ds:
ds=ds_emp-mean(ds_res);
%pval_ds=sum(ds_res>=ds_emp)/resamps; old version
pval_ds = dist_pval(ds_res-mean(ds_res),ds);

%dsc:
dsc=dsc_emp-mean(dsc_res);
%pval_dsc=sum(dsc_res>=dsc_emp)/resamps; old version
pval_dsc = dist_pval(dsc_res-mean(dsc_res),dsc);

%dj:
j_mean=mean(j_res);
dj_emp=max(j_emp-j_mean);

dj_res=j_res-ones(resamps,1)*j_mean; %the deviation of each resample from the mean Jaccard curve
dj_res=max(dj_res,[],2);

dj=dj_emp-mean(dj_res);
pval_dj=sum(dj_res>=dj_emp)/resamps;

%if indicated that the resamples should be exported:
if export_res == true
    c_mean_export = c_mean_res;
    var_ex_export = d_ex_res;
    var_col_export = d_col_res;
    ds_export = ds_res - mean(ds_res);
    dsc_export = dsc_res - mean(dsc_res);
    dj_export = (dj_res - mean(dj_res))';
else
    c_mean_export = NaN;
    var_ex_export = NaN;
    var_col_export = NaN;
    ds_export = NaN;
    dsc_export = NaN;
    dj_export = NaN;
end

end

    
```
