## Appendix S1 S2 for "A framework for quantifying deviations from dynamic equilibrium theory"

### Appendix S1 – Procedure for testing the methodology

#### General settings and tests of type I errors under DE

We first wish to test the methodological framework under DE and ensure that it does not detect spurious patterns by applying it to synthetic time series generated under DE. Despite being simple, DE actually has multiple degrees of freedom, namely the extinction probability $E_i$ and colonization probability $C_i$ for each of the $S_{reg}$ species in the pool. We now describe the specific scenarios we simulated. The general settings were also applied later to study statistical power and robustness.

First, it makes intuitive sense to assume that the rates of colonization and extinction follow a lognormal distribution. This is because species abundance (Preston 1948; Ulrich *et al.* 2010) and body mass (Hutchinson & MacArthur 1959; Brown *et al.* 1993) are often lognormally distributed. Hence, the rates for each species are drawn from a lognormal distribution. Given that the colonization or extinction rate of some species is λ, we calculate the probability of that event taking place between two years, $C_i$ or $E_i$ respectively, as $1 - e^{-\lambda}$.

To simplify, we considered communities with two levels of rates (either colonization or extinction):

1. High, with mean rate 0.4 and variance of rates 0.4. In this case, the corresponding mean probability ($E_i$ or $C_i$) was 0.26 and the median was 0.19. These probabilities resemble those observed in the North American Breeding Bird Survey, which we study empirically.
2. Low, with mean rate 0.04 and variance of rates 0.04. In this case, the corresponding mean probability ($E_i$ or $C_i$) was 0.03 and the median was 0.0078.

Overall, four combinations of rates were considered (high/low colonization and high/low extinction).

Another degree of freedom is the relationship between colonization and extinction probabilities. First, we consider a scenario of a negative tradeoff, indicating that species with low extinction rates would be those with high colonization rates and vice versa. This is a biologically likely scenario since such species are likely better adapted to the environment or simply have higher abundances. We also consider a scenario of no correlation between colonization and extinction rates.

We tested three levels of $S_{reg}$: 30, 90 and 270 species. Their rates were drawn from a lognormal distribution with the aforementioned parameters according to the defined levels (high/low) and transformed into probabilities. Next, in the negative correlation scenario, $C_i$s were assigned in ascending order and $E_i$s were assigned in descending order. These rates were drawn independently for each synthetic dataset.

To generate a synthetic dataset under DE, the community was initialized at a composition drawn from the steady state distribution (where the probability of species $i$ to occur is $\frac{C_i}{C_i+E_i}$) and the dynamics was simulated for 40 years. This is a relatively long but realistic time frame for ecological data, and we believe that taking a long period can help identify a tendency of the method to spuriously detect patterns (make type I errors). The PARIS framework was then applied to the synthetic data by generating 500 resamples and calculating the value and significance of the five statistics. In all analyses in this work we only considered datasets with at least one time series with presence-absence-presence periods and at least one time series with absence-presence-absence periods, that is, we do not consider datasets that cannot be resampled with the PARIS null model.

To study type I errors, we generated 5000 synthetic communities at each of the 24 scenarios (two levels of colonization rates, two levels of extinction rates, two levels of correlation between them and three levels of $S_{reg}$) and calculated the statistics for each. We present in Figure S1 the mean value of the five statistics and in figure 3 of the main text the proportion of significant results (the realized α).

#### Studying performance with an alternative model

The aim of this analysis is to show that the PARIS framework has reasonable power to detect deviations from the assumptions and predictions of DE. Further, we wish to study its performance in cases when only one of the assumptions (either species independence or rate uniformity) is violated and check whether the statistic testing the other assumption would have increased type I error rate. We are also interested to know the typical scales of effect sizes and their dependence on $S_{reg}$ and to make sure that they increase with the increase in the deviation from DE.

We addressed these questions using a novel model, the State-Dependent Extinction (hereafter SDE) model. This toy model of presence-absence dynamics represents a scenario when extinction depends on environmental drivers that fluctuate through time, and these drivers may be shared among species, facilitating a correlated response.

In the SDE model, each species can be in one of two states, basal and harsh, and the extinction probability is elevated in the latter. The elevated probability is obtained by adding a fixed number *c* (representing the degree of harshness of harsh years) to the extinction probability on the logit scale $\epsilon_i$, and back transformation. In other words, if $E_{i1}$ is the basic extinction probability of species *i*, it can be represented on the logit scale as:

(S1) $$\epsilon_{i1} = \text{logit}(E_{i1}) = \log\left(\frac{E_{i1}}{1-E_{i1}}\right).$$

The extinction probability under harsh conditions (on the logit scale), $\epsilon_{i2}$, is now obtained:

(S2) $$\epsilon_{i2} = \epsilon_{i1} + c.$$

From this, back transformation leads to the extinction probability of species *i* under harsh conditions on the arithmetic scale:

(S3) $$E_{i2} = logit^{-1}(\epsilon_{i2}) = \frac{e^{\epsilon_{i2}}}{e^{\epsilon_{i2}+1}}.$$

This 'mathematical back and forth' is used so that a fixed strength of environmental fluctuations *c* can be applied to all species despite the probabilities being different and bounded. If c = 0, the model is reduced to DE.

Importantly, species are divided into *g* fixed groups, and all the species in a group are at the same state and switch states together and independently of other groups. If g = 1, all the species respond in a similar manner (all are either at a harsh state or at the basal state simultaneously), and if g = $S_{reg}$ the species are independent. Hence, the model progresses like DE with rates according to the currents state, but after each time step, each group switches states with probability $\tau$, which represents the degree of temporal autocorrelation.

The features of this model allow breaking the assumptions of DE separately: If g = $S_{reg}$ but c > 0, species have fluctuating rates but are independent. On the other hand, if c > 0, $\tau$ = 0.5 and g < $S_{reg}$ species are correlated but they effectively have a uniform extinction rate.

Specifically, we used a default parameter regime of $\tau$ = 0.1, g = 2 and c = 1. We tested separately the effect of autocorrelation by simulating also $\tau$ = 0.02 and $\tau$ = 0.5, the effect of the number of groups by simulating also *g* = 1 and g = $S_{\mathrm{reg}}$ and the effect of fluctuation magnitude by simulating also c = 0, c = 0.5 and c = 2.

The procedure we used was largely similar to the one used to study type I errors. As before, we considered 24 levels of basal rates, correlations among rates and $S_{reg}$, but each such level was simulated also with the 3 levels of $\tau$, 3 levels of *g* and 4 levels of *c*. This resulted in 864 scenarios overall. Synthetic communities were initialized using the steady state distribution of the basal rates (as before) and were allowed $5/\tau$ time steps to equilibrate. For each scenario we generated 5000 synthetic communities and calculated the statistics and their significance.

#### Testing robustness

In dynamical analyses it is crucial to address the adverse and nontrivial effects of issues with the data. We considered the effect of three types of issues on the performance of the framework under DE (type I errors): a) incomplete detection, meaning that a species that is present is not detected; b) false detection, when species that are not present are detected; and c) missing years, when the entire composition in a given year is not recorded. The general settings here were identical to the testing under DE described above.

In the incomplete detection case, each year a species was present, there was a fixed probability $P_{miss}$ not to observe it. In the false detection scenario, each year a species was absent (including species that haven't arrived during the simulation), there was a probability $P_{false}$ to 'observe' it nonetheless. Finally, in the missing years scenario, *Y* years, excluding the first and the last year, were randomly chosen and the entire composition in these years was transformed into missing values.

Again, for each of the three effects and each of the 24 scenarios, 5000 synthetic datasets of 40 years under DE were generated, and subjected to either incomplete detection with $P_{miss}$ of 0, 0.1, 0.3 or 0.6, false detection with $P_{false}$ of 0, 0.01, 0.03 or 0.1 or number of removed years of 0, 1, 2, 4 or 8.

The results for a negative correlation between colonization and extinction are presented in Figures S4-S9. Results for no correlation are very similar. All results (for both correlation levels) are summarized in table 1 of the main text. For each of the five statistics we calculated the type I error rate and the standardized effect size (SES), defined as the average value of the statistic divided by the STD (calculated over the 5000 iterations). If SES is low, the typical effect size will be overwhelmed by the variability in the results. We then classified the 'sensitivity' of the statistic for a give $S_{reg}$ and level of issues in the data ($P_{miss}$, $P_{false}$ or number of removed years) in terms of both effect size and type I error rate in the following manner: For all eight combinations of rates (4) and correlations among them (2), we found the highest type I error rate (P) and the most extreme (high or low) effect size (SES). For this worst-case scenario, we defined no sensitivity (green, colors and symbols here refer to table 1) as $P < 0.075$; mild sensitivity (yellow) as $0.075 \leq P < 0.1$ and sensitivity (red) as $P \geq 0.1$. For SES, we defined no sensitivity (no symbol) as $|SES| < 0.1$; mild sensitivity (**X**) as $0.1 \leq |SES| < 0.2$ and sensitivity (**XX**) as $|SES| \geq 0.2$.

All simulations were performed in Matlab 2016a, with the full code provided in appendix S3.

#### Appendix S2 – Performance under an alternative model - results

Figure S1 and S2 show the behavior of the methodological framework, in terms of mean effect size and significance, under the SDE model in the case of high extinction and high colonization rates with a negative correlation between them. The results in the other cases are qualitatively very similar and are therefore not presented here. The main difference, naturally, is that at lower rates the power is (sometimes considerably) smaller.

First, it is noticeable that in all cases the values of $var_{col}$ are very small and their effective α is only sometimes higher than 0.05 (the maximal value we found is 0.088 for $S_{reg}$ = 270 species, high extinction, low colonization and negative correlation between them scenario). This indicated that the fact that extinction rates are not uniform and species are correlated can still have no serious impact on the rate uniformity of colonization. It can also be noticed that as c increases, the effect sizes and power increase as well. Since increasing *c* has the most straightforward effect of creating deviations from DE, this indicates that, indeed, the larger the deviations, the stronger and more detectable is the effect (with all statistics but $var_{col}$). Another trivial yet reassuring result is that power always increases with $S_{reg}$.

Our results also reveal a nice feature of $var_{ex}$ and $\overline{\text{cov}}$ – their effect size is not (or only slightly) affected by the regional number of species $S_{reg}$, indicating that they have an inherent universality property making them particularly valuable for comparison between communities. *dj* and *ds* (the latter not surprisingly!), on the other hand, are affected by regional richness.

When species are independent ($g = S_{reg}$), despite the rates being non-uniform, $\overline{\text{cov}}$ is effectively 0 and it's realized α is close to 0.05. This indicates that this statistic, testing the independence assumption, is robust to violations of the uniform rates assumption. Interestingly, *ds* and *dj* are also 0, indicating that non-uniform rates alone are not sufficient to generate large changes in richness or composition. These conclusions are valid only under the SDE model, of course. Another reassuring result is that the number of groups does not affect the mean value of $var_{ex}$.

Furthermore, when $\tau = 0.5$ and the rates are hence uniform, $var_{ex}$ is very close to 0 but the proportion of significant results here is slightly elevated, reaching 0.071 (for $S_{reg} = 270$) in Figure S3 and as much as ~0.085 in one of the scenarios that are not presented. This indicates that if species are correlated but rates are uniform, there is some increase in the type I error rate of detecting a non-uniform extinction rate, nevertheless. It is also worthwhile noting that both *ds* and *dj* in this case are elevated, indicating that correlation between species alone, even with uniform rates, is sufficient to generate large changes in richness and composition.

These results suggest that, overall, the methodological framework satisfies all our requirements. It has reasonable power (in the presented case of high rates), the values of the statistics and their significance increase with increasing deviations from the null, the tests of the assumptions have good specificity to the assumption they are testing and some of the statistics are independent of the number of species. It is noteworthy that, not surprisingly, the power can be much lower if colonization and extinction rates are low because not much dynamics is actually observed.

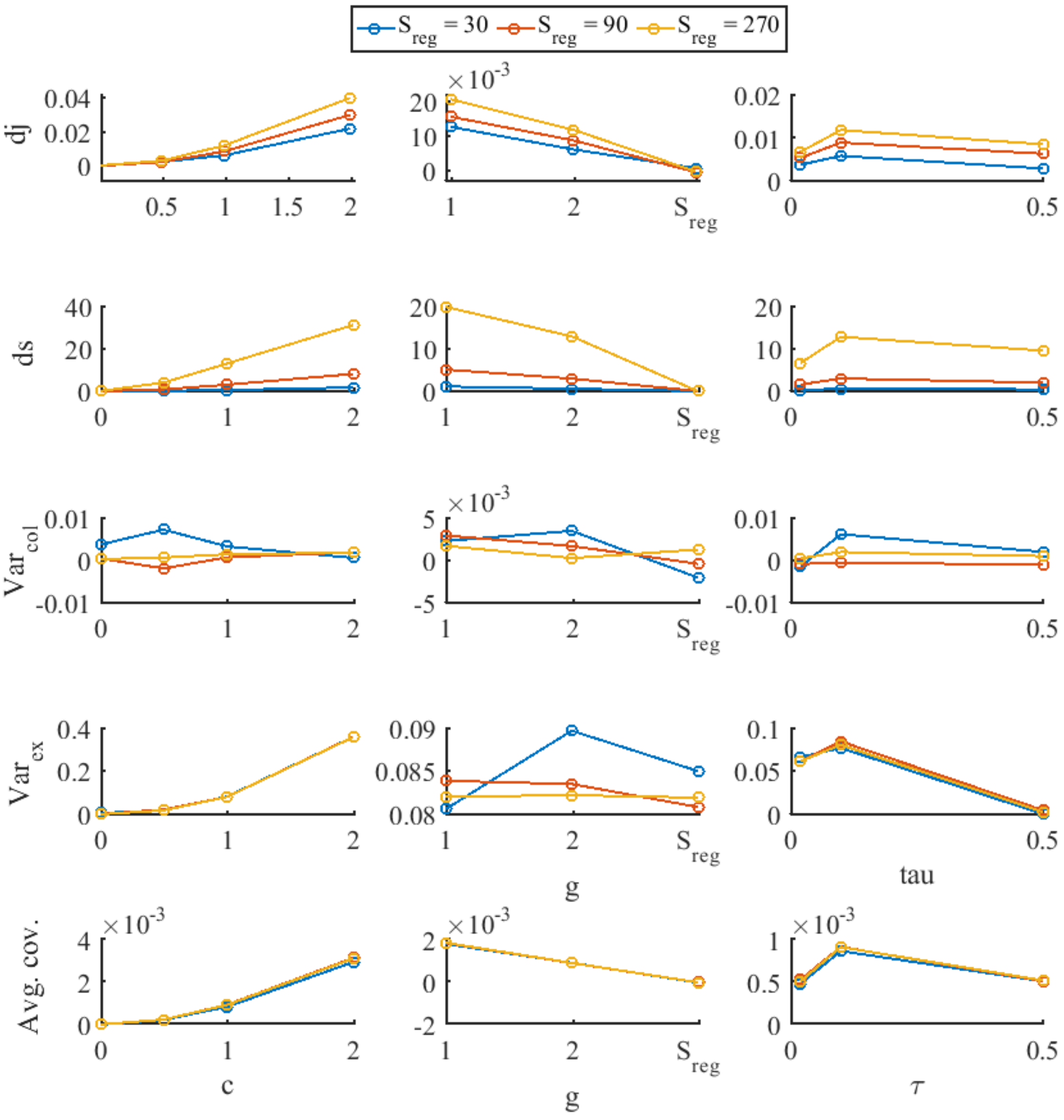


**Figure S1** – Mean value of the statistics under the alternative SDE model. Each row tests the performance of one statistic, while each column presents the effect of a different parameter (*c*, *g* or *τ*) on the mean value. Different colors represent different levels of $S_{reg}$. Here only the scenario of high colonization rates and high extinction rates with negative correlation between them is presented. The mean value is calculated over 5000 synthetic datasets.

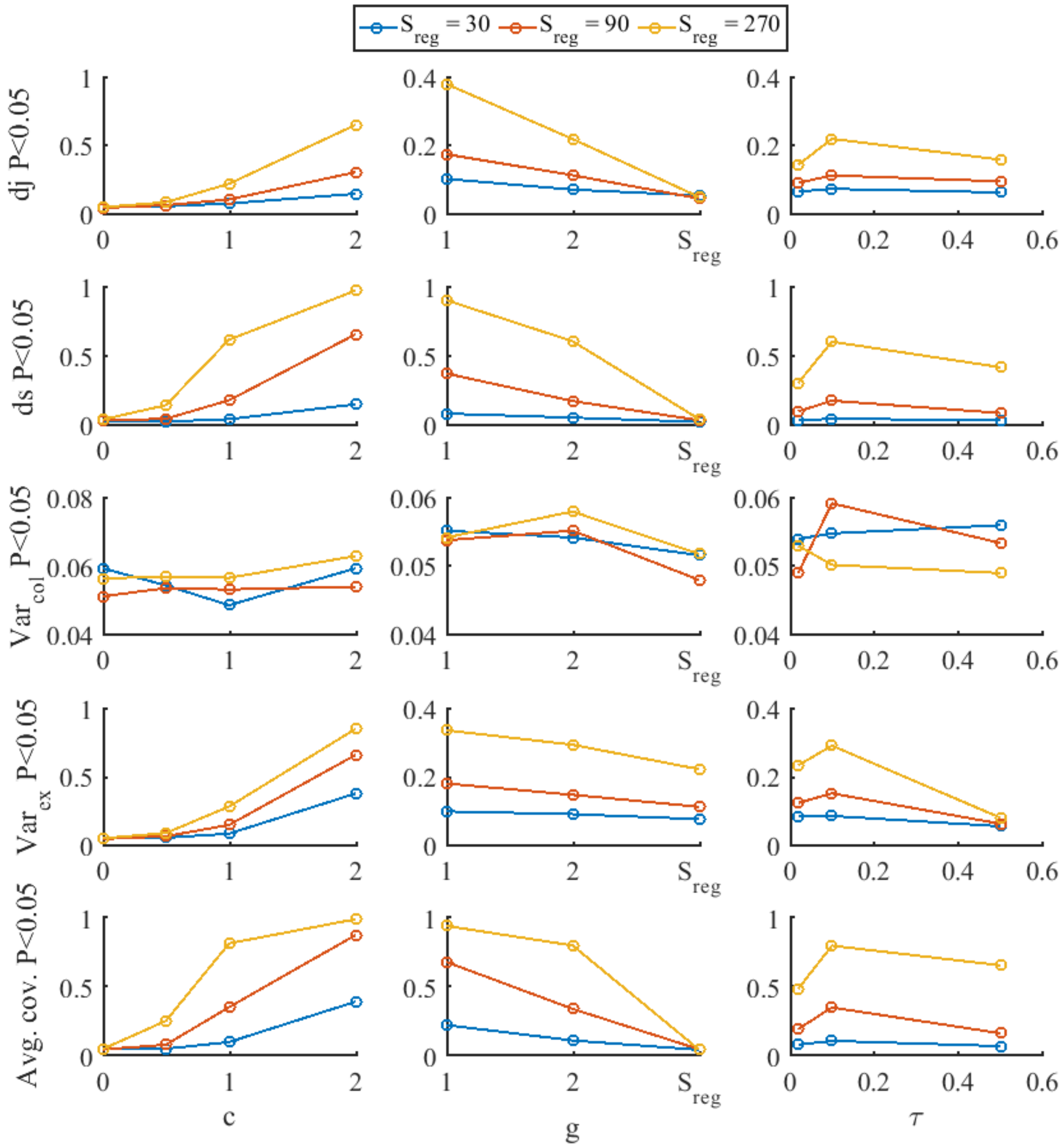


**Figure S2** – Significance of the statistics under the alternative SDE model. Each row presents the proportion of significant results (out of the 5000 synthetic datasets) for one statistic, which is the effective α of the test. Each column presents the effect of a different parameter (*c*, *g* or *τ*) on this α, and different colors represent different levels of $S_{reg}$. Here only the scenario of high colonization rates and high extinction rates with negative correlation between them is presented.

#### Additional supplementary figures

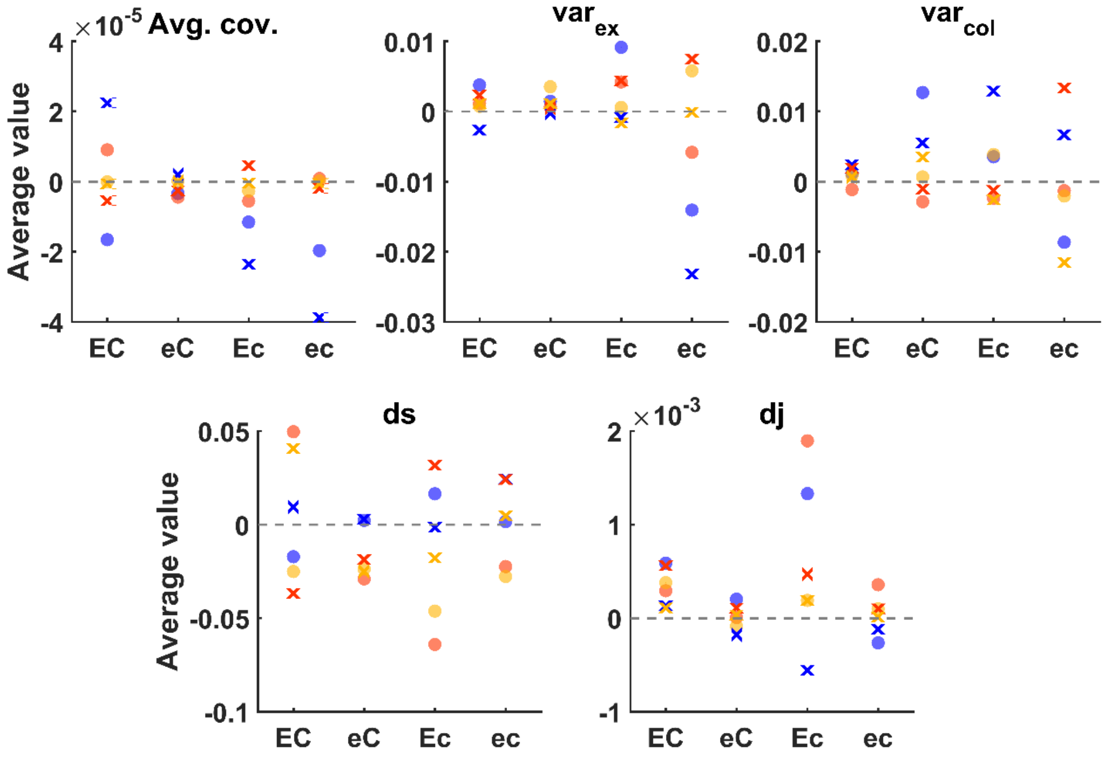


**Figure S3** – Average values of the summary statistics under DE at various parameter regimes. Four combinations of rates are considered – high and low average colonization (C and c) and high and low average extinction (E and e). simulations were run for each combination of rates, correlation between colonization and extinction ('o' = no correlation, 'x' = negative correlation) and size of the species pool (30 = blue, 90 = orange, 270 = yellow). Every point represents one of these combination, and the average statistics were calculated over 5000 synthetic communities. The grey dashed line represents a value of 0.

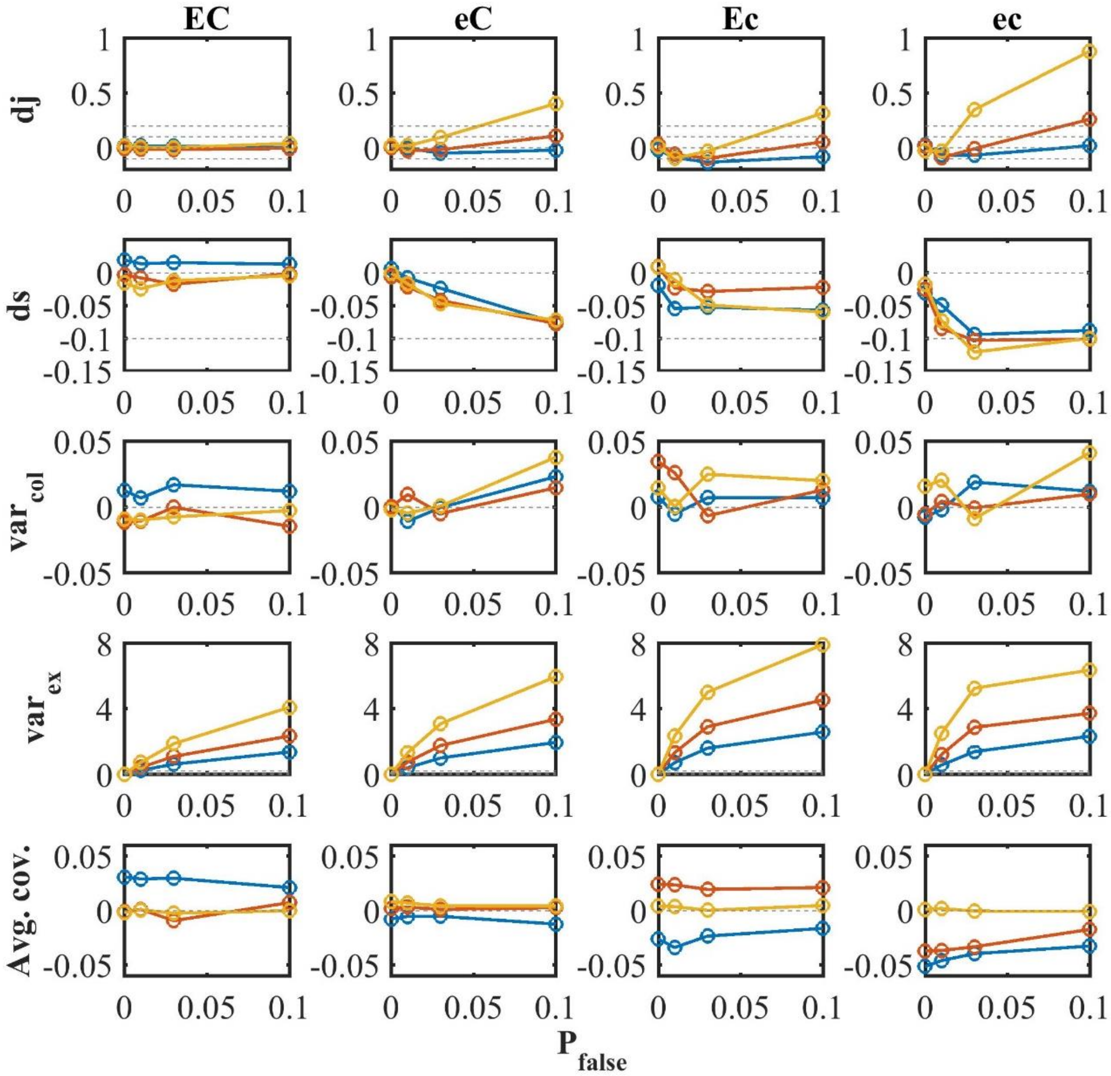


**Figure S4** – Standardized effect size of the statistics under DE, when each species in the pool has probability $P_{false}$ (x axis) to be detected each year when absent. Each row tests the performance of one statistic, while each column considers a different combination of rates (EC stands for high extinction and high colonization, eC stands for low extinction and high colonization and so forth). Different colors represent different levels of $S_{reg}$ (blue: $S_{reg}$ =30, red: $S_{reg}$ = 90, yellow: $S_{reg}$ = 270). Here only the scenario of negative correlation between colonization and extinction is presented. The standardized effect size is calculated as the average statistic over 5000 synthetic datasets divided by the SD. The grey horizontal lines highlight the values -0.1, 0, 0.1 and 0.2.

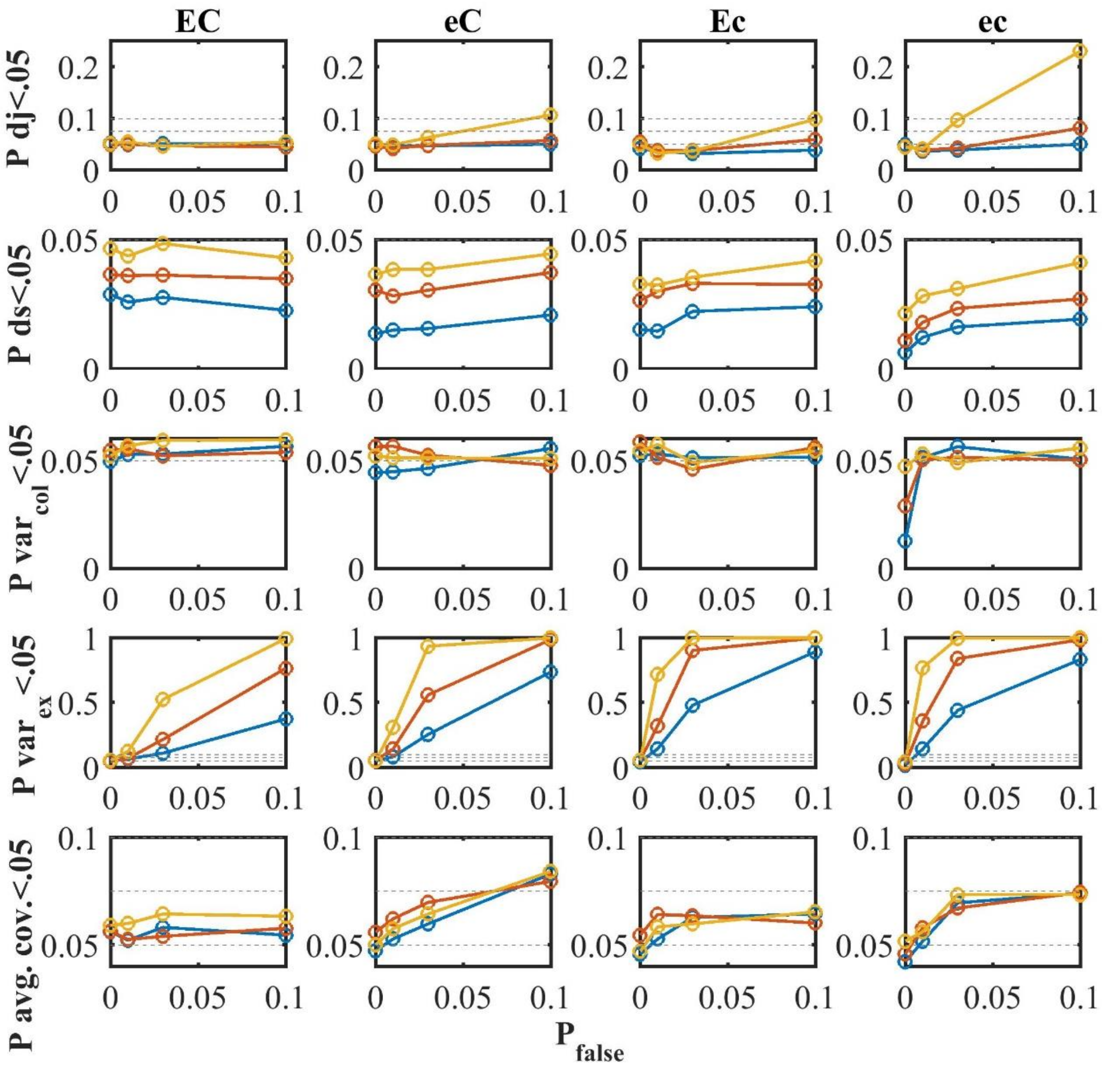


**Figure S5** – Significance of the statistics under DE, when each species in the pool has probability $P_{false}$ (x axis) to be detected each year when absent. Each row shows the performance of one statistic, presenting the proportion of significant results (out of the 5000 synthetic datasets), which is the effective α of the test. Each column considers a different combination of rates (EC stands for high extinction and high colonization, eC stands for low extinction and high colonization and so forth), while different colors represent different levels of $S_{reg}$ (blue: $S_{reg}$ =30, red: $S_{reg}$ = 90, yellow: $S_{reg}$ = 270). Here only the scenario of negative correlation between colonization and extinction is presented. The grey horizontal lines highlight the values of 0.05, 0.075 and 0.1.

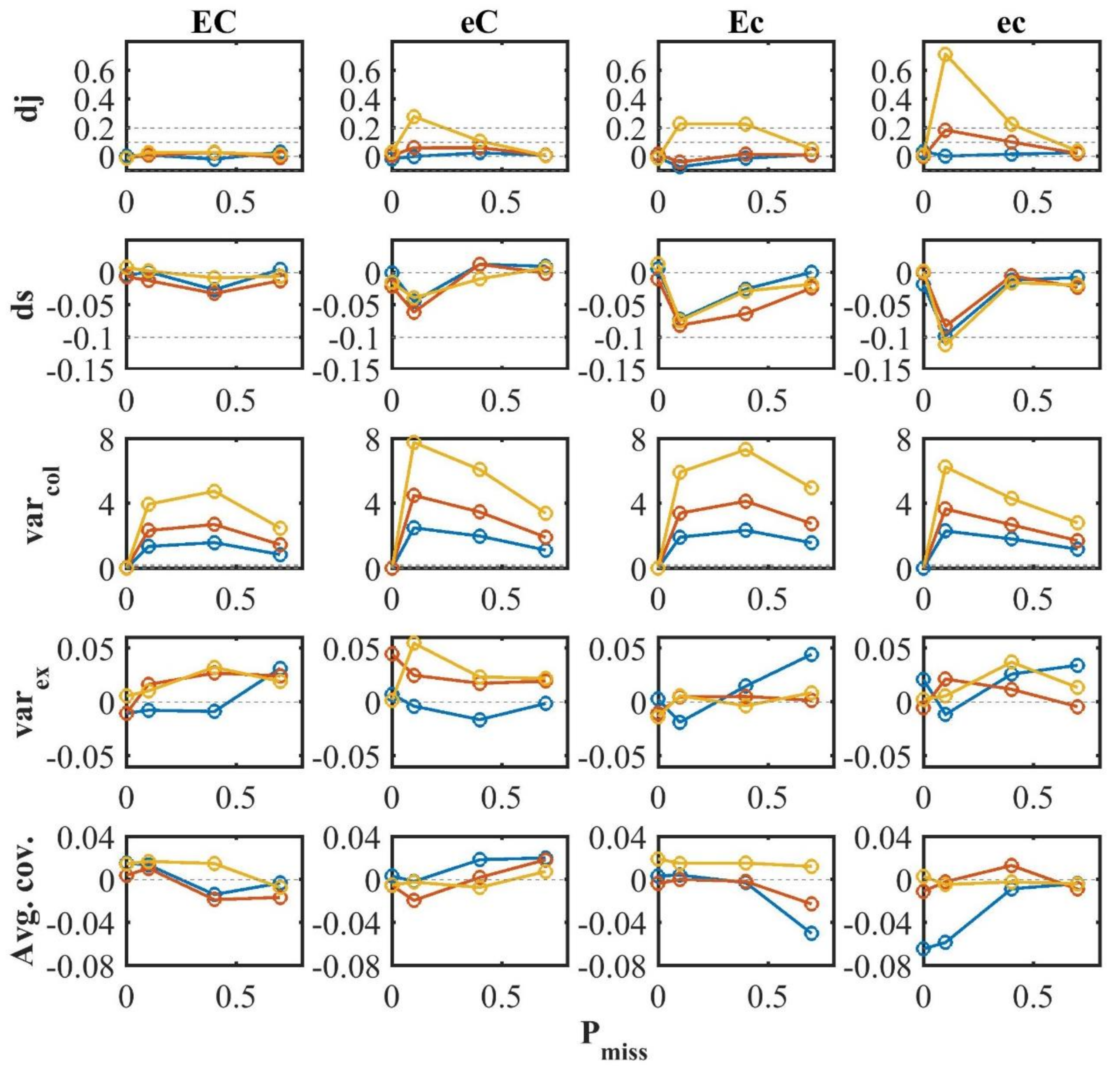


**Figure S6** – Standardized effect size of the statistics under DE, when each species present in the community has probability $P_{miss}$ (x axis) not to be detected each year. Each row tests the performance of one statistic, while each column considers a different combination of rates (EC stands for high extinction and high colonization, eC stands for low extinction and high colonization and so forth). Different colors represent different levels of $S_{reg}$ (blue: $S_{reg}$ =30, red: $S_{reg}$ = 90, yellow: $S_{reg}$ = 270). Here only the scenario of negative correlation between colonization and extinction is presented. The standardized effect size is calculated as the average statistic over 5000 synthetic datasets divided by the STD. The grey horizontal lines highlight the values -0.1, 0, 0.1 and 0.2.

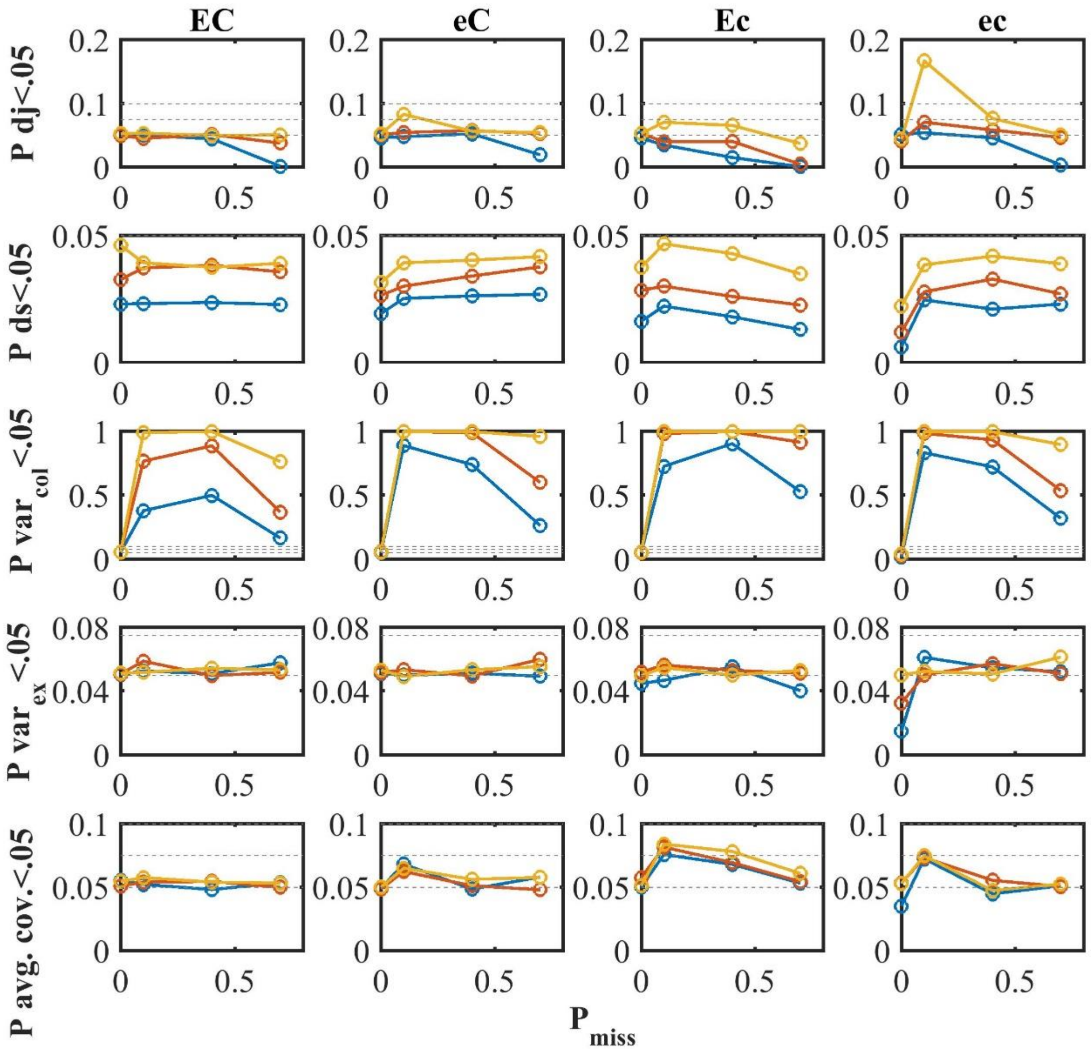


**Figure S7** – Significance of the statistics under DE, when each species present in the community has probability $P_{miss}$ (x axis) not to be detected each year. Each row shoes the performance of one statistic, presenting the proportion of significant results (out of the 5000 synthetic datasets), which is the effective α of the test. Each column considers a different combination of rates (EC stands for high extinction and high colonization, eC stands for low extinction and high colonization and so forth), while different colors represent different levels of $S_{reg}$ (blue: $S_{reg}$ =30, red: $S_{reg}$ = 90, yellow: $S_{reg}$ = 270). Here only the scenario of negative correlation between colonization and extinction is presented. The grey horizontal lines highlight the values of 0.05, 0.075 and 0.1.

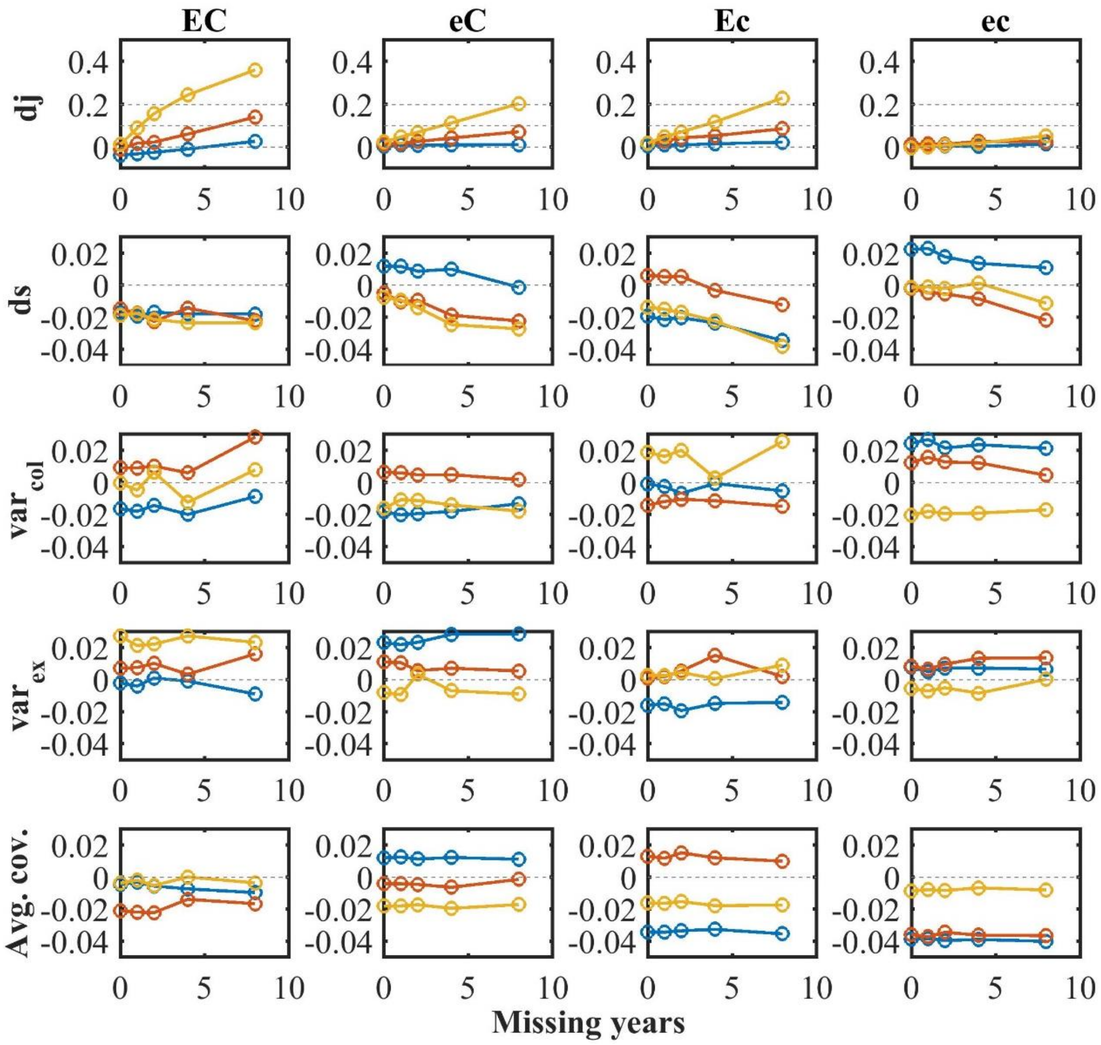


**Figure S8** – Standardized effect size of the statistics under DE, when the composition in a defined number of missing years (x axis) is transformed into missing values. Each row tests the performance of one statistic, while each column considers a different combination of rates (EC stands for high extinction and high colonization, eC stands for low extinction and high colonization and so forth). Different colors represent different levels of $S_{reg}$ (blue: $S_{reg}$ = 30, red: $S_{reg}$ = 90, yellow: $S_{reg}$ = 270). Here only the scenario of negative correlation between colonization and extinction is presented. The effect size is calculated as the average over 5000 synthetic datasets divided by the STD. The grey horizontal lines highlight the values -0.1, 0, 0.1 and 0.2.

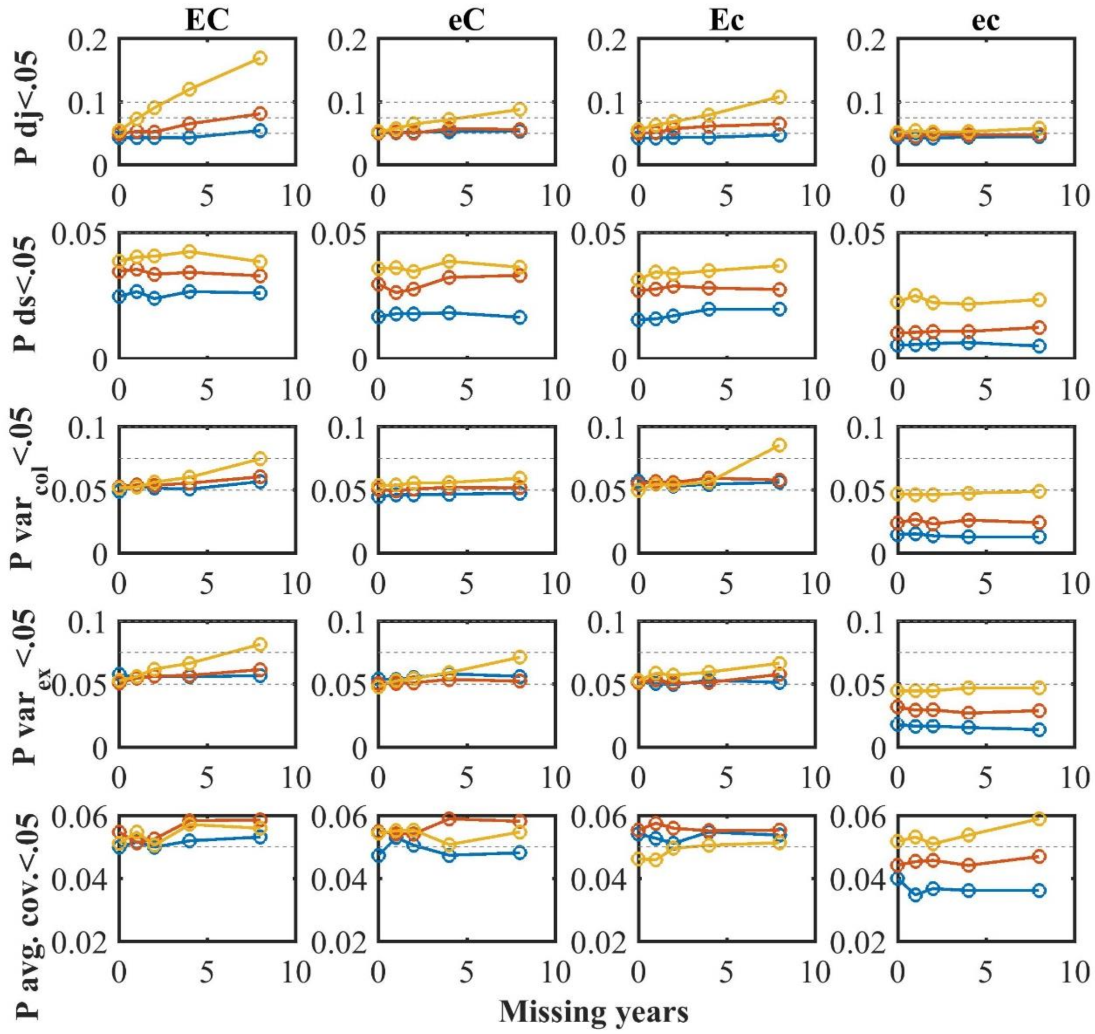


**Figure S9** – Significance of the statistics under DE, when the composition in a defined number of missing years (x axis) is transformed into missing values. Each row shoes the performance of one statistic, presenting the proportion of significant results (out of the 5000 synthetic datasets), which is the effective α of the test. Each column considers a different combination of rates (EC stands for high extinction and high colonization, eC stands for low extinction and high colonization and so forth), while different colors represent different levels of $S_{reg}$ (blue: $S_{reg}$ =30, red: $S_{reg}$ = 90, yellow: $S_{reg}$ = 270). Here only the scenario of negative correlation between colonization and extinction is presented. The grey horizontal lines highlight the values of 0.05, 0.075 and 0.1.

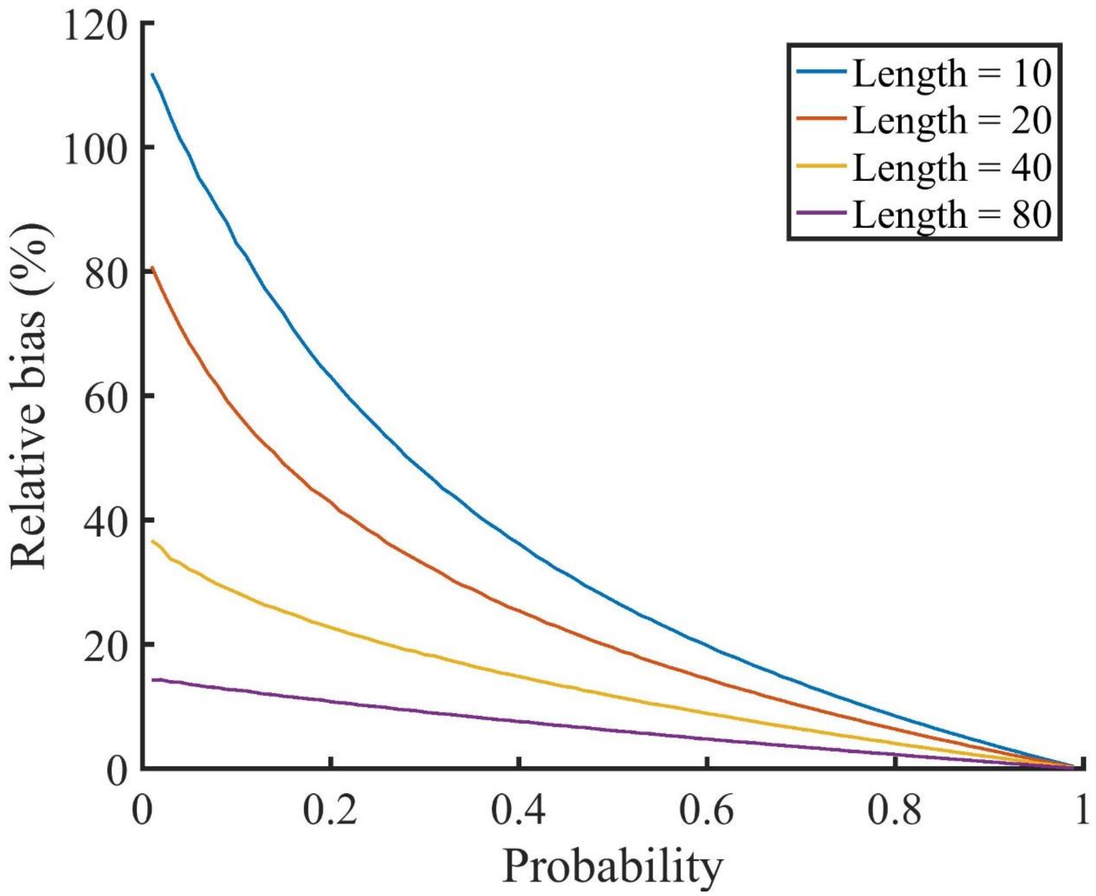


**Figure S10** – Relative bias in the maximum likelihood estimation of extinction probability (X axis) for different time series lengths. Since colonization and extinction are symmetric under DE, precisely the same bias would occur for colonization probabilities under analogous conditions. This analysis assumes that the species was initially present and that colonization probability is 0.1. $10^6$ time series were simulated for each extinction probability in the range 0.01 – 0.99 (in steps of 0.01) and then attempts were made to estimate this probability. We then calculated the relative bias, which is $(\hat{E} - E)/E$, where $E$ is the real extinction probability and $\hat{E}$ is the average estimate. Results given that the species is initially absent are qualitatively similar, but for short time series and high probability there is a negative bias ~ -20%.

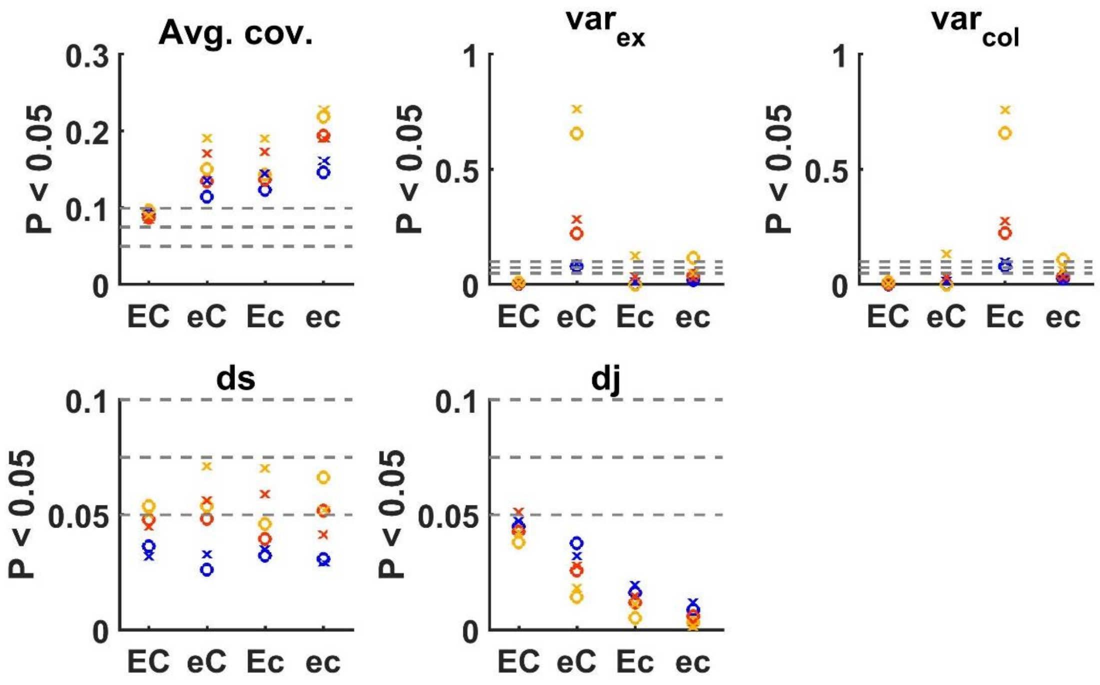


**Figure S11** – Proportion of rejected statistics under DE (type I errors) at various parameter regimes when the null is DE simulations with parameters estimated by maximum likelihood. Four combinations of rates are considered – high and low average colonization (C and c) and high and low average extinction (E and e). Simulations were run for each combination of rates, correlation between colonization and extinction ('o' = no correlation, 'x' = negative correlation) and size of the species pool (30 = blue, 90 = orange, 270 = yellow). Every point represents one of these combinations, and the proportion of significant results was calculated over 5000 synthetic communities. The grey horizontal lines highlight the values of 0.05, 0.075 and 0.1

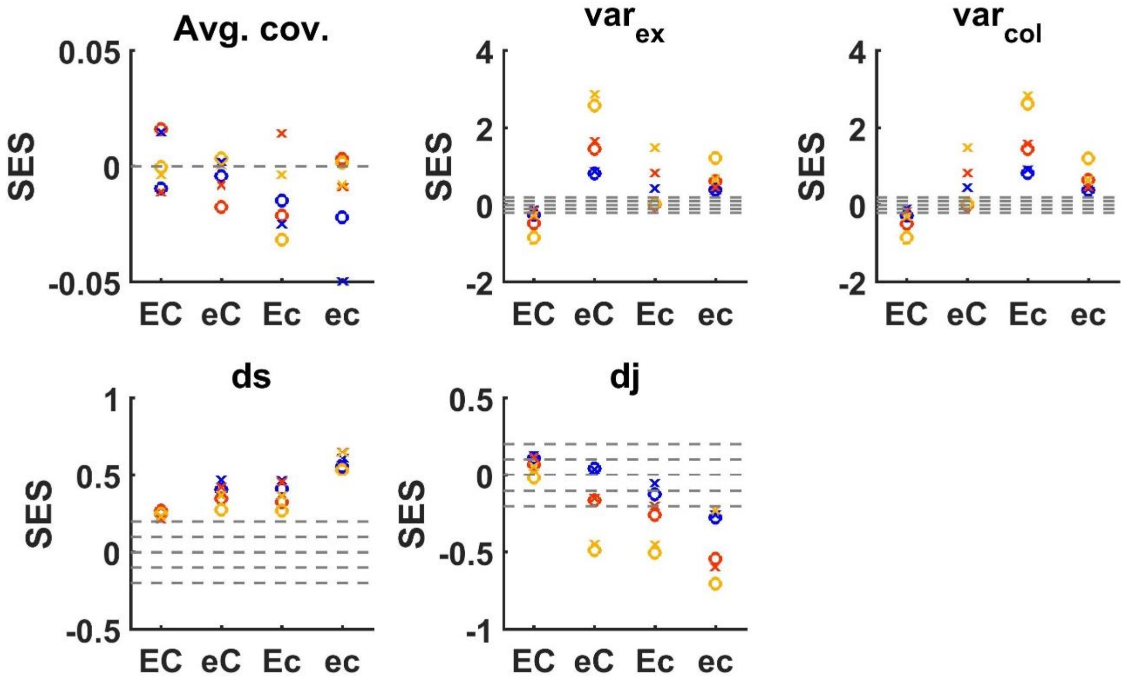


**Figure S12** – Standardized effect size of the statistics under DE at various parameter regimes when the null is DE simulations with parameters estimated by maximum likelihood. Four combinations of rates are considered – high and low average colonization (C and c) and high and low average extinction (E and e). Simulations were run for each combination of rates, correlation between colonization and extinction ('o' = no correlation, 'x' = negative correlation) and size of the species pool (30 = blue, 90 = orange, 270 = yellow). Every point represents one of these combinations, and the effect size is calculated as the average over 5000 synthetic datasets divided by the STD. The grey horizontal lines highlight the values -0.2, -0.1, 0, 0.1 and 0.2.

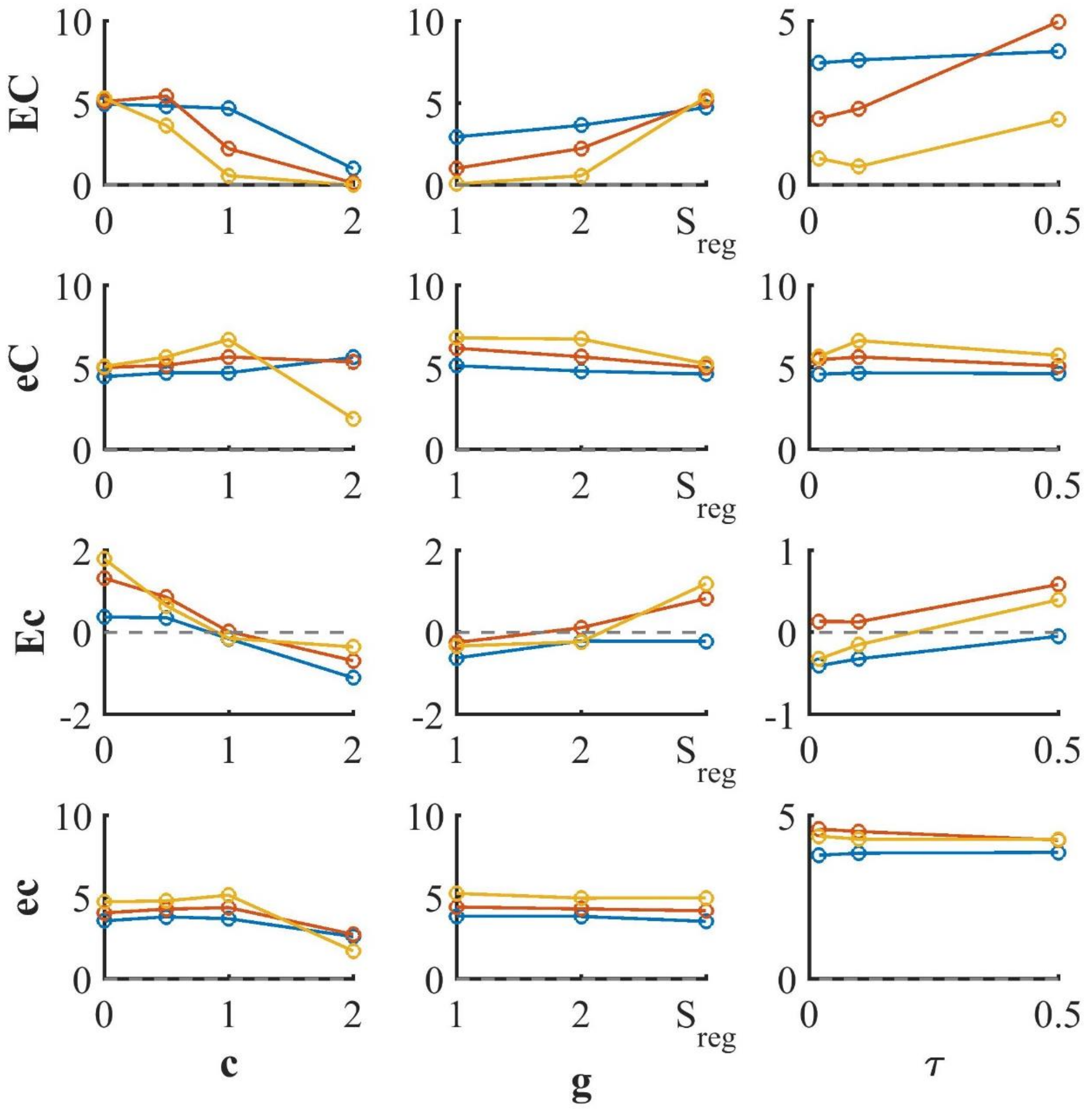


**Figure S13** – Natural logarithm of the ratio between the power of our methodology (*ds* statistic with PARIS null model) and the Variance to Mean Ratio (VMR) to detect deviations from DE under the alternative SDE model with a negative correlation between colonization and extinction rates. Four combinations of rates are considered – high and low average colonization (C and c) and high and low average extinction (E and e). Simulations were run for combinations of rates, parameters of the SDE model (c, g and τ), size of the species pool (30 = blue, 90 = orange, 270 = yellow) and under negative correlation between colonization and extinction. Each point represents 5000 simulations under a specified scenario, for which the proportion of significant *ds* values and VMR > 1 is recorded, and its log ratio is presented. The dashed grey line represents similar power, and above it our methodology is more powerful.

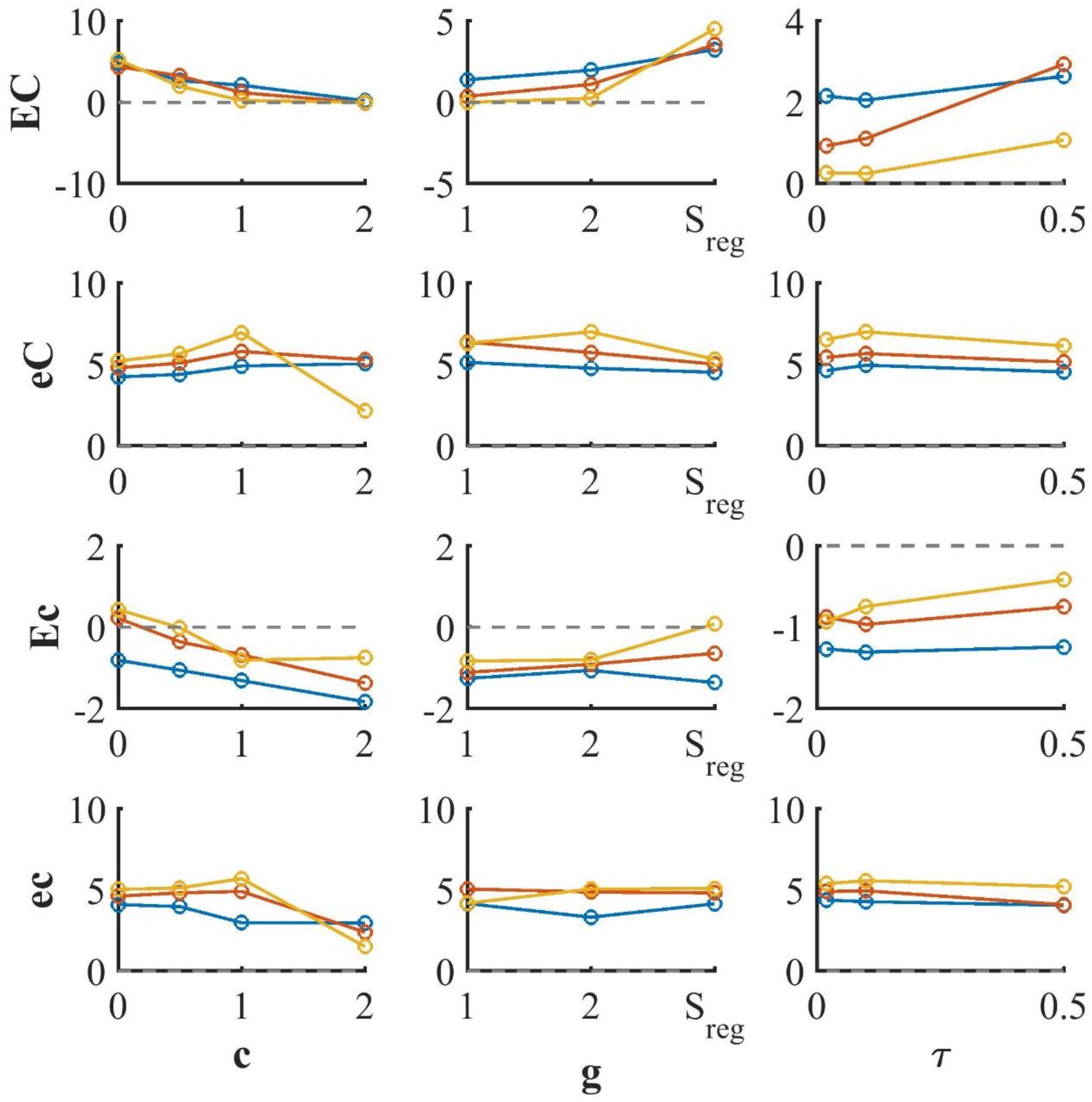


**Figure S14** – Natural logarithm of the ratio between the power of our methodology (*ds* statistic with PARIS null model) and the Variance to Mean Ratio (VMR) to detect deviations from DE under the alternative SDE model with no correlation between colonization and extinction rates. For generating the synthetic data, four combinations of rates are considered – high and low average colonization (C and c) and high and low average extinction (E and e). Simulations were run for combinations of rates, parameters of the SDE model (c, g and τ), size of the species pool (30 = blue, 90 = orange, 270 = yellow) and under negative correlation between colonization and extinction. Each point represents 5000 simulations under a specified scenario, for which the proportion of significant *ds* values and VMR > 1 is recorded, and its log ratio is presented. The dashed grey line represents similar power, and above it our methodology is more powerful.

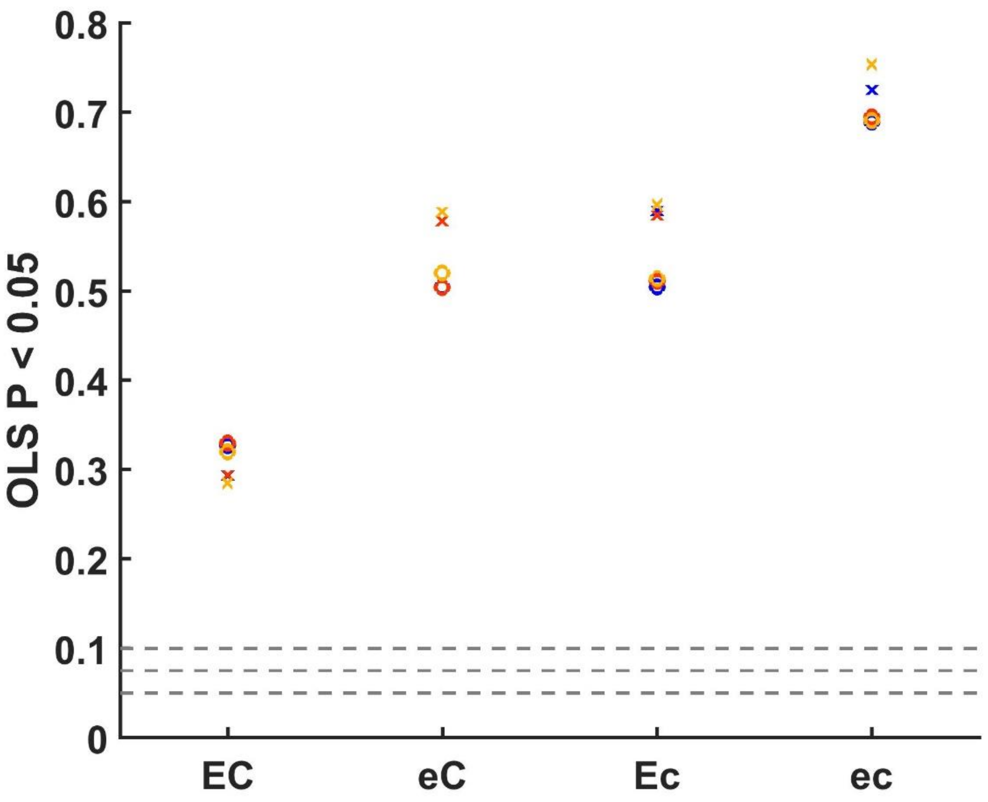


**Figure S15** – Proportion of rejected OLS regression slopes of richness vs. time under DE (type I errors) at various parameter regimes. For generating the synthetic data, four combinations of rates are considered – high and low average colonization (C and c) and high and low average extinction (E and e). Simulations were run for each combination of rates, correlation between colonization and extinction ('o' = no correlation, 'x' = negative correlation) and size of the species pool (30 = blue, 90 = orange, 270 = yellow). Every point represents one of these combinations, and the proportion of significant regression coefficients was calculated over 5000 synthetic communities. The grey horizontal lines highlight the values of 0.05, 0.075 and 0.1. The values for 30 species (blue) are sometimes identical to other values which obscure them.

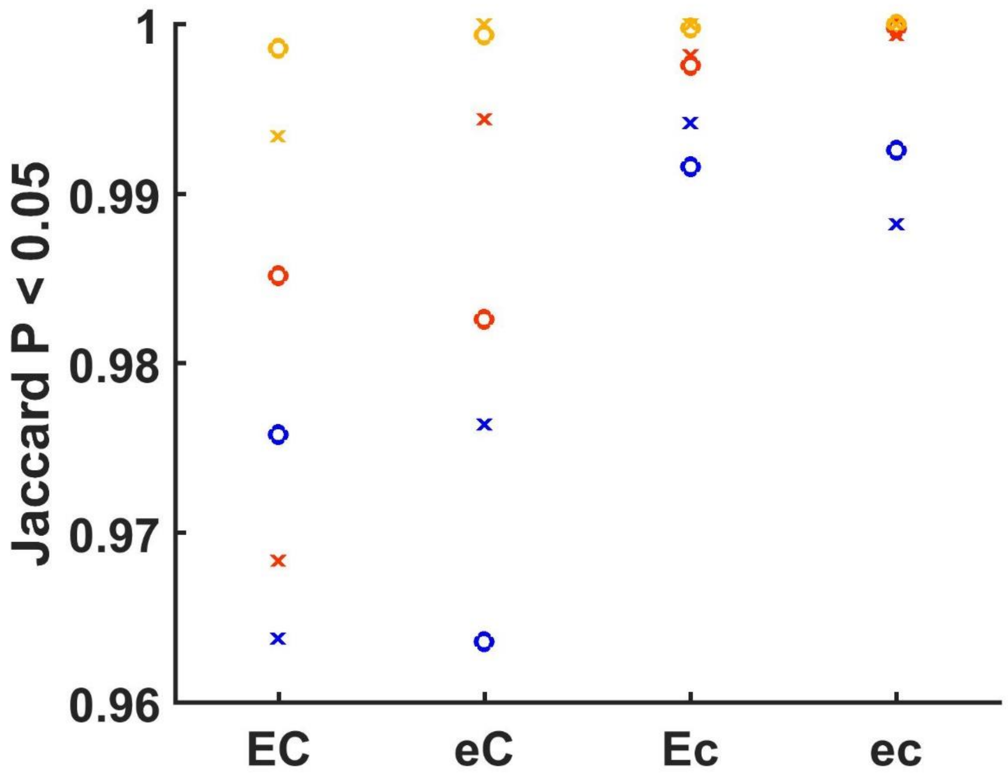


**Figure S16** – Proportion of rejected tests for compositional turnover using the method of (Dornelas *et al.* 2014)$) under DE (type I errors) at various parameter. For generating the synthetic data, four combinations of rates are considered – high and low average colonization (C and c) and high and low average extinction (E and e). Simulations were run for each combination of rates, correlation between colonization and extinction ('o' = no correlation, 'x' = negative correlation) and size of the species pool (30 = blue, 90 = orange, 270 = yellow). Every point represents one of these combinations, and the proportion of significant tests was calculated over 5000 synthetic communities.
